# Trihydroxybenzaldoximes are redox cycling inhibitors of ThDP-dependent DXP synthase

**DOI:** 10.1101/2025.03.03.641193

**Authors:** Charles R. Nosal, Ananya Majumdar, Netzahualcóyotl Arroyo-Currás, Caren L. Freel Meyers

## Abstract

Pathogenic bacteria must swiftly adapt to dynamic infection environments in order to survive and colonize in the host. 1-Deoxy-D-xylulose-5-phosphate synthase (DXPS) is thought to play a critical role in bacterial adaptation during infection and is a promising drug target. DXPS utilizes a thiamine diphosphate (ThDP) cofactor to catalyze the decarboxylative condensation of pyruvate and D-glyceraldehyde-3-phosphate (D-GAP) to form DXP, a precursor to isoprenoids and B vitamins. DXPS follows a ligand-gated mechanism in which pyruvate reacts with ThDP to form a long-lived lactyl-ThDP (LThDP) adduct which is coordinated by an active-site network of residues. D-GAP binding ostensibly disrupts this network to activate LThDP for decarboxylation. Our lab previously reported trihydroxybenzaldoximes inhibitors which are competitive with respect to D-GAP, and uncompetitive with respect to pyruvate, suggesting they bind after E-LThDP complex formation. Here, we conducted mechanistic studies to determine if these compounds inhibit DXPS by preventing LThDP activation or if they act as inducers of LThDP activation. We discovered that the catechol moiety of the trihydroxybenzaldoxime scaffold undergoes oxidation under alkaline aerobic conditions, and inhibitory potency is reduced under oxygen restriction. Leveraging long range ^1^H-^15^N HSQC NMR and electrochemical measurements, we demonstrated that the oxidized form of the trihydroxybenzaldoxime induces LThDP decarboxylation. The oxime moiety accepts electrons from the resulting carbanion, resulting in formation of acetyl-ThDP which hydrolyzes to form acetate. SAR studies revealed that the catechol attenuates the redox activity of the oxime moiety, and under aerobic conditions these compounds are oxidized and thus act as redox cycling inhibitors of DXPS. Further exploration of redox active DXPS probes may provide new insights for inhibition strategies and selective probe development.

## Introduction

Due to antibiotic resistance, infections which were previously treatable have become challenging to combat, and this resistance is predicted to contribute to more than 39 million deaths by 2050.^1^ Development of novel antibacterial strategies is required to combat the rapid evolution of bacterial resistance mechanisms. One approach is to investigate novel and underexploited therapeutic targets, such as processes in bacterial central metabolism.^2–6^ 1-Deoxy-D-xylulose-5-phosphate synthase (DXPS) is an essential central metabolic enzyme found in plants, bacteria and apicomplexan parasites.^7–12^ DXPS catalyzes synthesis of the branchpoint metabolite 1-deoxy-D-xylulose-5-phosphate (DXP) from pyruvate and D-glyceraldehyde-3-phosphate (D-GAP) utilizing thiamin diphosphate (ThDP) as cofactor (Fig 1). DXP serves as a precursor for isoprenoid building blocks isopentenyl diphosphate and dimethylallyl diphosphate as well as the biosynthesis of B vitamins, ThDP and pyridoxal phosphate (PLP).

**Figure 1.**
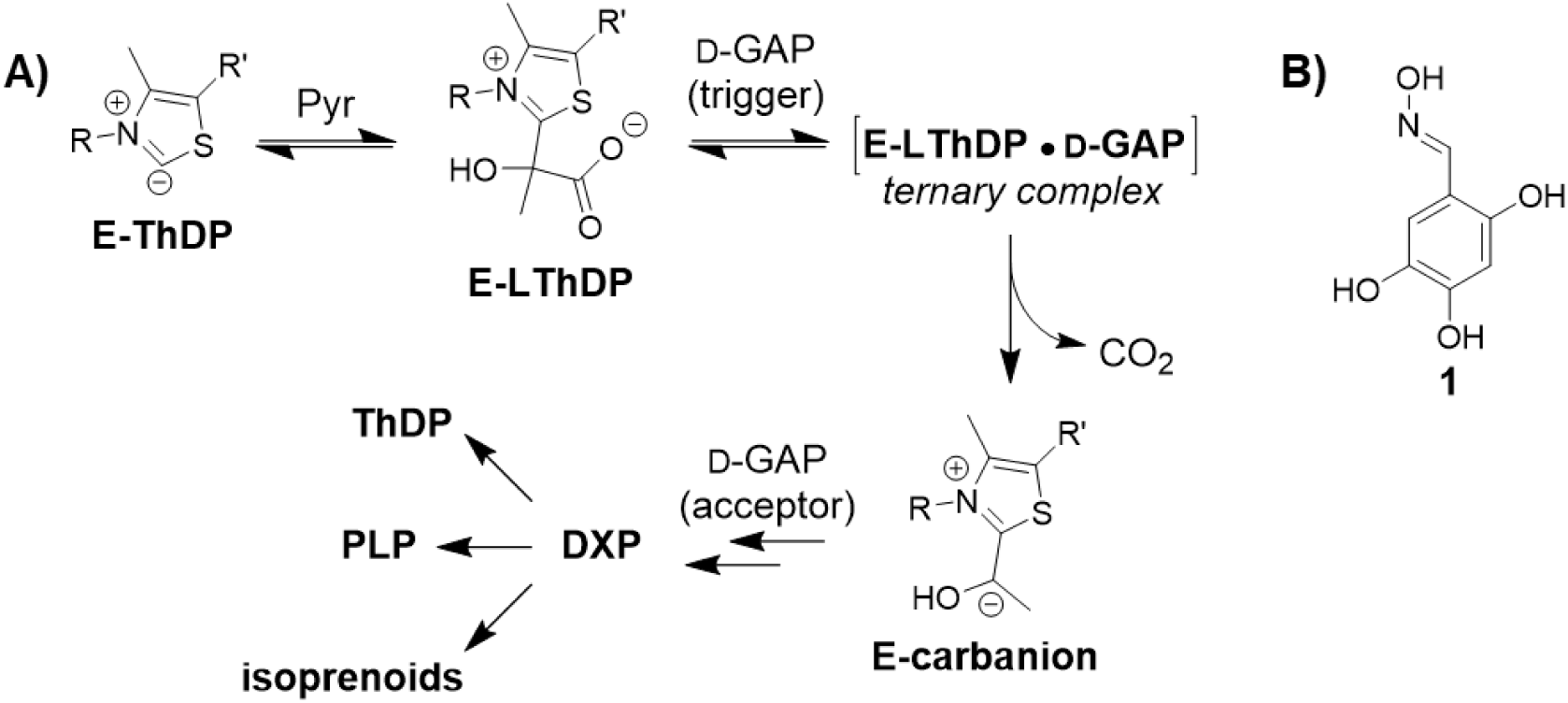
**A)** Mechanism of DXPS and its position at a branchpoint in bacterial central metabolism; **B)** Structure of trihydroxybenzaldoxime **1**.

Unlike other ThDP-dependent pyruvate decarboxylating enzymes, which generally follow ping-pong kinetics,^13–15^ DXPS utilizes a ligand-gated mechanism. On DXPS, pyruvate is activated to form lactyl-ThDP (LThDP), forming a stable E-LThDP complex in which an active site network of residues contributes to LThDP persistence in the absence of D-GAP.^16–27^ Upon GAP binding, a ternary complex is formed leading to activation of LThDP for decarboxylation and formation of the carbanion. Carboligation of the carbanion to D-GAP results in formation of DXP. Thus, D-GAP serves two roles, as an inducer or “trigger” of LThDP decarboxylation presumably via disruption of the coordinating active site network, and subsequently as an acceptor substrate in the carboligation step en route to DXP.^17,19,27,28^ D-GAP is not the only acceptor tolerated by DXPS, and DXPS can process other aldehydes,^24,29^ aromatic nitroso compounds,^30^ and even O_2_ via an electron transfer mechanism resulting in peracetate formation.^31^ In addition, other aldehydes and O_2_ act as inducers of LThDP decarboxylation.^24,31^ The substrate and catalytic promiscuity of DXPS suggests it could be inhibited by other unstudied triggers and/or acceptors.

We previously reported trihydroxybenzaldoximes that exhibited uncompetitive or noncompetitive modes of inhibition (MOI) relative to pyruvate and were competitive relative to D-GAP,^32^ with 2,4,5-trihydroxybenzaldoxime (**1**, Fig 1B) emerging as one of the most potent analogs. The uncompetitive mode of inhibition of **1** suggests the inhibitor binds after pyruvate and formation of the LThDP complex. Here, we have conducted mechanistic studies of this interesting mode of inhibition, toward our longer terms goal to develop selective, functional probes of DXPS and DXPS-targeting antibiotics. Our results indicate that **1** is selective for DXPS over a ThDP-dependent enzyme which does not use a gated mechanism. Altering the D-GAP binding pocket does not affect the potency of **1**, while substitution of active site network residue *Ec*R99^16^ leads to reduced potency, suggesting **1** may bind near LThDP in the E-LThDP complex. Nonenzymatic oxidative decarboxylation of pyruvate occurs in the presence of **1** in its oxidized form, promoted under aerobic alkaline conditions. Further, we found that **1** is most potent in its oxidized form, contrary to our previous report. Oxime potency correlates with electrophilicity of the oxime moiety, and the reduction potential of oxime compounds was attenuated by the catechol moiety. In its oxidized form, **1** induces LThDP decarboxylation and accepts electrons from the resulting carbanion to give reduced **1** which, in turn, is recycled via autoxidation; this result explains how **1** can potently inhibit DXPS under the aerobic reducing conditions reported previously by our lab. Taken together, our results demonstrate that trihydroxybenzaldoxime inhibitors act as selective redox cycling inhibitors of DXPS, revealing a new DXPS activity and inducer of LThDP decarboxylation. This study is significant as it provides insights that could be leveraged in the future development of potent and selective redox-active probes and antimicrobials.

## Results

### Study design

In contrast to competitive inhibition, in which the inhibitor may be outcompeted by accumulating substrate,^33^ the uncompetitive mode of inhibition requires the formation of enzyme-substrate complex. Uncompetitive inhibitors are highly sought after as they block enzyme activity in the presence of high substrate levels causing detrimental effects on metabolic flux. Thus, we are interested in understanding the selectivity and mechanism of uncompetitive inhibition by trihydroxybenzaldoximes such as **1**, toward a goal to design more potent, selective inhibitors of DXPS that act by this coveted mechanism. Our study explores the selectivity of **1** for DXPS over pyruvate decarboxylase (PDC) and conducts a preliminary mutagenesis study toward identifying the binding site for uncompetitive inhibition. We also conduct studies considering three mechanistic models (Fig 2) to determine which step downstream of LThDP formation is inhibited by **1**. In Model 1, **1** stabilizes LThDP and prevents decarboxylation. In Model 2, **1** induces the decarboxylation of LThDP, and the resulting E-carbanion is either stabilized by **1** [E-carbanion • **1**] or redirected into an unproductive substrate wasting pathway. Finally, in Model 3, **1** binds to E-carbanion following LThDP activation by another trigger molecule (e.g., D-GAP or O_2_).

**Figure 2.**
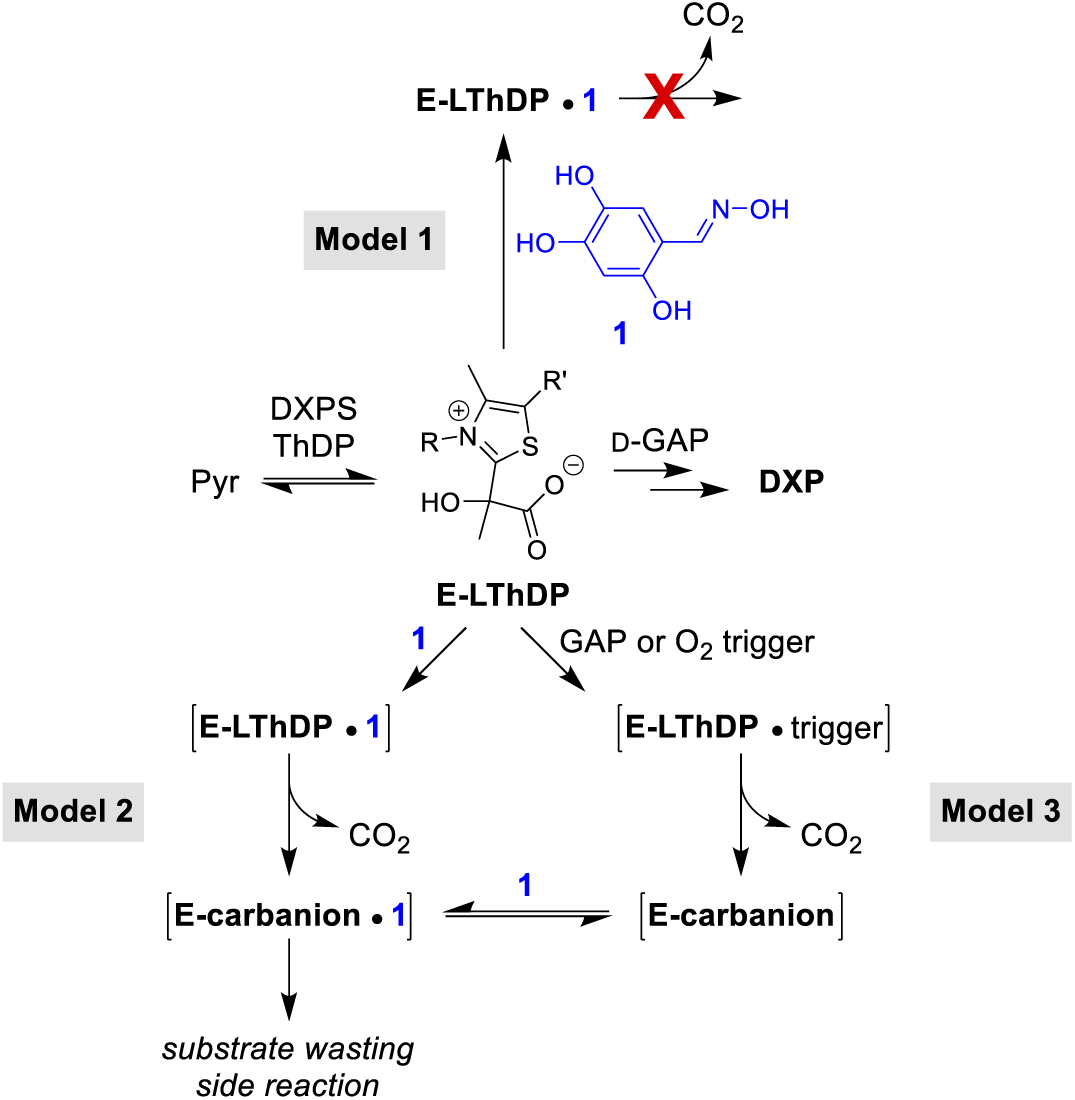
Mechanistic models for inhibition of DXPS by **1**.

### Oxime 1 is selective for DXPS

The uncompetitive mode of inhibition by **1** suggests it binds to the enzyme after formation of the stable E-LThDP complex, a feature which is unique to DXPS. We reasoned that **1** would display comparable potency against other DXPS homologs sharing this feature but may lack potency against pyruvate decarboxylase enzymes that do not use ligand gating. Thus, we assessed the potency of **1** against two other DXPS homologs relative to *Ec*DXPS, as well as pyruvate decarboxylase (PDC) which does not follow a ligand-gated mechanism. PDC was selected over PDH for this study since conditions to assay PDH contain cysteine, a thiol which have previously shown to react with this oxime class.^32,34,35^ DXPS homologs tested included *Deinoccocus radiodurans* DXPS (*Dr*DXPS) used for structural studies,^22,36^ and *Pseudomonas aeruginosa* DXPS (*Pa*DXPS), a pathogenic homolog that displays a stable E-LThDP complex, but differences in susceptibility to bisubstrate analog inhibitors relative to *Ec*DXPS.^37^ The *K*_i_ values for **1** against *Dr*DXPS when varying either pyruvate or D-GAP are comparable to *Ec*DXPS (within 2-fold) (Fig 3, S1, Table S1). A modest 3.5-fold reduction in potency was observed for **1** against *Pa*DXPS when varying pyruvate (Fig 3, S1, Table S1). Notably, **1** displayed uncompetitive or competitive modes of inhibition with respect to pyruvate or D-GAP, respectively (Fig S1), similar to its MOI against *Ec*DXPS and confirming its comparable behavior on three DXPS homologs.

**Figure 3.**
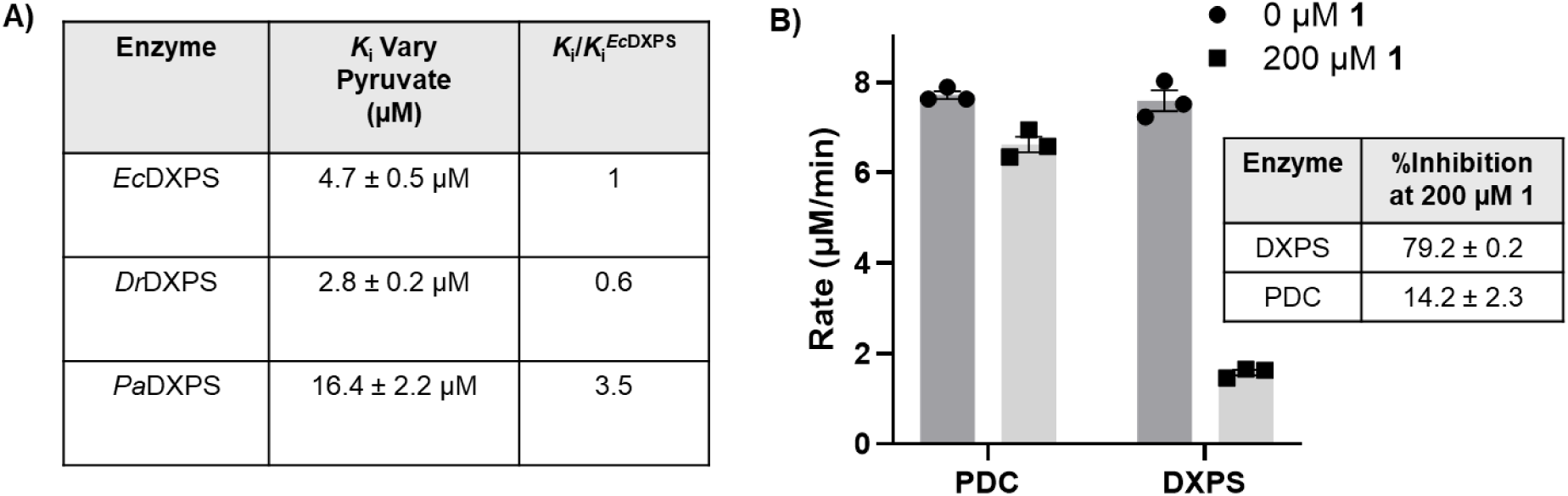
**A)** Summary of the activity of **1** against DXPS homologs. Under conditions of varying pyruvate (0-500 µM), **1** displayed an uncompetitive mode of inhibition against all homologs. **B)** Comparison of inhibitory activity of **1** (200 µM) compared to the no inhibitor control (0 μM **1**) against PDC (MES pH 6.2), relative to WT DXPS (HEPES pH 8), n=3. Standard error of the mean of reported % inhibition calculated from 3 replicates.

Evaluation of inhibitory activity of **1** against PDC from *S. cerevisiae* (Fig 3) was conducted using an alcohol dehydrogenase (ADH) coupled assay.^38,39^ We observed minimal inhibition of PDC (∼14%) in the presence of **1** at 200 μM, the highest concentration of **1** that does not interfere significantly with absorbance measurements in the coupled assay. In contrast, ∼79% inhibition of *Ec*DXPS activity was observed under comparable conditions. Oxime **1** did not inhibit ADH at this concentration (Fig S2). Since PDC activity is optimal at lower pH (∼pH 6)^38,40^ relative to DXPS (pH 8), **1** was tested against PDC at higher pH to determine if the potency of **1** depends upon pH. Inhibition of PDC by **1** at pH 7 or 8 was not observed (Fig S2). Thus, **1** is selective for DXPS over PDC, corroborating the selectivity of **1** for the ligand-gated DXPS mechanism.

### Investigating the enzymatic determinants of inhibition by 1 on *Ec*DXPS

To better understand how oxime **1** may interact with DXPS and the E-LThDP complex, we investigated a series of DXPS residues known to contribute to D-GAP binding and LThDP stability. As **1** inhibits DXPS competitively with respect to D-GAP, we reasoned that D-GAP and **1** share at least some portion of the D-GAP binding site (Fig 4A). Active site residues previously identified as being important for D-GAP binding include *Ec*R478 and *Ec*R420.^27^ Due to the extremely inefficient turnover of pyruvate and D-GAP on R420A DXPS,^27,41^ we investigated the potency of **1** against R478A to determine if R478 is involved in inhibitor binding. Similarly, the uncompetitive mode of inhibition requiring LThDP formation before binding suggests that **1** may bind at or near LThDP on the E-LThDP complex. Thus, **1** was also evaluated against active site network variants *Ec*D427A, *Ec*H431A, and *Ec*R99A, previously demonstrated to play a role in LThDP persistence on *Ec*DXPS (Fig 4A).^16^

**Figure 4.**
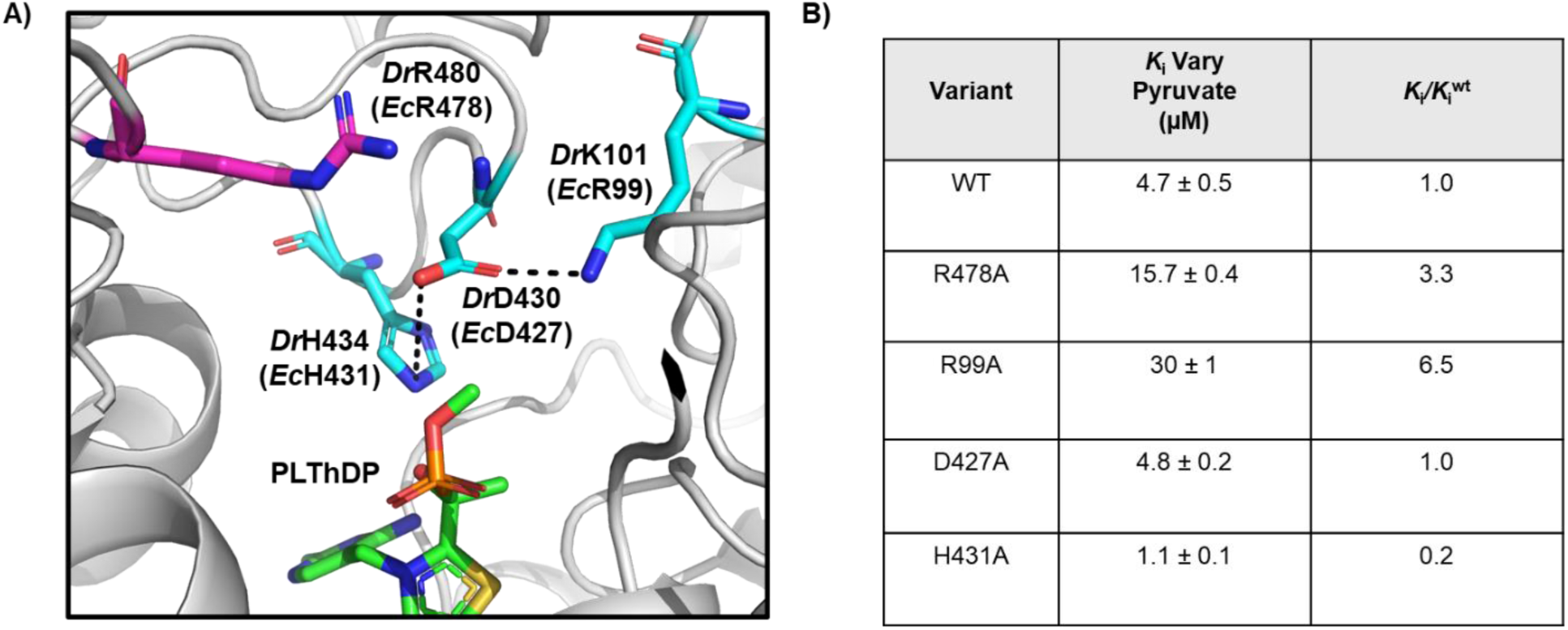
**A)** Active site of methyl acetyl phosphonate bound *Dr*DXPS (6OUV)^22^. PLThDP (green) is shown in proximity to coordinating active site network residues^16^ (cyan) *Dr*H434 (*Ec*H431A), *Dr*D430 (*Ec*D427), and *Dr*K101 (*Ec*R99). D-GAP binding residue^27^ (magenta) *Dr*R480 (*Ec*R478) is found in the active site in proximity to the network and PLThDP. **B)** Summary of potencies of **1** against active site network variants when varying pyruvate (0-500 µM). Oxime **1** displays an uncompetitive mode of inhibition against all variants under this condition. Error reported as standard error calculated from 3 replicates.

### R478A

To study the effect of the R478A substitution on the potency of **1**, the apparent *K*_i_ of **1** against the R478A variant was determined and compared to WT DXPS, using the DXPS-IspC coupled assay.^41,42^ In this experiment, pyruvate was varied from 12.5-500 µM and D-GAP was maintained at 2x *K*_m._. The potency of **1** against R478A DXPS (*K*_i_ = 15.7 ± 0.4 µM, Fig 4B, S3, Table S1) was modestly reduced (∼3-fold) relative to WT DXPS, and the uncompetitive mode of inhibition with respect to pyruvate was retained. In comparison, the DXPS bisubstrate inhibitor D-phenylalanine triazol acetyl phosphonate (D-PheTrAP)^43^ which interacts directly with the analogous residue *Dr*R480^36^, exhibits a > 30-fold loss in potency on *Ec*R478A.^37,43^ Thus, it appears that *Ec*R478A is not absolutely critical for binding **1**.

### R99A, D427A, and H431A

To determine the role of active site network residues in the activity of oximes, inhibitory activity of **1** was assessed against *Ec*D427A, *Ec*H431A, and *Ec*R99A by the DXPS-IspC coupled assay^41,42^ (Fig 4B, S3, Table S1). *K*_i_ determination was performed by varying [pyruvate] in order to assess the impact of residue substitutions on the uncompetitive mode of inhibition. Oxime **1** displayed an uncompetitive mode of inhibition on all three variants, with the *K*_i_ of **1** against D427A DXPS (4.8 ± 0.2 µM) comparable to WT *Ec*DXPS, and a ∼4-fold lower *K*_i_ of **1** against H431A DXPS (1.1 ± 0.1 µM) relative to WT. Interestingly, **1** showed a ∼6-fold reduction in potency against R99A DXPS (30 ± 1 µM) relative to WT DXPS. Further exploration revealed a lesser but measurable 4-fold reduction in potency of **1** on R99A DXPS relative to WT when varying [D-GAP] (8.7 ± 1.2 µM, Fig S3, Table S1). These data suggest that residue R99 plays a role in the activity of oxime **1** on DXPS. This result could be consistent with either Model 1 or 2 in which binding of **1** could serve to either strengthen (Model 1) or disrupt (Model 2) the active site network that R99A participates in and is responsible for stabilizing LThDP (Fig 2).

### Nonenzymatic conversion of pyruvate to acetate occurs in the presence of 1

To investigate whether **1** acts as an inducer of LThDP decarboxylation, we monitored the fate of ^13^C-labeled pyruvate after addition of **1** to pre-formed DXPS-LThDP, using 1D ^1^H-NMR and 1D ^1^H-NMR which selectively detects H{^13^C} and H_3_{^13^C} spectra via a proton multiplicity filter, as previously reported.^16,24^ Initial results revealed significant acetate formation in samples containing **1** (Fig S5), suggesting **1** may induce LThDP decarboxylation to give E-carbanion. E-carbanion could then undergo oxidation to acety-lThDP followed by hydrolysis to form acetate, as we have previously reported.^16^ However, a negative control experiment lacking DXPS also showed significant acetate formation from pyruvate in the presence of **1** (Fig S4). To identify the cause of this activity, reaction mixtures were generated in which components were removed with the exception **1**, pyruvate, and D_2_O in the presence or absence of gadobutrol (250 µM) which was used to amplify NMR signal. Under these conditions, acetate formation was not observed; however, upon addition of a small volume of HEPES pH 8 (∼3 mM final concentration), the solutions changed to a light pink color, and further NMR analysis revealed the evolution of acetate (Fig S5). This activity was ultimately determined to be O_2_-dependent, as acetate formation was hindered in anaerobic reactions (Fig 5, S6-7). Notably, shifts in the ^1^H-NMR peaks corresponding to **1** were also observed under conditions in which nonenzymatic decarboxylation of pyruvate was observed (Fig 5, HEPES pH 8, aerobic). The autoxidation of catechols has been previously reported to occur at basic pH^44–49^ with concomitant generation of reactive oxygen species (ROS)^47–50^ which are capable of reacting with pyruvate to form bicarbonate and acetate (Fig S8),^51,52^ consistent with what we observed by NMR. Thus, oxidation of **1** and associated pyruvate consumption were investigated in several buffer systems between pH 6.2 and pH 9 to assess trends in buffer and pH effects on catechol-dependent pyruvate oxidation.

**Figure 5.**
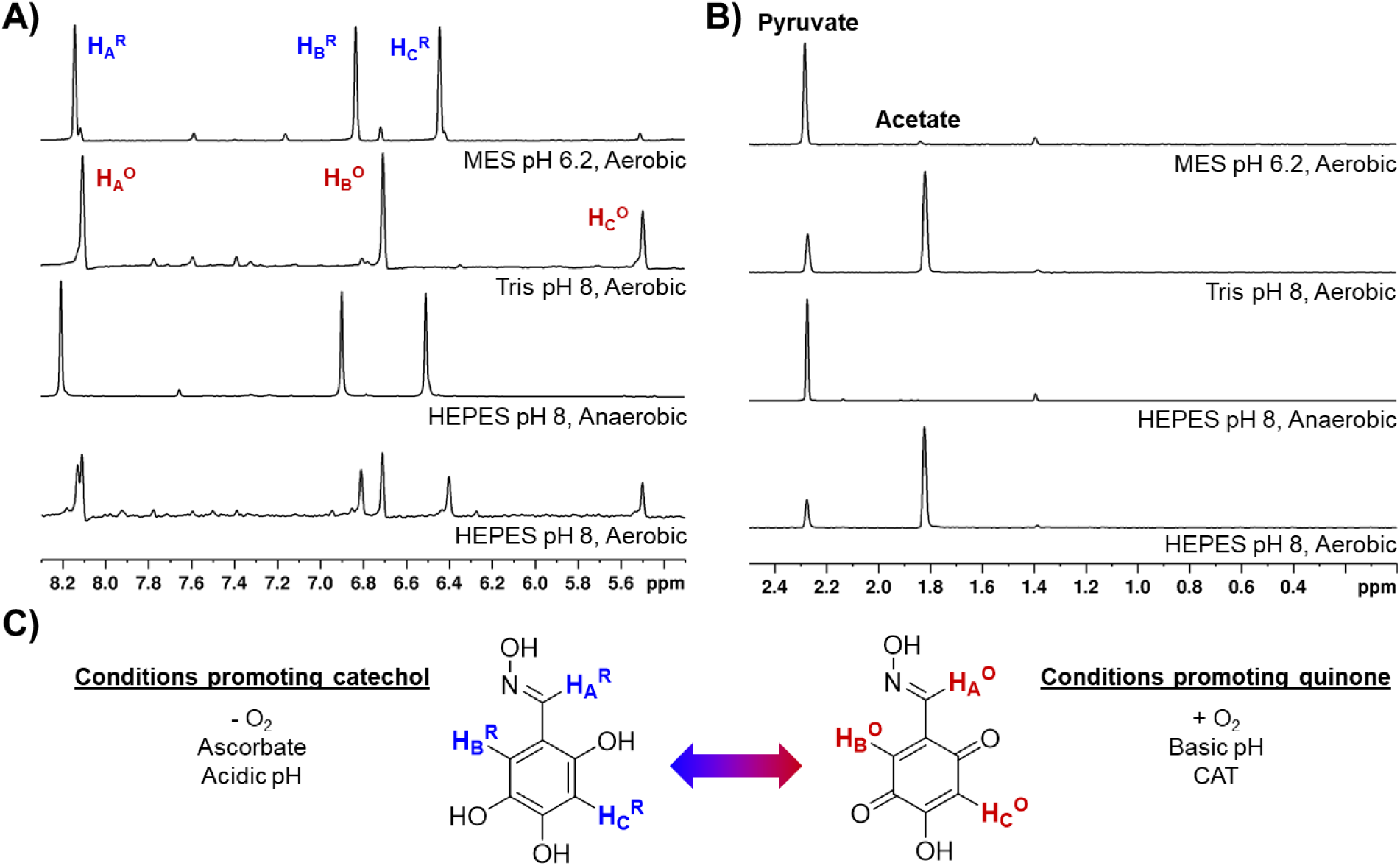
**A)** 1D ^1^H-NMR spectra of of **1** (500 µM) in varying buffers (50 mM buffer, 250 µM gadobutrol, Bruker Avance and Bruker Avance NEO 600 MHz). Oxime **1** appears to be oxidized under alkaline aerobic conditions and is reduced in acidic or anaerobic conditions. **B)** 1D ^1^H detected H{^13^C} and H_3_{^13^C} spectra via a proton multiplicity filter of ^13^C-labeled pyruvate and products (Bruker Avance and Bruker Avance NEO 600 MHz). Nonenzymatic consumption of pyruvate coincides with the oxidation of **1. C)** Comparison of redox forms of **1** and their associated conditions.

Our results indicated that autoxidation of **1** is pH-dependent, occurring readily under neutral-alkaline conditions, but not under acidic conditions (Fig 5, S6-7,9), consistent with previous reports.^46,49^ In addition, formation of acetate via decarboxylation of pyruvate was positively correlated with the oxidation of **1** (Fig 5, S6-7,9). Previously, we showed that ascorbate maintained **1** in its reduced form without appearing to affect potency.^32^ Thus, ascorbate (5 mM) was added to reaction mixtures containing **1** and pyruvate, in the presence of O_2_ (Fig S10), and conversion of pyruvate to acetate was assessed. The inclusion of excess ascorbate maintained **1** in its reduced form (Fig S10), but did not prevent pyruvate consumption and formation of acetate (Fig S10). This observation is in accordance with previous reports of ascorbate/quinone redox cycling activity^53^ and suggests that in the presence of O_2_, ascorbate promotes the redox cycle, reducing the quinone (or semiquinone) form of **1** after autoxidation of **1** in its catechol form and generation of ROS (Fig S11).

To generate further evidence of the oxidative decarboxylation of pyruvate by ROS, we added the H_2_O_2_ scrubbing enzyme catalase (CAT) to remove ROS generated via autoxidation of **1** (Fig S8,12).^54^ An excess of CAT (0.25 mg/mL or 500-1250 U/mL) was added since superoxide generated by autoxidation has been shown to inhibit CAT activity (Fig S6).^55,56^ Addition of CAT prevented oxidative decarboxylation of pyruvate under alkaline aerobic conditions. Notably, the oxidized form of **1** was predominant in the presence of CAT (Fig S12). These results support a role of ROS in the nonenzymatic oxidative decarboxylation of pyruvate and raise questions about how the oxidation state of **1** may impact its DXPS inhibitory activity.

### The redox state of 1 influences inhibitory activity

Given the impact of assay conditions on the oxidation state of oxime **1**, we studied the effect of oxidation state on inhibitor potency. As noted, we previously reported that **1** is active in its reduced form,^32^ supported by the observation that **1** exists primarily in its reduced form by ^13^C-NMR analysis, and the observation that **1** inhibits DXPS in the presence of ascorbate. However, here we note that at the high concentrations (>10 mM) required for natural abundance ^13^C-NMR, **1** appears to exist primarily in the reduced form in HEPES buffer pH 8 (Fig S13), which is not relevant to enzyme assay conditions conducted using lower concentrations of **1**. Additionally, in previous enzyme inhibition experiments conducted in the presence of ascorbate,^32^ **1** was pre-incubated with pyruvate and D-GAP at 37 °C for 10 min. In light of our findings presented above, we reasoned that the inhibitory activity observed for **1** in our previous report was influenced by the nonenzymatic depletion of pyruvate; redox cycling of **1** facilitated by addition of ascorbate (Fig S11) likely promoted the generation of ROS, leading to oxidative decarboxylation of pyruvate.

Given that CAT prevents nonenzymatic oxidative decarboxylation of pyruvate (Fig S8, S12), but influences the redox state of **1**, we evaluated the inhibitory activity of **1** in the presence of CAT (Fig 6). Likewise, the inhibitory activity of **1** was evaluated in the presence of ascorbate to confirm our previous conclusion about the role of ascorbate in maintaining oxime potency by promoting redox cycling in the presence of O_2_ (Fig 6). In this experiment, **1** was incubated at 25°C for 5 min in the presence of DXPS, IspC, and either CAT or ascorbate prior to initiation by addition of pyruvate and D-GAP. Addition of CAT, which promotes the oxidized form of **1**, removes H_2_O_2_, and prevents oxidative decarboxylation of pyruvate, does not impact the potency of **1** under aerobic conditions (80% inhibition at 100 µM **1**, Fig 6). This outcome indicates that **1** in its oxidized form inhibits DXPS in the absence of ROS formation and nonenzymatic consumption of pyruvate. In contrast, the inclusion of ascorbate results in a significant loss in potency of **1** (23% inhibition at 100 µM **1**, Fig 6). In the presence of ascorbate, **1** is primarily in the reduced form according to NMR (Fig S10). This result supports our conclusion that preincubation of **1** with substrates and ascorbate influenced measured DXPS activity via substrate consumption and suggests that **1** is less potent in the reduced form.

**Figure 6.**
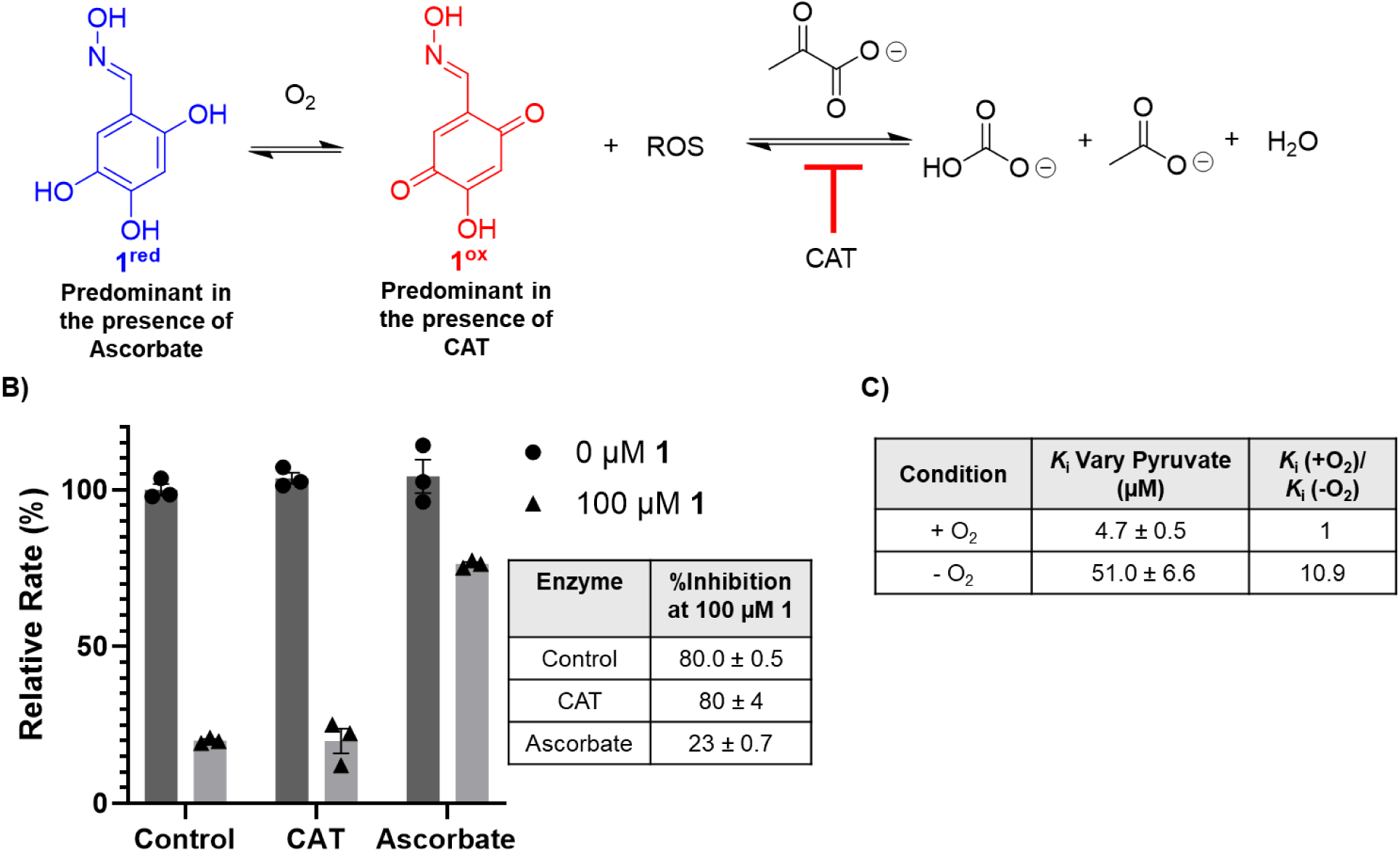
**A)** Autoxidation of **1** produces ROS that can react with pyruvate to form bicarbonate and acetate. **B)** Comparison of DXPS (150 nM) activity in the presence of ROS scrubbing enzymes at 0 and 100 µM **1**. X-axis labels indicate the inclusion of ROS scrubbing enzyme CAT (0.25 mg/mL) or non-nucleophilic reducing agent ascorbate (5 mM). Initial rates arithmetically corrected by subtracting background rates (no substrates). Standard error of the mean for reported % inhibition was calculated from three replicates. **C)** Summary of the anaerobic characterization of **1**. Data was fitted to an uncompetitive model (when varying pyruvate). Standard error of the mean for reported *K*_i_s was calculated from three replicates.

To learn how conditions which promote the reduced form of **1** in the absence of ROS impact inhibitor potency, we characterized the inhibitory activity of **1** under anaerobic conditions. In this case, **1** exists primarily in its reduced form (Fig 5, S6) and does not undergo autoxidation with concomitant ROS generation and nonenzymatic consumption of pyruvate.

Assays were performed in an anaerobic chamber after reagents were allowed to deoxygenate for ∼2 hours (Fig 6, S14). Under these conditions, the potency of **1** (*K*_i_ = 51.0 ± 6.6) decreased ∼10-fold relative to its potency under aerobic conditions (*K*_i_ = 4.7 ± 0.5), suggesting that **1** is a more potent uncompetitive inhibitor in its oxidized state. This finding contradicts our previous conclusion that **1** inhibits DXPS in its reduced form;^32^ however, as noted above, those studies were conducted under aerobic conditions in the presence of ascorbate where redox cycling of **1** and nonenzymatic pyruvate consumption were likely occurring.

### Inhibitory activity is attenuated by electronic properties of substituents

Previous work demonstrates that modifications can be made to the oxime and catechol moieties which alter potency and mode of inhibition.^32^ To better understand how the oxime moiety impacts the inhibitory activity of **1**, we compared the potencies of oximes bearing moieties with varied abilities to donate electron density to the oxime nitrogen, from least (**1**, oxime) to most electron-rich (**4**, *N-*methyl hydrazone) (Fig 7 S15, Table S2). We observed that analogs bearing electron-rich oxime substituents displayed reduced potency relative, with oxime analog **4** (*N-*methyl hydrazone) being the least potent, and analog **1** being the most potent. While **4** was previously reported as lacking measurable potency,^32^ this was determined in the presence of ascorbate which reduces the molecule, which we have demonstrated here to negatively impact potency. In fact, hydrazones **3** and **4** do exhibit measurable activity (*K*_i_ = 17.5 ± 0.6 µM and 56 ± 11 µM respectively) that is less potent than that of **1** or **2** (*K*_i_ = 4.7 ± 0.5 µM and 15.6 ± 0.7 µM^32^ respectively). Taken together, our observations suggest that electrophilicity of the nitrogen of **1** may be key for inhibitory activity.

**Figure 7.**
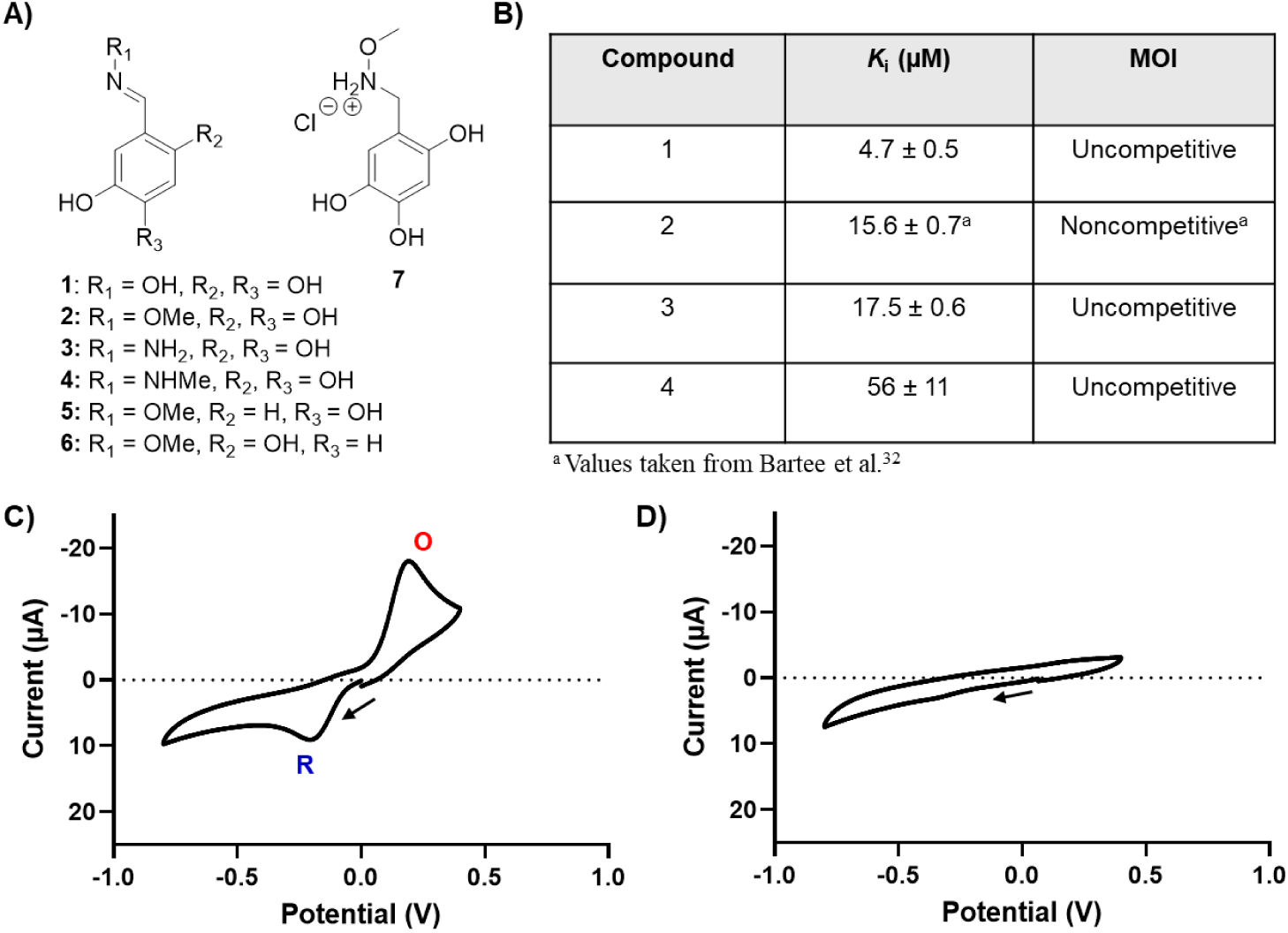
**A)** Structures of selected analogues of **1** investigated in this study. **B)** Table of potencies and MOI of oxime analogs when varying pyruvate. Initial rates arithmetically corrected for **4** by subtracting background rates (no substrates). Standard error of the mean of reported *K*_i_s was calculated from 3 replicates. **C)** CV of **1** and **D)** CV of **7** were collected by sweeping first in the negative direction. Oxidation (**O**), and reduction (**R**) are labeled for clarity. The start and direction of voltage scanning is indicated by a black arrow.

To further investigate the relevance of catechol oxidation to the potency of **1** and its derivatives, we utilized cyclic voltammetry to measure the electrochemical oxidation and reduction of a representative sample of catechol and oxime analogs. These analogs were selected for their varying activity and electronic properties of substituents. In these experiments compounds were prepared at 1 mM in phosphate-buffered saline (100 mM PO ^3-^ + 100 mM NaCl) pH 8 (Fig 7). Solutions were then deoxygenated via active bubbling with argon. Cyclic voltammograms (CVs) were measured using a three-electrode cell, consisting of a glassy carbon disk working electrode (5 mm diameter), a Ag|AgCl reference electrode, and a platinum counter electrode. First, a negative CV scan was performed to measure any reduction reactions, followed by a positive scan to probe for oxidations. Interestingly, **1** and **2**, which demonstrate similar potencies, appeared to undergo a reduction at around −0.2 V (Fig 7, S16). In contrast, inactive compounds **5**-**7**, did not show this reduction peak, though a small reduction peak was observed around −0.34 V for **6** (Fig S16). The large oxidation peaks observed in compounds **1** (∼0.2 V), **2** (∼0.18 V), **5** (∼0.35 V) and **6** (∼0.39 V) (Fig 7, S16) appeared to qualitatively correlate with the number of hydroxyl substituents on each oxime (three for **1**, **2** versus two for **5**, **6**) and this is presumed to be related to oxidation of the catechol.^57^ When the oxime moiety was reduced to methoxylamine (**7**), we observed a loss of reduction and oxidation peaks (Fig 7). Compounds **5** and **6** exhibited large reduction peaks at 0.24 and 0.14V respectively (Fig S16), which is counterintuitive given their lack of potency, as this suggests they are more favorable electron acceptors. However, these compounds lack one hydroxyl group. Overall, it appears that oximes with inhibitory activity show a reduction peak at ∼-0.2 V and require a trihydroxy catechol. It is likely that the trihydroxy substitution modulates the reduction potential of oximes **1** and **2** via inductive effects, an effect that may play an important role in the inhibitory mechanism of **1**. Our previous study showed that a trihydroxy catechol is not strictly required for activity, further corroborating that the role of the hydroxyl groups in this case is likely modulation of the oxime’s redox potential.^58^ These results taken together suggest that redox activity may play an important role in the mechanism of inhibition of DXPS by **1** in which redox potential must be carefully tuned for potent inhibitory activity.

### Oxime is reduced by E-carbanion

In the mechanism of DXPS, E-LThDP undergoes decarboxylation to form E-carbanion in the presence of a trigger molecule (Fig 1). Enzymes which catalyze redox reactions involving carbanions frequently rely on cofactors like ThDP, flavin mononucleotide, and NAD(P)H among others to facilitate one and two electron transfers.^59–61^ In our previous studies of DXPS variants that readily activate LThDP for decarboxylation in the absence of a trigger, we observed acetate production, presumably via oxidation of the carbanion to acetyl-ThDP followed by hydrolysis.^16^ We have also shown that the C2-α-carbanion transfers an electron to O_2_ to form a C2-(α-hydroxy)-ethylideneThDP radical with subsequent recombination with superoxide en route to peracetate.^31^ Thus, it is plausible that the C2-α-carbanion can reduce the oxidized form of **1**. Toward establishing this mechanism on DXPS, we prepared **1** using hydroxylamine enriched in ^15^N (**8**, Fig 8) and evaluated its reduced and oxidized forms as well as its reactivity in the presence of DXPS and substrates, using long range ^1^H-^15^N HSQC NMR. By this method, we readily distinguished the oxidized and reduced forms of **8** via ^2^J_HN_ (3 Hz) and ^4^J_HN_ (<1 Hz) couplings due to the significant difference in the chemical shift of the ^15^N peaks when **8** is reduced or oxidized (Fig 8A). The true ^15^N chemical shifts fall in the regions of δ355 ppm (N^R^) and δ374 ppm (N^O^) (Fig S17). In the 2D spectra acquired here, ^15^N spectral widths have been shortened to optimize data acquisition, resulting in the ^15^N peaks appearing at the appropriately “folded” positions indicated (Fig 8).

**Figure 8.**
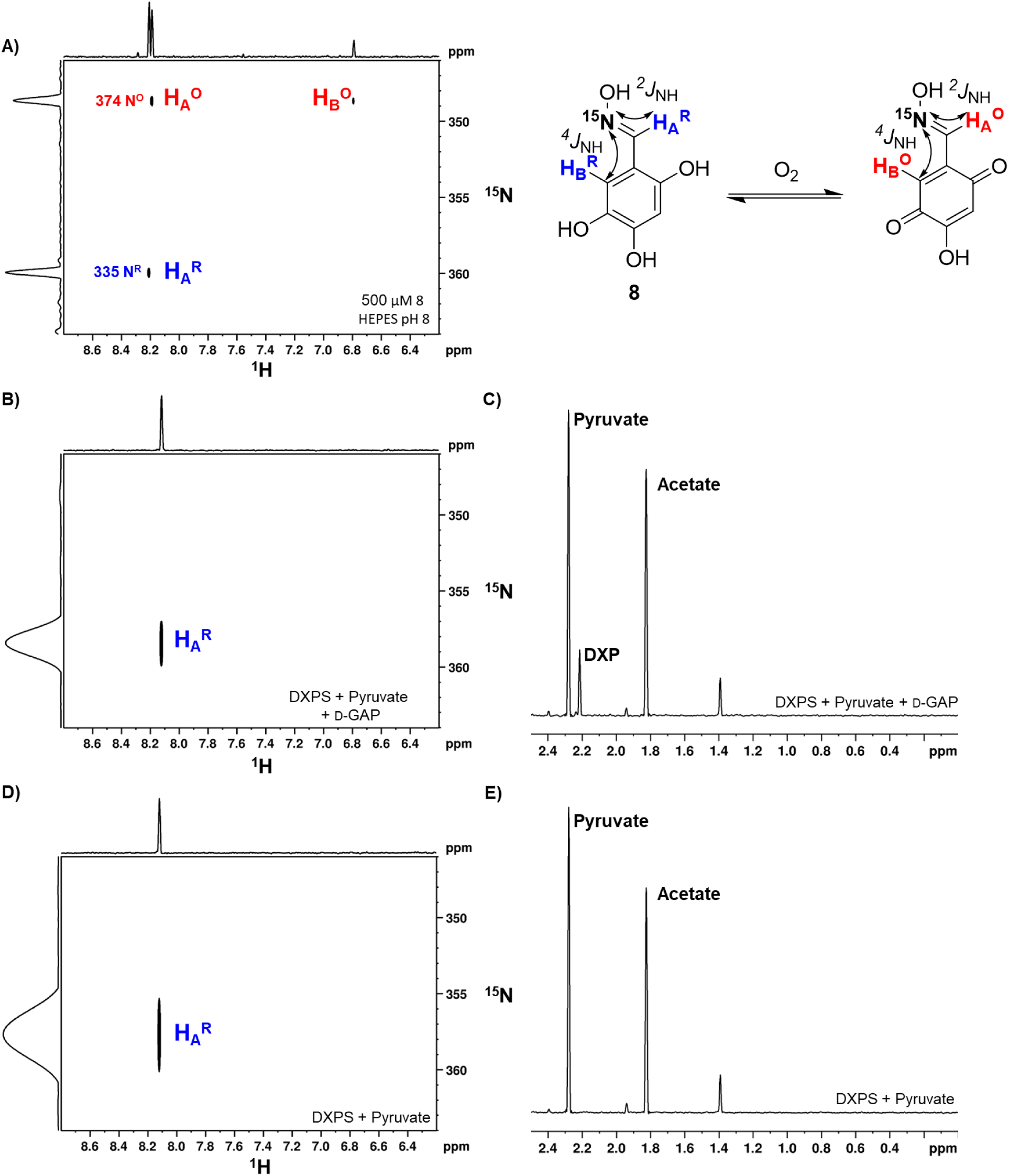
**A)** Aerobic preparation of **8** (500 µM) in HEPES (50 mM pH 8). Two and four-bond *J*_NH_ couplings were used to distinguish between oxidized and reduced forms. The spectral width along ^15^N has been reduced for optimized data acquisition, resulting in the folding of the N^R^ and N^O^ peaks along the ^15^N dimension. The true ^15^N chemical shifts (Fig S17) are labeled here for clarity. **B)** Long range **^1^**H-^15^N HSQC of **8** after incubation with DXPS (1 µM), pyruvate (2 mM), and D-GAP (500 µM). Oxime is reduced after reacting with DXPS and substrates. **C)** 1D ^1^H detected H{^13^C} and H_3_{^13^C} spectrum of **8** (via a proton multiplicity filter of ^13^C-labeled pyruvate and products) after incubation with DXPS (1 µM), pyruvate (2 mM), and D-GAP (500 µM). DXP and acetate formation are observed. **D) ^1^**H-^15^N HSQC of **8** after incubation with DXPS (1 µM) and pyruvate (2 mM). Oxime is reduced after reacting with DXPS and pyruvate without the presence of an external inducer of decarboxylation (e.g. D-GAP or O_2_). **E)** 1D ^1^H detected H{^13^C} and H_3_{^13^C} spectrum of **1** (via a proton multiplicity filter of ^13^C-labeled pyruvate and products) after incubation with DXPS (1 µM) and pyruvate (2 mM). Acetate is produced in the course of this reaction. Spectra collected on a Bruker Avance NEO 600 MHz spectrometer.

In order to determine if **1** is reduced by the C2-α-carbanion on DXPS, we prepared the ^15^N-labeled analog **8** aerobically in Tris pH 8 (to bias the oxidized form, Fig S7) and then transferred this mixture to the anaerobic chamber to minimize O_2_-mediated re-oxidation of **8** following reduction by the C2-α-carbanion (Fig S18). Control experiments containing ascorbate, DXPS in the absence of substrates, or pyruvate and D-GAP in the absence of DXPS, were conducted to demonstrate the capability of this technique to monitor oxidation/reduction of **8** (Fig S19-21). Reactions containing **8** (500 μM) were then initiated by the addition of DXPS (1 µM) and ^13^C-labeled pyruvate (2 mM), in the presence or absence of 500 µM GAP (Fig 8B-E, S22-23). Samples were then quenched by boiling and vortexing before being transferred into degassed NMR tubes and sealed with rubber septa and parafilm to preclude reoxygenation. As predicted, **8** was readily reduced in the presence of DXPS and substrates. In the presence of DXPS or absence of substrates, **8** was maintained in the oxidized form, while partial reversion to the reduced form was observed in the presence of substrates and absence of DXPS (Fig S20-21). Of note, **8** was fully reduced in the presence of DXPS and pyruvate, without GAP added as a trigger of LThDP decarboxylation, indicating that **8** is capable of inducing LThDP decarboxylation over multiple turnovers, consistent with Model 2 (Fig 2). Further evidence supporting Model 2 in which **1** induces LThDP decarboxylation and promotes substrate wasting, was obtained by detection of products using 1D ^1^H-NMR which selectively detects H{^13^C} and H_3_{^13^C} spectra via a proton multiplicity filter (Fig 8, S23-24). As expected, significant acetate production was observed in the presence of **8**, formed via hydrolysis of the acetyl-ThDP intermediate generated during reduction of **8** (Fig S25). Production of DXP relative to acetate in the presence of D-GAP was reduced in the presence of **8** (Fig S24), consistent with a mechanism in which **1** competes with GAP as an acceptor and in agreement with the previously characterized competitive mechanism of inhibition with respect to D-GAP.^32^ Some acetate was observed in reactions containing substrates in the absence of DXPS which could be attributed to the formation of ROS during the aerobic incubation period at the start of the procedure.

To provide additional evidence for DXPS-dependent reduction of **1**, we used electrochemistry to continuously monitor the redox state of **1** during the course of a DXPS reaction. For these experiments, we used a gold ultramicroelectrode, which allowed us to observe the steady-state current corresponding to the reduction of **1**, under diffusion control, during the progression of the reaction. A steady state CV of **1** in a reaction buffer containing Tris pH 8 prepared aerobically followed by deoxygenation by argon via active bubbling revealed a reduction with half-height potential of ∼-0.35 V (Fig 9). Using Equation 4 (Electrochemical Measurements, Materials and Methods), it was estimated that the observed pseudo steady state current at −0.50 V corresponded to a one-electron transfer reaction. We used this voltage to interrogate the reduction of **1** during reaction with enzyme and substrates. The idea is as follows: if **1** is chemically reduced in the bulk solution via DXPS, a drop in the steady state current should be observed as the population of **1** capable of undergoing electrochemical reduction at the electrode surface diminishes. Reactions were initiated in the cell via addition of different combinations of DXPS and substrates which were previously deoxygenated in an anaerobic chamber (Fig 9, S26). In accordance with the NMR results, a decrease in current was observed due to the dilution of **1** when DXPS or substrates are added individually to reaction mixtures, but the current stabilized and returned to a steady state. However, when DXPS and substrates were added together, a rapid and continuous decrease in current was observed until reaching a near zero rate. Also in accordance with NMR results, the drop in current observed when GAP was excluded was more rapid, reinforcing that **1** is capable of inducing decarboxylation on DXPS and competes with GAP for the resulting electrons. The reduction of **1** was also observed via the color of solution; the oxidized form of **1** turned the solution pink, whereas after reduction, the solution became yellow (Fig S27). Returning to the CV, when measured after the chemical reaction transpires, the reduction peak nearly vanished as expected (Fig 9, S28). Oddly, the oxidation peak that was observed did not appear to change concurrently with the reduction (Fig S28), corroborating the previous conclusion that the oxidation is related to the catechol and not tied to the reduction of the oxime moiety.

**Figure 9.**
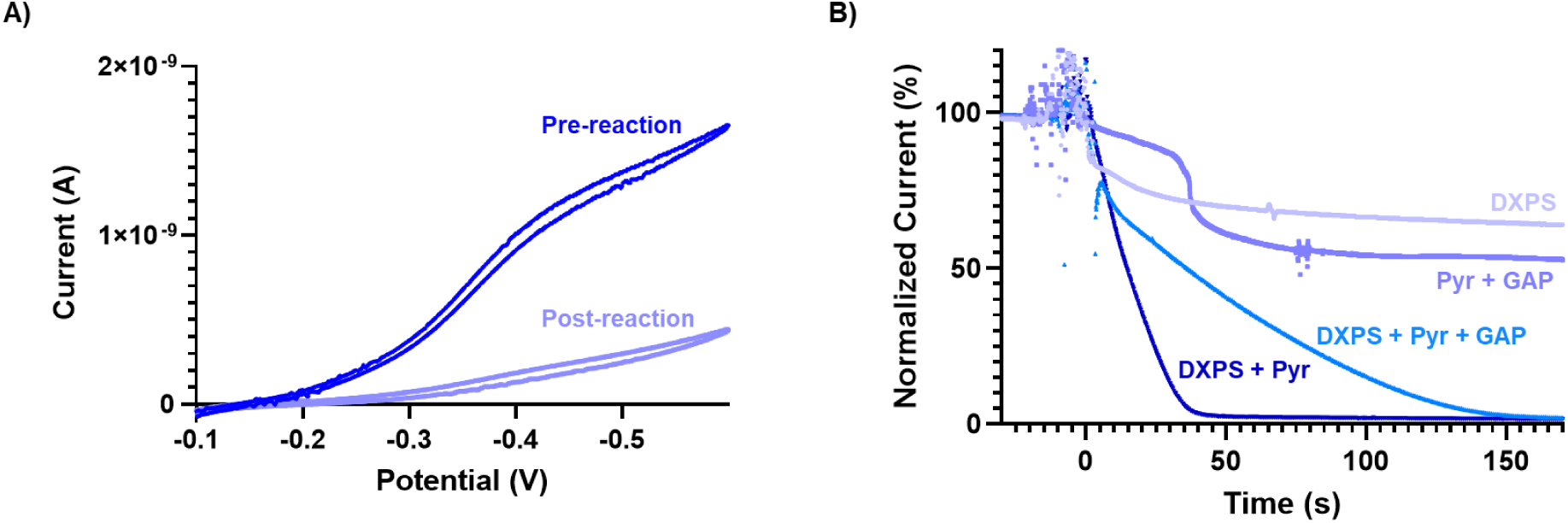
**A)** Anaerobic CV of **1** was conducted using a gold UME sweeping negative collected before and after reaction with DXPS (10 µM), pyruvate (2 mM), and D-GAP (500 µM). **B)** Normalized amperoteric i-t curves were measured at −0.5 V monitoring chemical reduction of **1** by loss in measured current. Reaction initiation was set to 0 seconds. Reactions contained either DXPS (10 µM), Pyruvate (2 mM) + D-GAP (500 µM), DXPS (10 µM) + Pyruvate (2 mM) + D-GAP (500 µM), or DXPS (10 µM) + Pyruvate (2 mM).

### Working model of inhibition by 1

Taken together, our results support a mechanism of inhibition consistent with Model 2 (Fig 2). The oxidized form of **1** (**1^ox^**, Fig 10), present at pH 8 under aerobic conditions (Fig 5), activates LThDP for decarboxylation. Subsequently, electron transfer occurs from the C2-α-carbanion to **1^ox^**, reducing **1** to the catechol (**1^red^**) and oxidizing the C2-α-carbanion to acetyl-ThDP. Hydrolysis of the acetyl-ThDP intermediate produces acetate as the end product of pyruvate, and under aerobic conditions, **1** is re-oxidized to complete the redox cycle.

**Figure 10.**
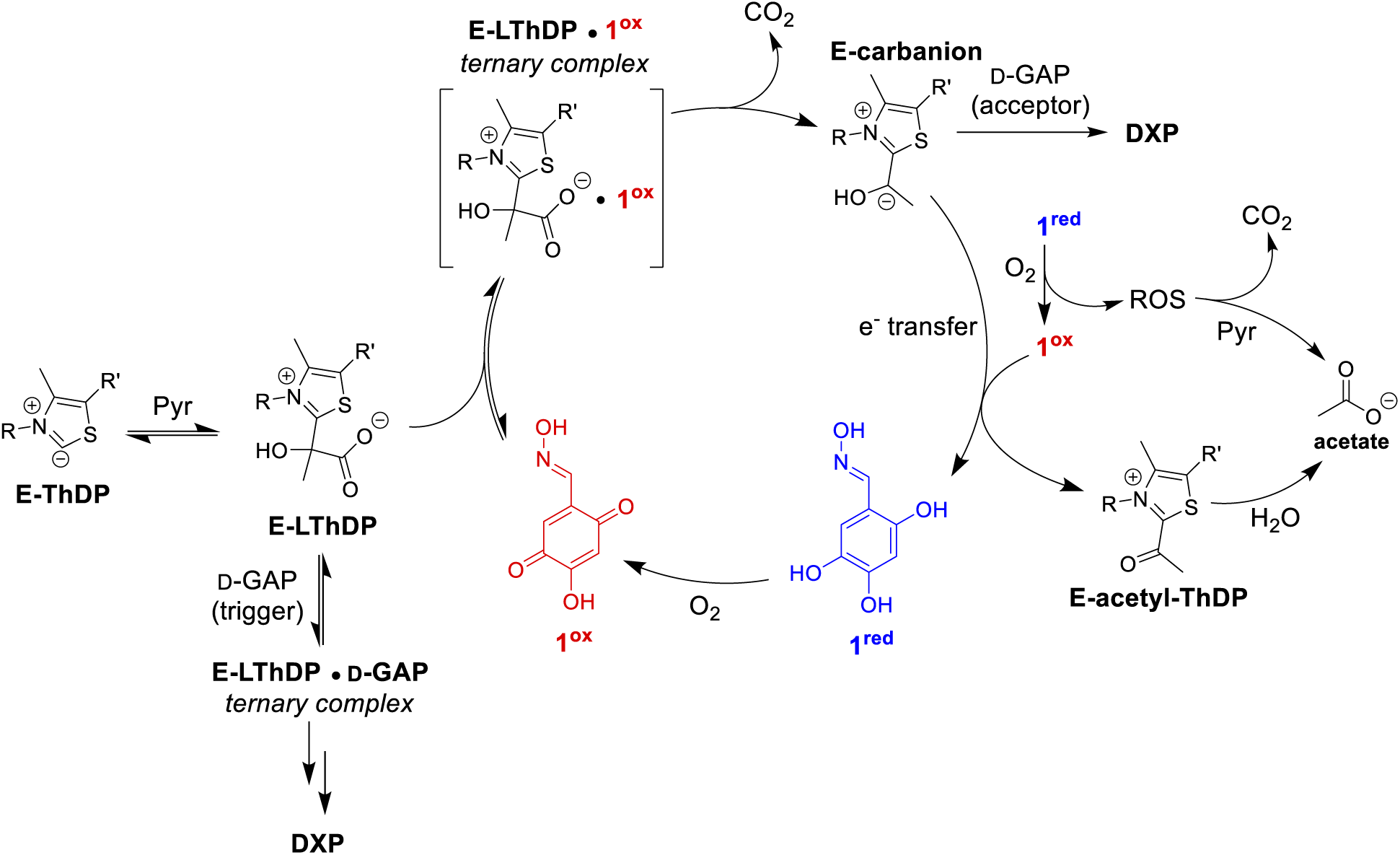
Working model for redox cycling mechanism of oxime-based inhibitors of DXPS and nonenzymatic oxidative decarboxylation of pyruvate.

## Discussion

This work investigated the selectivity and mechanism of inhibition of previously discovered trihydroxybenzaldoxime inhibitors of DXPS. The observed selectivity of **1** for multiple DXPS homologs over PDC suggests that it targets the unique ligand-gated mechanism of DXPS. The modestly reduced potency of **1** against *Pa*DXPS relative to *Ec*DXPS or *Dr*DXPS is not surprising given the recent discovery that the bisubstrate analog inhibitor D-PheTrAP also exhibits lower potency and different behavior on *Pa*DXPS relative to *Ec*DXPS and *Dr*DXPS.^37^ While *Ec*R99 contributes to the time-dependent behavior of D-PheTrAP on *Ec*DXPS, the analogous residues on *Pa*DXPS, R106, does not. This is notable given our finding that R99 contributes to the potency of **1** on *Ec*DXPS. Perhaps *Ec*R99 and *Pa*R106 are positioned differently in the distinct E-LThDP complexes of *Ec*DXPS and *Pa*DXPS recognized by **1**, which could contribute to the reduced potency of **1** on *Pa*DXPS and suggests it may be possible to design species-specific uncompetitive inhibitors of DXPS. Despite their close proximity to R99, substitution of other active site network residues did not lead to reduced potency of **1**; in fact, **1** was more potent against *Ec*H431A. These variants are known to readily activate LThDP to the C2-α-carbanion in the absence of a trigger molecule. Oxime **1** may still bind and accept electrons from the C2-α-carbanion on these variants; it follows that these substitutions would not necessarily negatively affect potency. Thus, we speculate that either R99 interacts with **1**, or substitution of R99 alters the conformation of the E-LThDP complex in a manner which lowers affinity or reduces redox activity of **1** in an altered active site environment.

In studies to determine how oxime inhibitors may interact with the E-LThDP complex, we discovered nonenzymatic redox activity of **1** which may be relevant to its inhibitory activity and could challenge our the idea that these oximes are active in their catechol forms under aerobic conditions.^32^ The generation of ROS via autoxidation of catechols under alkaline conditions is known,^44–46,49^ as is the reaction of pyruvate with ROS and the antioxidant properties of ketoacids.^51,62^ Ultimately, the current study revealed an oxygen-dependent redox cycling mechanism in which **1** in its oxidized form induces multiple turnovers of DXPS and accepts electrons from the C2α-carbanion, preventing DXP formation by competing with D-GAP. The oxidized form of **1** is, in fact, a more potent inhibitor than **1** in its reduced form. The previously observed low μM inhibition by **1** under aerobic conditions in the presence of ascorbate^32^ is now explained by this mechanism: Ascorbate promotes the redox cycle in the presence of O_2_ (as observed previously with other quinones)^53^, reducing the oxidized form of **1** following autoxidation of the catechol. ROS generation during this process reacts with pyruvate, reducing substrate levels which can, in turn, influence the measured potency.

Although the oxidized form of **1** was found to be more potent than the catechol, weak inhibitory activity of the catechol of **1** was nevertheless observed, yet its mechanism of inhibition is not fully understood. Perhaps the catechol binds to E-LThDP or E-carbanion and is capable of blocking D-GAP binding. It is unknown if the catechol induces LThDP decarboxylation; if so, we would expect acetate to be the product of pyruvate as we observed in a recent study of inducers of LThDP decarboxylation.^24^ Alternatively, residual O_2_ present under anaerobic conditions could promote redox cycling at a low level to account for the observed weak potency.

The redox activity of ThDP-dependent enzymes is well-studied.^31,60,63–66^ ThDP-dependent enzymes have been reported to catalyze reactions involving single electron transfer and radical recombination.^31,63,64,67^ Redox dye reporters such as DCPIP are commonly used to assess the activity of enzymes such as PDH by detection of carbanion formation. Similarly, DXPS can reduce O_2_ to superoxide as well as donate electrons to redox reporters such as PMS/MTT.^24,31^

Thus, it is not surprising that **1** engages in a redox cycling mechanism with DXPS, and this mechanism could have potential for future development of redox-active DXPS inhibitors. Redox cycling antimicrobial agents already have precedent in the clinic. Nitrofuran antimicrobials act by a redox cycling mechanism, undergoing activation by microbial nitroreductases and resulting in the generation of ROS, similar to **1**.^68,69^ Off-target activity of these drugs is associated with a number of side effects, but their clinical efficacy and low resistance are promising for this class of redox cycling antimicrobials. By consuming pyruvate necessary for critical metabolite precursor synthesis and generating damaging ROS, oxime inhibitors of DXPS display promising characteristics for drug development. Additionally, the promotion of acetate formation by **1** along with previous observations of acetate formation catalyzed by DXPS^16,24^ could hint at potential links between DXPS and bacterial acetate metabolism.^70,71^ The apparent selectivity of **1** for DXPS over PDC and its ability to engage the uniquely stable DXPS-LThDP complex presents a potentially interesting opportunity to develop redox cycling inhibitors that are selective for DXPS.

The redox activity of DXPS now characterized with O_231_ and trihydroxybenzaldoximes, and our observation of oxidative decarboxylation of pyruvate in the absence of O_2_,^16^ could suggest that DXPS plays a role in managing other redox sensitive components in bacteria, or is utilized by bacteria for production of acetate. Our results are significant because they highlight yet another interesting activity of DXPS which has important implications for inhibitor design and offers potential insights into DXPS target function in pathogenic bacteria.

## Materials and Methods

### General Methods

Unless otherwise noted, all materials were obtained from commercial sources. Glyceraldehyde 3-phosphate (GAP) was purchased as a racemic mixture (Sigma#G5251, CAS# 591-59-3) and used without further purification. In this mixture, D-GAP is the preferred substrate for DXPS, thus, all reported GAP concentrations refer to D-GAP. Pyruvate decarboxylase (PDC) from baker’s yeast (*S. cerevisiae*, Sigma#P9474, CAS# 9001-04-1), Catalase from bovine liver (CAT, Sigma#C9322, CAS# 9001-05-2) and alcohol dehydrogenase (ADH, Sigma#55689, CAS# 9031-72-5) were purchased from Sigma Aldrich. *E. coli*, *P. aeruginosa*, and *D. radiodurans* DXPS, and *E. coli* IspC were overexpressed and purified as previously reported.^22,37,72^ The preparation of R478A, R99A, H431A, and D427A variants has been previously reported.^16,27^ A vinyl anaerobic chamber (Coy Laboratory Products; Grass Lake, MI) was used to deoxygenate samples and/or conduct anaerobic experiments. Aerobic kinetic characterizations were performed using a BioTek Epoch 2 microplate reader. Anaerobic kinetic characterizations were performed using a Tecan Infinite 200 Pro microplate reader. CH Instruments Electrochemical Analyzer (CHI 1040C, Austin, TX) multichannel potentiostats and associated software were used for all electrochemical measurements.

### Synthesis

Unless noted, all compounds were prepared as previously described^32^ from commercially available starting materials. Relevant chromatograms and spectra are located in the supplementary information (S29-38).

#### Determination of inhibitor purity

Reversed phase (RP) HPLC and high field NMR (500 MHz) were used to determine purity of new DXPS inhibitors (compounds **3** and **8**) presented in this study. A fresh 0.125 mM solution of inhibitor in water (**8**) or methanol (**3**) prepared immediately prior to injection. Then, 5 uL of the 0.125 mM inhibitor solution was injected onto a ZORBAX 80Å extend-C18 column (4.6 x 50 mm, 3.5 μm) with a flow rate of 1 mL/min. Mobile phases used include water (solvent A) and acetonitrile (solvent B). The following method was used: 30 seconds at 0% B, ramp to 95% B over 5.5 minutes, and hold 95% B for 60 seconds, then back down to 0% B over 30 seconds and hold for 30 seconds. Compound **3** was evaluated at 380 nm and compound **8** was evaluated at 320 nm using the HPLC diode array detector (DAD).

#### Assignment of redox states and geometric isomers

Due to the ability of oxime **1** and related compounds to exist in multiple oxidation state/geometric isomers depending on reaction conditions, we have made tentative assignments in the characterization, based on collected data and predictions based on literature precedence. We predict that oxime **1** exists as the *para*-quinone in its oxidized form based on precedence for stability of the *para-*quinone over the *ortho*-quinone.^47^ NMR peak assignments for oxime **1** and **8** are supported by ^2^*J*_NH_ and ^4^*J*_NH_ couplings from LR HSQC experiments (Fig 8), and peaks assignments for the reduced forms are supported by experiments conducted in the presence of ascorbate (Fig S10, S19). The characterization of **3** and its geometric isomers is based on peak integration ratios and the prediction that the *E* isomer would be favored on the basis of reduced steric hindrance.

### (*E,Z*)-5-(hydrazineylidenemethyl)benzene-1,2,4-triol (3)

A solution of 2,4,5-trihydroxybenzaldehyde (0.100 g, 0.649 mmol) and hydrazine (0.020 mL, 0.649 mmol) in ethanol (1.0 mL) was heated to reflux for 2 hours. The solution was allowed to cool to ambient temperature, and the solvent was removed under reduced pressure. The resulting solid was purified via silica flash chromatography (ethyl acetate/hexanes/methanol 60:39:1 – 100% ethyl acetate) to yield an orange powder (0.063g, 57% yield) (Fig S29-33). (E isomer) R_T_ = 2.54 min λ_max_ = 262 nm. (Z isomer) R_T_ = 2.88 min λ_max_ = 380 nm ^1^H NMR (500 MHz, MeOD-d_4_) δ (E isomer) 7.80 (s, 1H), 6.54 (s, 1H), 6.27 (s, 1H); δ (*Z* isomer) 8.5 (s, 1H), 6.77 (s, 1H), 6.34 (s, 1H); ^13^C NMR (126 MHz, DMSO-d6) δ (*E* isomer) 151.1, 147.0, 143.8, 138.3, 115.1, 111.0, 103.8; (*Z* isomer) 161.5, 154.0, 151.4, 139.2, 116.5, 109.5, 103.8. HRMS (ESI) m/z: (**3**, reduced) calc’d 169.0613 (M+H^+^); found 169.0612 (M+H^+^) (**3**, oxidized) calc’d 167.0457 (M+H^+^); found 167.0452 (M+H^+^)

### 15N-labeled (E)-2,4,5-Trihydroxybenzaldehyde oxime (8)

A solution of 2,4,5-trihydroxybenzaldehyde (0.100 g, 0.649 mmol), ^15^N-hydroxyl amine hydrochloride (0.053 g, 0.715 mmol, Toronto Research Chemicals Inc, Toronto, Canada), and sodium acetate (0.0852 g, 1.04 mmol) was prepared in methanol (1.5 mL) and water (35 drops). The reaction mixture was stirred overnight at room temperature. The crude reaction solution was purified directly via silica flash chromatography (ethyl acetate/hexanes 1:1 isocratic) to yield a light brown powder (0.0975 g, 88% yield) (Fig S34-38). R_T_ = 2.65 min λ_max_ = 254 nm. ^1^H NMR (500 MHz, MeOD-d_4_) δ (E isomer) 8.00 (s, 1H), 6.62 (s, 1H), 6.30 (s, 1H). ^13^C NMR (126 MHz, MeOD) δ (*E* isomer) 151.5, 150.6, 148.0, 138.1, 115.3, 108.4, 102.8. HRMS (ESI) m/z: (**8**, reduced) calc’d 171.0424 (M+H^+^); found 171.0423 (M+H^+^) (**8**, oxidized) calc’d 169.0267 (M+H^+^); found 169.0270 (M+H^+^)

### Kinetic Characterization of Inhibitors

#### General conditions for DXPS-IspC coupled assay

Inhibitory activity of compounds against DXPS were monitored via a continuous spectrophotometric coupled assay with DXPS and IspC as previously described.^41,42^ Briefly, Reaction mixtures containing HEPES (100 mM, pH 8.0), MgCl_2_ (2 mM), NaCl (5 mM), ThDP (1 mM), BSA (1 mg/mL), DXPS (125-500 nM), IspC (1.7 µM), NADPH (160 µM), DMSO (5%, v/v), and inhibitor (0-200 µM) were preincubated at 25 °C (or chamber temperature 25-30 °C) for 5 min. Enzymatic reactions were initiated via the addition of 20 µL of a 10x substrate mix for a final well volume of 200 µL. Reactions were performed in triplicate unless otherwise noted. See the following sections for details and modifications.

#### Aerobic Apparent K_i_

When varying pyruvate (0-500 µM), GAP was maintained at 50 µM (3 mM for R478A). When varying GAP (0-500 µM), pyruvate was maintained at 500 µM. The following [DXPS] were used when varying **1**: WT *Ec*DXPS, R478A, H431A, D427A (200 nM), R99A (500 nM), WT *Dr*DXPS, WT *Pa*DXPS (150 nM). Analogs **3** and **4** were tested using WT *Ec*DXPS (150 nM). Reactions were incubated and initiated as described above. Immediately following initiation, NADPH depletion was monitored at 340 nm at 25 ℃ for 7-10 min with a 14-20 second time interval. Initial velocities of DXP formation were determined from the linear portion of the NADPH decay. Initial velocities were arithmetically corrected for background for compound **4**. Initial velocities were plotted vs. substrate concentration, and the data were fit to equation 1 (mixed model), 2 (competitive), or 3 (uncompetitive) based on the calculated α value and goodness of fit in GraphPad Prism. Reported standard errors of the mean for *K*_i_s are calculated from three replicates.

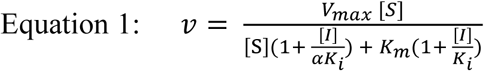

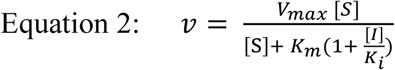

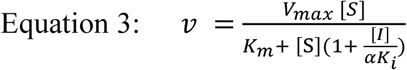

#### Anaerobic Apparent K_i._

Reaction components were brought into the anaerobic chamber and allowed to deoxygenate for ∼2 hours. WT *Ec*DXPS was used at 125 nM. When varying pyruvate (0-500 µM), GAP was maintained at 50 µM. When varying GAP (0-500 µM), pyruvate was maintained at 500 µM. Reactions were incubated and initiated as described above. Immediately following initiation, NADPH depletion was monitored at 340 nm at chamber temperature (25-30℃) for 5 min with a ∼17 second time interval. Initial velocities of DXP formation were determined as above, plotted vs. substrate concentration, and the data were fit to either equation 2, or 3 based on the calculated α value and goodness of fit in GraphPad Prism. Reported standard errors of the mean for *K*_i_s are calculated from three replicates.

#### Inhibition in the presence of CAT or ascorbate

Reaction solutions were prepared containing *Ec*DXPS (150 nM). Solutions contained either DXPS alone or were supplemented with CAT (0.25 mg/mL) or ascorbate (5 mM). Reaction mixtures were incubated with either water or **1** (100 µM) for 5 min at 25 ℃. Reactions were initiated with a mixture of pyruvate (110 µM) and GAP (60 µM), or water for background controls. Immediately following initiation, NADPH depletion was monitored at 340 nm at 25 ℃ for 7.5 min with a 14 second time interval. Initial velocities of DXP formation were measured as described above and arithmetically corrected for background rates and normalized to the DXPS only control reaction for comparison. Reactions were performed in triplicate, and % inhibition was determined arithmetically from initial velocities. Reported standard errors of the mean for *K*_i_s are calculated from three replicates.

#### Inhibition of PDC

Reaction conditions were optimized from a previously reported assay.^38^ Briefly, reaction mixtures containing MES (100 mM, pH 6.2), MgCl_2_ (2 mM), ThDP (5 mM), PDC (0.06 U/mL), ADH (16 U/mL), NADH (170 µM), and **1** (0 or 200 µM) were preincubated at 25 °C for 5 min. Enzymatic reactions were initiated via the addition of 20 µL of 10x pyruvate (final 1mM), or water for background controls, for a final well volume of 200 µL. Immediately following initiation, NADH depletion was monitored at 340 nm at 25 ℃ for 7 min with a 14 second time interval. Initial velocities of acetaldehyde formation were measured and arithmetically corrected for background rates. Standard errors of reported % inhibition values were determined from 3 replicates. pH controls were conducted with phosphate buffer (pH 7 and 8); background subtraction was not performed on these samples. Reported standard errors of the mean for *K*_i_s are calculated from three replicates.

#### Inhibition of ADH

Reaction mixtures containing MES (100 mM, pH 6.2), MgCl_2_ (2 mM), ThDP (5 mM), ADH (16 U/mL), NADH (170 µM), and **1** (0 or 200 µM) were preincubated at 25 °C for 5 min. Enzymatic reactions were initiated via the addition of 20 µL of 10x acetaldehyde (final 1mM), or water for background controls, for a final well volume of 200 µL. Immediately following initiation, NADH depletion was monitored at 340 nm at 25 ℃ for 6 min with a 11 second time interval. Initial velocities of ethanol formation were measured as above and arithmetically corrected for background rates. Reported standard errors of the mean for *K*_i_s are calculated from three replicates.

### NMR detection of redox state of 1 and products of nonenzymatic decarboxylation of pyruvate

NMR experiments were performed as previously described^16,24^ with the following modifications. Experiments were performed aerobically unless otherwise noted. Anaerobic experiments were performed in the anaerobic chamber up to the point of transfer to NMR tubes which were capped with rubber septa to exclude oxygen once removed from the chamber. To monitor nonenzymatic decarboxylation of pyruvate, 500 µM sodium pyruvate-^13^C_3_ was added to buffer (50 mM HEPES pH 8, Tris pH 8, phosphate pH 8, MES pH 6.2, CHES pH 9, or Phosphate pH 7) containing 10% D_2_O and 250 µM gadobutrol (MedChemExpress, NJ). Solutions were also prepared aerobically in the presence of ascorbate (5 mM) and CAT (0.25 mg/mL). Oxime **1** (500 µM) was then added and solutions were incubated at 4 ℃ for 20 min.

After incubation, 600 µL of each solution were transferred to labeled NMR tubes. 1D ^1^H-NMR and 1D ^1^H detected H{^13^C} and H_3_{^13^C} spectra via a proton multiplicity filter, which selects for ^13^C nuclei attached to an odd number of protons, were acquired on a Bruker Avance 600 MHz or a Bruker Avance NEO 600 MHz spectrometer (HEPES anaerobic and Tris anaerobic) equipped with a TCI cryogenic probe equipped with z-gradients with a cryo-cooled ^13^C preamplifier, resulting in high sensitivity for ^13^C detection. Spectra were acquired at 600 MHz (^1^H) and 150 MHz (^13^C), at 25 °C utilizing a 90° ^1^H excitation pulse (optimized for maximum signal-to-noise per unit time), 128 scans/FID (^1^H-NMR), 200 ms acquisition time/FID, and a 3s relaxation day between successive scans, or 64 scans/FID (^1^H detected H{^13^C} and H_3_{^13^C}), 200 ms acquisition time/FID, and a 5s relaxation day between successive scans.

### NMR detection of chemical reduction of 1 by DXPS and substrates

Reaction buffer (50 mM Tris pH 8, 2 mM MgCl_2_, 100 mM NaCl, 1 mM ThDP, 10% D_2_O) was prepared aerobically. Oxime **8** (500 µM) was added, and the mixture was incubated at 4 ℃ for 30 min. Reaction mixtures were then brought into the anaerobic chamber and allowed to deoxygenate ∼2 hrs. Reactions were initiated by addition of 1 µM DXPS with 2 mM sodium pyruvate-^13^C_3_ or 2 mM sodium pyruvate-^13^C_3_ and 500 µM GAP. Reaction mixtures were incubated at 25 ℃ for 1 hr. Reactions were quenched by boiling for 5 min at 95 ℃ followed by vortexing for 15 seconds. The quenched mixtures were transferred to labeled NMR tubes which were sealed with rubber septa and parafilm prior to removal from the chamber to exclude oxygen once removed from the chamber. 2D Multiple Bond Long Range (LR) ^1^H-^15^N-HSQC and 1D ^1^H detected H{^13^C} and H_3_{^13^C} spectra via a proton multiplicity filter which selects for ^13^C nuclei attached to an odd number of protons were acquired on a Bruker Avance NEO 600 MHz spectrometer equipped with a TCI cryogenic probe equipped with z-gradients with a cryo-cooled ^13^C preamplifier, resulting in high sensitivity for ^13^C detection. Spectra were acquired at 600 MHz (^1^H), 150 MHz (^13^C), and 61 MHz (^15^N) at 25 °C. Parameters for various categories of spectra are as follows: (LR ^1^H-^15^N HSQC): 16 scans and 150 ms acquisition time (t_2_) per FID and a ^1^H spectral width of 15.1376 ppm; since the chemical shifts of N^R^ (δ 335) and N^O^ (δ 374) are separated by ∼ 40 ppm, data collection times were optimized by positioning the ^15^N carrier at 354 ppm and using an ^15^N spectral width of 25 ppm and an acquisition time of 13 ms along ^15^N (t_1_); with a 1.5 s relaxation delay between successive scans, the total data collection period per 2D spectrum was 20 minutes. ^1^H detected H{^13^C} and H_3_{^13^C} filtered 1D spectra: 200 ms acquisition time/FID, and a 5s relaxation day between successive scans, 1.5 minutes per spectrum. Controls were also prepared in which reactions were initiated by addition of 1 µM DXPS with no substrates, as well as 2 mM ^13^C-labeled pyruvate + 500 µM GAP with DXPS dialysis buffer in place of enzyme. A “mixed” redox state control of **1** (500 µM) was prepared aerobically in 50 mM HEPES pH 8 with 10% D_2_O.

### Electrochemical measurements

#### Cyclic Voltammetry

Solutions of **1**, **2**, **5**, **6**, and **7** (1 mM) were prepared in 100 mM phosphate buffer pH 8 and 100 mM NaCl. A three-electrode cell configuration consisting of a glassy carbon disk (5 mm diameter) working electrode, a coiled platinum wire counter electrode, and an Ag|AgCl reference electrode was used. Cyclic voltammetry measurements used a voltage window from either 0.4 to −0.8 V (**1**, **2**, **7**), 0.6 to −0.8 V (**5**, **6**) or 0.8 to −1.2 V (Blank) and all measurements used a scanning rate of 0.1 V/s after a quiet time of 2 s starting with a negative sweep.

#### Continuous monitoring of **1**

Reaction solutions were prepared containing 2 mM MgCl_2_, 100 mM NaCl, 50 mM Tris pH 8, 1 mM ThDP, and **1** (final 1 mM) under aerobic conditions to allow for oxidation of **1**. Solutions were transferred to a cell prepared from a 15 mL centrifuge tube and capped with a rubber septum. Solutions were deoxygenated by bubbling with argon. A baseline CV was collected for the reaction with DXPS, pyruvate, and GAP prior to initiation. In the anaerobic chamber, solutions of DXPS, pyruvate, GAP, water, and DXPS dialysis buffer were allowed to deoxygenate for at least 2 hours. Amperometric i-t curve measurements were started prior to initiation. Just before initiation, reaction initiation solutions containing either DXPS (final 10 µM) + pyruvate (final 2 mM) + GAP (final 500 µM), DXPS (final 10 µM) + pyruvate (final 2 mM), DXPS (final 10 µM) + water, or DXPS dialysis buffer + pyruvate (final 2 mM) + GAP (final 500 µM) were prepared and taken up via gas tight Hamilton syringe and capped for transfer outside of the chamber. Reactions were initiated and mixed via gas tight Hamilton syringe at ∼300 s and measured for at least 200 s. A CV after the reaction with DXPS, pyruvate, and GAP was collected. A three-electrode cell configuration consisting of a gold ultra micro working electrode (6.25 µm diameter), a platinum wire counter electrode, and an Ag|AgCl reference electrode was used. Cyclic voltammetry measurements used a voltage window from 0.4 to −0.6 V and a scanning rate of 0.02 V/s after a quiet time of 10 s starting with a negative sweep. Amperometric i-t curve measurements were collected using −0.5 V with a sample interval of 0.1 s after a quiet time of 0 s. The number of electrons being transferred in the electrochemical reduction measured at 0.5 V was determined via the following equation where i_ss_ is the steady state current, F is the Faraday constant, d is the diffusion coefficient (estimated as 5×10^-7^), c is the concentration of the analyte, and a is the radius of the electrode.

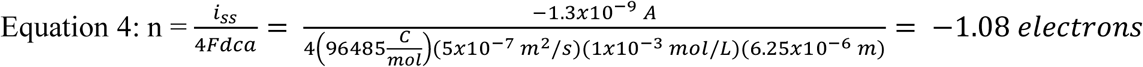

## Acknowledgements

We would like to thank Eucolona M. Toci for providing *Ec*R99A, *Ec*H431A, and *Ec*D427A and Jacob Weber for assistance in the collection of CVs. This work was supported by the National Institute of Health grants R01GM143810 and T32GM080189, and NSF DGE2139757. ^1^H and ^13^C NMR spectra for **3** and **8** were acquired using the JHU-Pharmacology JEOL JNM-ECZL500R spectrometer and we are thankful to the NIH for supporting the NMR facility through award number S10OD034217. We thank the University of Illinois Urbana-Champaign School of Chemical Sciences Mass Spectrometry Lab for assistance in collecting HRMS data for compounds **3** and **8**.

## Supporting Information

Inhibition of *Ec*, *Pa*, and *Dr*DXPS by **1**; Inhibition of ADH and PDC at pH 7 and 8 by **1**; Inhibition of *Ec* R478A, D427A, H431A, and R99A by **1**; NMR replicates of nonenzymatic decarboxylation of pyruvate in the presence of **1**; Mechanism of autoxidation and production of ROS; NMR replicates of pH dependence of the autoxidation of **1**; NMR replicates of the redox activity of **1** with CAT and ascorbate; Mechanism of redox cycling of **1** with ascorbate; NMR of **1** (12.5 mM) in HEPES pH 8; Anaerobic inhibition of *Ec*DXPS by **1**; Inhibition of *Ec*DXPS by **3** and **4**; Cyclic voltammograms of **1-2**, **5-7**; Experimental scheme for the detection of the reduction of **8** by DXPS and substrates monitored by LR HSQC; NMR replicates of LR ^1^H-^15^N HSQC experiments; NMR replicates of DXPS product formation experiments; Raw amperoteric i-t curves of **1** with DXPS and substrates; Cyclic voltammogram of **1** before and after reaction with DXPS and substrates; characterization of **3** and **8**; Tables of inhibition by **1**-**4** (DOC)

## Author Contributions

C.R.N and C.L.F.M designed the study. C.R.N performed all synthesis, biochemical, biophysical, and electrochemical characterization of compounds and activity and collected all NMR spectra and CVs. A.M designed and optimized NMR experiment parameters and assisted in NMR acquisition. N.A.C designed and assisted with electrochemical measurements. C.R.N. and C.L.F.M prepared the manuscript.

## References

(1) GBD 2021 Antimicrobial Resistance Collaborators. Global Burden of Bacterial Antimicrobial Resistance 1990–2021: A Systematic Analysis with Forecasts to 2050. Lancet 2024, 404 (10459), 1199–1226. 10.1016/S0140-6736(24)01867-1.

(2) Murima, P.; McKinney, J. D.; Pethe, K. Targeting Bacterial Central Metabolism for Drug Development. Chem. Biol. 2014, 21 (11), 1423–1432. 10.1016/j.chembiol.2014.08.020.

(3) Tong, M.; Brown, E. D. Food for Thought: Opportunities to Target Carbon Metabolism in Antibacterial Drug Discovery. Ann. N. Y. Acad. Sci. 2023. 10.1111/nyas.14991.

(4) Stokes, J. M.; Lopatkin, A. J.; Lobritz, M. A.; Collins, J. J. Bacterial Metabolism and Antibiotic Efficacy. Cell Metab. 2019, 30 (2), 251–259. 10.1016/j.cmet.2019.06.009.

(5) Shen, Y.; Liu, J.; Estiu, G.; Isin, B.; Ahn, Y.-Y.; Lee, D.-S.; Barabási, A.-L.; Kapatral, V.; Wiest, O.; Oltvai, Z. N. Blueprint for Antimicrobial Hit Discovery Targeting Metabolic Networks. Proc. Natl. Acad. Sci. U. S. A. 2010, 107 (3), 1082–1087. 10.1073/pnas.0909181107.

(6) Golden, M. M.; Post, S. J.; Rivera, R.; Wuest, W. M. Investigating the Role of Metabolism for Antibiotic Combination Therapies in Pseudomonas Aeruginosa. ACS Infect. Dis. 2023. 10.1021/acsinfecdis.3c00452.

(7) Rohmer, M.; Knani, M.; Simonin, P.; Sutter, B.; Sahm, H. Isoprenoid Biosynthesis in Bacteria: A Novel Pathway for the Early Steps Leading to Isopentenyl Diphosphate. Biochem. J 1993, 295 ( Pt 2) (Pt 2), 517–524. 10.1042/bj2950517.

(8) Jomaa, H.; Wiesner, J.; Sanderbrand, S.; Altincicek, B.; Weidemeyer, C.; Hintz, M.; Türbachova, I.; Eberl, M.; Zeidler, J.; Lichtenthaler, H. K.; Soldati, D.; Beck, E. Inhibitors of the Nonmevalonate Pathway of Isoprenoid Biosynthesis as Antimalarial Drugs. Science 1999, 285 (5433), 1573–1576. 10.1126/science.285.5433.1573.

(9) Lois, L. M.; Campos, N.; Putra, S. R.; Danielsen, K.; Rohmer, M.; Boronat, A. Cloning and Characterization of a Gene from Escherichia Coli Encoding a Transketolase-like Enzyme That Catalyzes the Synthesis of D-1-Deoxyxylulose 5-Phosphate, a Common Precursor for Isoprenoid, Thiamin, and Pyridoxol Biosynthesis. Proc. Natl. Acad. Sci. U. S. A. 1998, 95 (5), 2105–2110. 10.1073/pnas.95.5.2105.

(10) Sprenger, G. A.; Schörken, U.; Wiegert, T.; Grolle, S.; de Graaf, A. A.; Taylor, S. V.; Begley, T. P.; Bringer-Meyer, S.; Sahm, H. Identification of a Thiamin-Dependent Synthase in Escherichia Coli Required for the Formation of the 1-Deoxy-D-Xylulose 5-Phosphate Precursor to Isoprenoids, Thiamin, and Pyridoxol. Proc. Natl. Acad. Sci. U. S. A. 1997, 94 (24), 12857–12862. 10.1073/pnas.94.24.12857.

(11) Du, Q.; Wang, H.; Xie, J. Thiamin (Vitamin B1) Biosynthesis and Regulation: A Rich Source of Antimicrobial Drug Targets? Int. J. Biol. Sci. 2011, 7 (1), 41–52. 10.7150/ijbs.7.41.

(12) Schwarz, M. K. Terpene Biosynthesis in Ginkgo Biloba: A Surprising Story. ETH-Zurich Dissertation 1994.

(13) Patel, M. S.; Nemeria, N. S.; Furey, W.; Jordan, F. The Pyruvate Dehydrogenase Complexes: Structure-Based Function and Regulation. J. Biol. Chem. 2014, 289 (24), 16615–16623. 10.1074/jbc.R114.563148.

(14) Frank, R. A. W.; Leeper, F. J.; Luisi, B. F. Structure, Mechanism and Catalytic Duality of Thiamine-Dependent Enzymes. Cell. Mol. Life Sci. 2007, 64 (7–8), 892–905. 10.1007/s00018-007-6423-5.

(15) Frank, R. A. W.; Titman, C. M.; Pratap, J. V.; Luisi, B. F.; Perham, R. N. A Molecular Switch and Proton Wire Synchronize the Active Sites in Thiamine Enzymes. Science 2004, 306 (5697), 872–876. 10.1126/science.1101030.

(16) Toci, E. M.; Austin, S. L.; Majumdar, A.; Lee Woodcock, H.; Freel Meyers, C. L. Disruption of an Active Site Network Leads to Activation of C2α-Lactylthiamin Diphosphate on the Antibacterial Target 1-Deoxy-D-Xylulose-5-Phosphate Synthase. Biochemistry 2024, 63 (5), 671–687. 10.1021/acs.biochem.3c00735.

(17) Brammer, L. A.; Smith, J. M.; Wade, H.; Meyers, C. F. 1-Deoxy-D-Xylulose 5-Phosphate Synthase Catalyzes a Novel Random Sequential Mechanism. J. Biol. Chem. 2011, 286 (42), 36522–36531. 10.1074/jbc.M111.259747.

(18) Handa, S.; Dempsey, D. R.; Ramamoorthy, D.; Cook, N.; Guida, W. C.; Spradling, T. J.; White, J. K.; Woodcock, H. L.; Merkler, D. J. Mechanistic Studies of 1-Deoxy-D-Xylulose-5-Phosphate Synthase from Deinococcus Radiodurans. Biochem Mol Biol J 2018, 4 (1). 10.21767/2471-8084.100051.

(19) Patel, H.; Nemeria, N. S.; Brammer, L. A.; Freel Meyers, C. L.; Jordan, F. Observation of Thiamin-Bound Intermediates and Microscopic Rate Constants for Their Interconversion on 1-Deoxy-D-Xylulose 5-Phosphate Synthase: 600-Fold Rate Acceleration of Pyruvate Decarboxylation by D-Glyceraldehyde-3-Phosphate. J. Am. Chem. Soc. 2012, 134 (44), 18374–18379. 10.1021/ja307315u.

(20) Eubanks, L. M.; Poulter, C. D. Rhodobacter Capsulatus 1-Deoxy-D-Xylulose 5-Phosphate Synthase: Steady-State Kinetics and Substrate Binding. Biochemistry 2003, 42 (4), 1140–1149. 10.1021/bi0205303.

(21) Tittmann, K.; Golbik, R.; Uhlemann, K.; Khailova, L.; Schneider, G.; Patel, M.; Jordan, F.; Chipman, D. M.; Duggleby, R. G.; Hübner, G. NMR Analysis of Covalent Intermediates in Thiamin Diphosphate Enzymes. Biochemistry 2003, 42 (26), 7885–7891. 10.1021/bi034465o.

(22) Chen, P. Y.-T.; DeColli, A. A.; Freel Meyers, C. L.; Drennan, C. L. X-Ray Crystallography-Based Structural Elucidation of Enzyme-Bound Intermediates along the 1-Deoxy-d-Xylulose 5-Phosphate Synthase Reaction Coordinate. J. Biol. Chem. 2019, 294 (33), 12405–12414. 10.1074/jbc.RA119.009321.

(23) Nemeria, N. S.; Chakraborty, S.; Balakrishnan, A.; Jordan, F. Reaction Mechanisms of Thiamin Diphosphate Enzymes: Defining States of Ionization and Tautomerization of the Cofactor at Individual Steps. FEBS J. 2009, 276 (9), 2432–2446. 10.1111/j.1742-4658.2009.06964.x.

(24) Toci, E. M.; Majumdar, A.; Meyers, C. L. F. Aldehyde-Based Activation of C2α-Lactylthiamin Diphosphate Decarboxylation on Bacterial 1-Deoxy-d-Xylulose 5-Phosphate Synthase. Chembiochem 2024, 25 (23), e202400558. 10.1002/cbic.202400558.

(25) Battistini, M. R.; Shoji, C.; Handa, S.; Breydo, L.; Merkler, D. J. Mechanistic Binding Insights for 1-Deoxy-D-Xylulose-5-Phosphate Synthase, the Enzyme Catalyzing the First Reaction of Isoprenoid Biosynthesis in the Malaria-Causing Protists, Plasmodium Falciparum and Plasmodium Vivax. Protein Expr. Purif. 2016, 120, 16–27. 10.1016/j.pep.2015.12.003.

(26) White, J. K.; Handa, S.; Vankayala, S. L.; Merkler, D. J.; Woodcock, H. L. Thiamin Diphosphate Activation in 1-Deoxy-d-Xylulose 5-Phosphate Synthase: Insights into the Mechanism and Underlying Intermolecular Interactions. J. Phys. Chem. B 2016, 120 (37), 9922–9934. 10.1021/acs.jpcb.6b07248.

(27) Basta, L. A. B.; Patel, H.; Kakalis, L.; Jordan, F.; Meyers, C. L. F. Defining Critical Residues for Substrate Binding to 1-Deoxy-D-Xylulose 5-Phosphate Synthase--Active Site Substitutions Stabilize the Predecarboxylation Intermediate C2α-Lactylthiamin Diphosphate. FEBS J. 2014, 281 (12), 2820–2837. 10.1111/febs.12823.

(28) Sisquella, X.; de Pourcq, K.; Alguacil, J.; Robles, J.; Sanz, F.; Anselmetti, D.; Imperial, S.; Fernàndez-Busquets, X. A Single-Molecule Force Spectroscopy Nanosensor for the Identification of New Antibiotics and Antimalarials. FASEB J. 2010, 24 (11), 4203–4217. 10.1096/fj.10-155507.

(29) Brammer, L. A.; Meyers, C. F. Revealing Substrate Promiscuity of 1-Deoxy-D-Xylulose 5-Phosphate Synthase. Org. Lett. 2009, 11 (20), 4748–4751. 10.1021/ol901961q.

(30) Morris, F.; Vierling, R.; Boucher, L.; Bosch, J.; Freel Meyers, C. L. DXP Synthase-Catalyzed C-N Bond Formation: Nitroso Substrate Specificity Studies Guide Selective Inhibitor Design. Chembiochem 2013, 14 (11), 1309–1315. 10.1002/cbic.201300187.

(31) DeColli, A. A.; Nemeria, N. S.; Majumdar, A.; Gerfen, G. J.; Jordan, F.; Freel Meyers, C. L. Oxidative Decarboxylation of Pyruvate by 1-Deoxy-d-Xyulose 5-Phosphate Synthase, a Central Metabolic Enzyme in Bacteria. J. Biol. Chem. 2018, 293 (28), 10857–10869. 10.1074/jbc.RA118.001980.

(32) Bartee, D.; Morris, F.; Al-Khouja, A.; Freel Meyers, C. L. Hydroxybenzaldoximes Are D-GAP-Competitive Inhibitors of E. Coli 1-Deoxy-D-Xylulose-5-Phosphate Synthase. Chembiochem 2015, 16 (12), 1771–1781. 10.1002/cbic.201500119.

(33) Ramsay, R. R.; Tipton, K. F. Assessment of Enzyme Inhibition: A Review with Examples from the Development of Monoamine Oxidase and Cholinesterase Inhibitory Drugs. Molecules 2017, 22 (7). 10.3390/molecules22071192.

(34) Strumilo, S.; Czerniecki, J.; Dobrzyn, P. Regulatory Effect of Thiamin Pyrophosphate on Pig Heart Pyruvate Dehydrogenase Complex. Biochem. Biophys. Res. Commun. 1999, 256 (2), 341–345. 10.1006/bbrc.1999.0321.

(35) Smith, J. M.; Vierling, R. J.; Meyers, C. F. Selective Inhibition of E. Coli 1-Deoxy-D-Xylulose-5-Phosphate Synthase by Acetylphosphonates. Medchemcomm 2012, 3, 65–67. 10.1039/C1MD00233C.

(36) Coco, L. B.; Toci, E. M.; Chen, P. Y.-T.; Drennan, C. L.; Freel Meyers, C. L. Potent Inhibition of E. Coli DXP Synthase by a Gem-Diaryl Bisubstrate Analog. ACS Infect Dis 2024. 10.1021/acsinfecdis.3c00734.

(37) Stephanie Henriquez, Charles R. Nosal, Joseph R. Knoff, Lauren B. Coco, Caren L. Freel Meyers. Bisubstrate Analog Inhibitors of DXP Synthase Show Species Specificity. Biochemistry. 10.1021/acs.biochem.4c00549.

(38) Alvarez, F. J.; Ermer, J.; Huebner, G.; Schellenberger, A.; Schowen, R. L. Catalytic Power of Pyruvate Decarboxylase. Rate-Limiting Events and Microscopic Rate Constants from Primary Carbon and Secondary Hydrogen Isotope Effects. J. Am. Chem. Soc. 1991, 113 (22), 8402–8409. 10.1021/ja00022a030.

(39) Coco, L. B.; Freel Meyers, C. L. An Activity-Based Probe for Antimicrobial Target DXP Synthase, a Thiamin Diphosphate-Dependent Enzyme. Front Chem Biol 2024, 3. 10.3389/fchbi.2024.1389620.

(40) Gounaris, A. D.; Turkenkopf, I.; Buckwald, S.; Young, A. Pyruvate Decarboxylase. J. Biol. Chem. 1971, 246 (5), 1302–1309. 10.1016/s0021-9258(19)76974-9.

(41) Johnston, M. L.; Toci, E. M.; DeColli, A. A.; Freel Meyers, C. L. Antibacterial Target DXP Synthase Catalyzes the Cleavage of D-Xylulose 5-Phosphate: A Study of Ketose Phosphate Binding and Ketol Transfer Reaction. Biochemistry 2022, 61 (17), 1810–1823. 10.1021/acs.biochem.2c00274.

(42) Altincicek, B.; Hintz, M.; Sanderbrand, S.; Wiesner, J.; Beck, E.; Jomaa, H. Tools for Discovery of Inhibitors of the 1-Deoxy-D-Xylulose 5-Phosphate (DXP) Synthase and DXP Reductoisomerase: An Approach with Enzymes from the Pathogenic Bacterium Pseudomonas Aeruginosa. FEMS Microbiol. Lett. 2000, 190 (2), 329–333. 10.1111/j.1574-6968.2000.tb09307.x.

(43) Bartee, D.; Freel Meyers, C. L. Targeting the Unique Mechanism of Bacterial 1-Deoxy-d-Xylulose-5-Phosphate Synthase. Biochemistry 2018, 57 (29), 4349–4356. 10.1021/acs.biochem.8b00548.

(44) Pinnataip, R.; Lee, B. P. Oxidation Chemistry of Catechol Utilized in Designing Stimuli-Responsive Adhesives and Antipathogenic Biomaterials. ACS Omega 2021, 6 (8), 5113–5118. 10.1021/acsomega.1c00006.

(45) Haemers, S.; Koper, G. J. M.; Frens, G. Effect of Oxidation Rate on Cross-Linking of Mussel Adhesive Proteins. Biomacromolecules 2003, 4 (4), 1098–1098. 10.1021/bm030048f.

(46) Maier, G. P.; Bernt, C. M.; Butler, A. Catechol Oxidation: Considerations in the Design of Wet Adhesive Materials. Biomater Sci 2018, 6 (2), 332–339. 10.1039/c7bm00884h.

(47) Sugumaran, M. Reactivities of Quinone Methides versus O-Quinones in Catecholamine Metabolism and Eumelanin Biosynthesis. Int. J. Mol. Sci. 2016, 17 (9), 1576. 10.3390/ijms17091576.

(48) Bolton, J. L.; Dunlap, T. Formation and Biological Targets of Quinones: Cytotoxic versus Cytoprotective Effects. Chem. Res. Toxicol. 2017, 30 (1), 13–37. 10.1021/acs.chemrestox.6b00256.

(49) Forooshani, P. K.; Meng, H.; Lee, B. P. Catechol Redox Reaction: Reactive Oxygen Species Generation, Regulation, and Biomedical Applications. In ACS Symposium Series; ACS symposium series. American Chemical Society; American Chemical Society: Washington, DC, 2017; pp 179–196. 10.1021/bk-2017-1252.ch010.

(50) Liehr, J. G.; Roy, D. Free Radical Generation by Redox Cycling of Estrogens. Free Radic. Biol. Med. 1990, 8 (4), 415–423. 10.1016/0891-5849(90)90108-u.

(51) Lopalco, A.; Dalwadi, G.; Niu, S.; Schowen, R. L.; Douglas, J.; Stella, V. J. Mechanism of Decarboxylation of Pyruvic Acid in the Presence of Hydrogen Peroxide. J. Pharm. Sci. 2016, 105 (2), 705–713. 10.1002/jps.24653.

(52) Drachman, N.; Kadlecek, S.; Duncan, I.; Rizi, R. Quantifying Reaction Kinetics of the Non-Enzymatic Decarboxylation of Pyruvate and Production of Peroxymonocarbonate with Hyperpolarized 13C-NMR. Phys. Chem. Chem. Phys. 2017, 19 (29), 19316–19325. 10.1039/c7cp02041d.

(53) Njus, D.; Asmaro, K.; Li, G.; Palomino, E. Redox Cycling of Quinones Reduced by Ascorbic Acid. Chem. Biol. Interact. 2023, 373, 110397. 10.1016/j.cbi.2023.110397.

(54) Alfonso-Prieto, M.; Biarnés, X.; Vidossich, P.; Rovira, C. The Molecular Mechanism of the Catalase Reaction. J. Am. Chem. Soc. 2009, 131 (33), 11751–11761. 10.1021/ja9018572.

(55) Kono, Y.; Fridovich, I. Superoxide Radical Inhibits Catalase. J. Biol. Chem. 1982, 257 (10), 5751–5754. 10.1016/s0021-9258(19)83842-5.

(56) Shimizu, N.; Kobayashi, K.; Hayashi, K. The Reaction of Superoxide Radical with Catalase. Mechanism of the Inhibition of Catalase by Superoxide Radical. J. Biol. Chem. 1984, 259 (7), 4414–4418. 10.1016/s0021-9258(17)43062-6.

(57) Qi, H.; Zhang, C. Simultaneous Determination of Hydroquinone and Catechol at a Glassy Carbon Electrode Modified with Multiwall Carbon Nanotubes. Electroanalysis 2005, 17 (10), 832–838. 10.1002/elan.200403150.

(58) Swain, C. G.; Unger, S. H.; Rosenquist, N. R.; Swain, M. S. Substituent Effects on Chemical Reactivity. Improved Evaluation of Field and Resonance Components. J. Am. Chem. Soc. 1983, 105 (3), 492–502. 10.1021/ja00341a032.

(59) Kang, D.-K.; Kim, S.-H.; Sohn, J.-H.; Sung, B. H. Insights into Enzyme Reactions with Redox Cofactors in Biological Conversion of CO2. J. Microbiol. Biotechnol. 2023, 33 (11), 1403–1411. 10.4014/jmb.2306.06005.

(60) Tittmann, K. Reaction Mechanisms of Thiamin Diphosphate Enzymes: Redox Reactions. FEBS J. 2009, 276 (9), 2454–2468. 10.1111/j.1742-4658.2009.06966.x.

(61) Tanner, M. E. Transient Oxidation as a Mechanistic Strategy in Enzymatic Catalysis. Curr. Opin. Chem. Biol. 2008, 12 (5), 532–538. 10.1016/j.cbpa.2008.06.016.

(62) Rodemeister, S.; Hill, K. Pyruvate Diminishes the Cytotoxic Activity of Ascorbic Acid in Several Tumor Cell Lines in Vitro. Biochem. Biophys. Res. Commun. 2017, 493 (3), 1184– 1189. 10.1016/j.bbrc.2017.09.138.

(63) Tittmann, K.; Wille, G.; Golbik, R.; Weidner, A.; Ghisla, S.; Hübner, G. Radical Phosphate Transfer Mechanism for the Thiamin Diphosphate-and FAD-Dependent Pyruvate Oxidase from Lactobacillus Plantarum. Kinetic Coupling of Intercofactor Electron Transfer with Phosphate Transfer to Acetyl-Thiamin Diphosphate via a Transient FAD Semiquinone/Hydroxyethyl-ThDP Radical Pair. Biochemistry 2005, 44 (40), 13291–13303. 10.1021/bi051058z.

(64) Mansoorabadi, S. O.; Seravalli, J.; Furdui, C.; Krymov, V.; Gerfen, G. J.; Begley, T. P.; Melnick, J.; Ragsdale, S. W.; Reed, G. H. EPR Spectroscopic and Computational Characterization of the Hydroxyethylidene-Thiamine Pyrophosphate Radical Intermediate of Pyruvate:Ferredoxin Oxidoreductase. Biochemistry 2006, 45 (23), 7122–7131. 10.1021/bi0602516.

(65) Pierce, E.; Mansoorabadi, S. O.; Can, M.; Reed, G. H.; Ragsdale, S. W. Properties of Intermediates in the Catalytic Cycle of Oxalate Oxidoreductase and Its Suicide Inactivation by Pyruvate. Biochemistry 2017, 56 (22), 2824–2835. 10.1021/acs.biochem.7b00222.

(66) Nemeria, N. S.; Gerfen, G.; Nareddy, P. R.; Yang, L.; Zhang, X.; Szostak, M.; Jordan, F. The Mitochondrial 2-Oxoadipate and 2-Oxoglutarate Dehydrogenase Complexes Share Their E2 and E3 Components for Their Function and Both Generate Reactive Oxygen Species. Free Radic. Biol. Med. 2018, 115, 136–145. 10.1016/j.freeradbiomed.2017.11.018.

(67) Tse, M. T.; Andschloss, J. V. Theoxygenasereactionofacetolactate Synthase. Biochemistry 1993, 32, 10398–10403.

(68) Gallardo-Garrido, C.; Cho, Y.; Cortés-Rios, J.; Vasquez, D.; Pessoa-Mahana, C. D.; Araya-Maturana, R.; Pessoa-Mahana, H.; Faundez, M. Nitrofuran Drugs beyond Redox Cycling: Evidence of Nitroreduction-Independent Cytotoxicity Mechanism. Toxicol. Appl. Pharmacol. 2020, 401 (115104), 115104. 10.1016/j.taap.2020.115104.

(69) McOsker, C. C.; Fitzpatrick, P. M. Nitrofurantoin: Mechanism of Action and Implications for Resistance Development in Common Uropathogens. J. Antimicrob. Chemother. 1994, 33 Suppl A, 23–30. 10.1093/jac/33.suppl_a.23.

(70) Pinhal, S.; Ropers, D.; Geiselmann, J.; de Jong, H. Acetate Metabolism and the Inhibition of Bacterial Growth by Acetate. J. Bacteriol. 2019, 201 (13). 10.1128/JB.00147-19.

(71) Millard, P.; Gosselin-Monplaisir, T.; Uttenweiler-Joseph, S.; Enjalbert, B. Acetate Is a Beneficial Nutrient for E. Coli at Low Glycolytic Flux. EMBO J. 2023, 42 (15), e113079. 10.15252/embj.2022113079.

(72) Johnston, M. L.; Freel Meyers, C. L. Revealing Donor Substrate-Dependent Mechanistic Control on DXPS, an Enzyme in Bacterial Central Metabolism. Biochemistry 2021, 60 (12), 929–939. 10.1021/acs.biochem.1c00019.

