## Supporting Information for "Trihydroxybenzaldoximes are redox cycling inhibitors of ThDP-dependent DXP synthase"

### Table of Contents

**Figure S1.** $K_i$ determination of **1** on DXPS homologs. ....S3
**Figure S2.** Inhibition of ADH and PDC at pH 7 and 8 by **1** ....S4
**Figure S3.** $K_i$ determination of **1** on *Ec*R478A, *Ec*D427A, *Ec*H431A, and *Ec*R99A ....S5
**Figure S4.** NMR detection of nonenzymatic decarboxylation of pyruvate ....S6
**Figure S5.** pH dependence of nonenzymatic decarboxylation of pyruvate ....S7
**Figure S6.** NMR detection of pyruvate consumption in HEPES pH 8....S7
**Figure S7.** NMR detection of pyruvate consumption in Tris pH 8 ....S8
**Figure S8.** Mechanism of nonenzymatic oxidative decarboxylation of pyruvate by ROS....S9
**Figure S9.** pH dependence of autoxidation of 1....S10
**Figure S10.** NMR detection of pyruvate consumption in the presence of ascorbate ....S11
**Figure S11.** Mechanism of redox cycling of **1** with ascorbate.... S11
**Figure S12.** NMR detection of pyruvate consumption in the presence of catalase.... S12
**Figure S13.** Pre-saturation $^1$H-NMR of > 10 mM 1 in HEPES pH 8 .... S13
**Figure S14.** Anaerobic $K_i$ determination of 1.... S14
**Figure S15.** $K_i$ determination of 3 and 4 .... S14
**Figure S16.** Cyclic Voltammograms of catechol SAR .... S15
**Figure S17.** Long Range $^1$H-$^{15}$N HSQC NMR of 8.... S16
**Figure S18.** Experimental scheme for detection of reduction of 1 by DXPS .... S17
**Figure S19.** Long Range $^1$H-$^{15}$N HSQC NMR of 8 with ascorbate .... S18
**Figure S20.** Long Range $^1$H-$^{15}$N HSQC NMR of 8 with DXPS .... S18
**Figure S21.** Long Range $^1$H-$^{15}$N HSQC NMR of 8 with Pyruvate and D-GAP.... S19
**Figure S22.** Long Range $^1$H-$^{15}$N HSQC NMR of 8 with DXPS, Pyruvate and D-GAP .... S20
**Figure S23.** Long Range $^1$H-$^{15}$N HSQC NMR of 8 with DXPS and Pyruvate .... S21
**Figure S24.** NMR detection of DXPS reaction products.... S22
**Figure S25.** Mechanism of acetate formation from reduction of 1 .... S23
**Figure S26.** Raw amperoteric i-t measurements of reactions with 1.... S23

**Figure S27.** Color change of oxime 1 based on redox form ........ S24
**Figure S28.** Full CVs of 8 measured on gold UME ........ S24
**Figure S29.** $^{1}$H-NMR of 3 ........ S25
**Figure S30.** $^{13}$C-NMR of 3 ........ S26
**Figure S31.** HRMS detection of $3^{R}$ ........ S27
**Figure S32.** HRMS detection of $3^{O}$ ........ S28
**Figure S33.** HPLC of 3 ........ S29
**Figure S34.** $^{1}$H-NMR of 8 ........ S30
**Figure S35.** $^{13}$C-NMR of 8 ........ S31
**Figure S36.** HRMS detection of $8^{R}$ ........ S32
**Figure S37.** HRMS detection of $8^{O}$ ........ S33
**Figure S38.** HPLC of 8 ........ S34

**Table S1.** Summary of apparent $K_i$s of 1 ........ S35
**Table S2.** Summary of apparent $K_i$s of 1, 2, 3, and 4 ........ S35
**References Cited** ........ S36

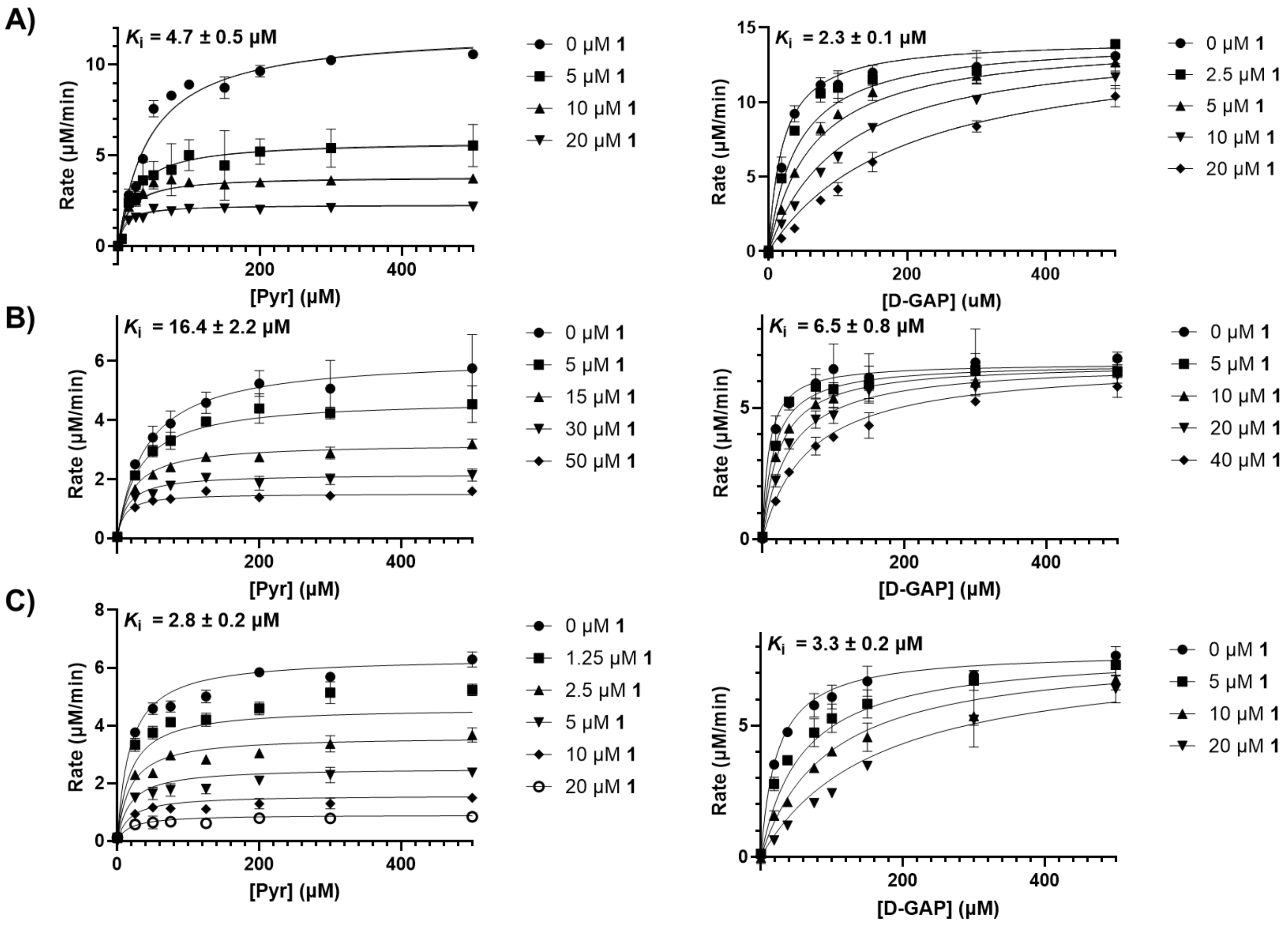


**Figure S1. Determination of $K_i$ for 1 against DXPS homologs.** Determination of apparent $K_i$ for **1**, varying pyruvate (left, uncompetitive) or D-GAP (right, competitive) on **A**) *Ec*DXPS, **B**) *Pa*DXPS, and **C**) *Dr*DXPS. Error bars represent standard deviation from 3 replicates at each concentration. Standard error of the mean for reported $K_i$ values was determined from 3 replicates.

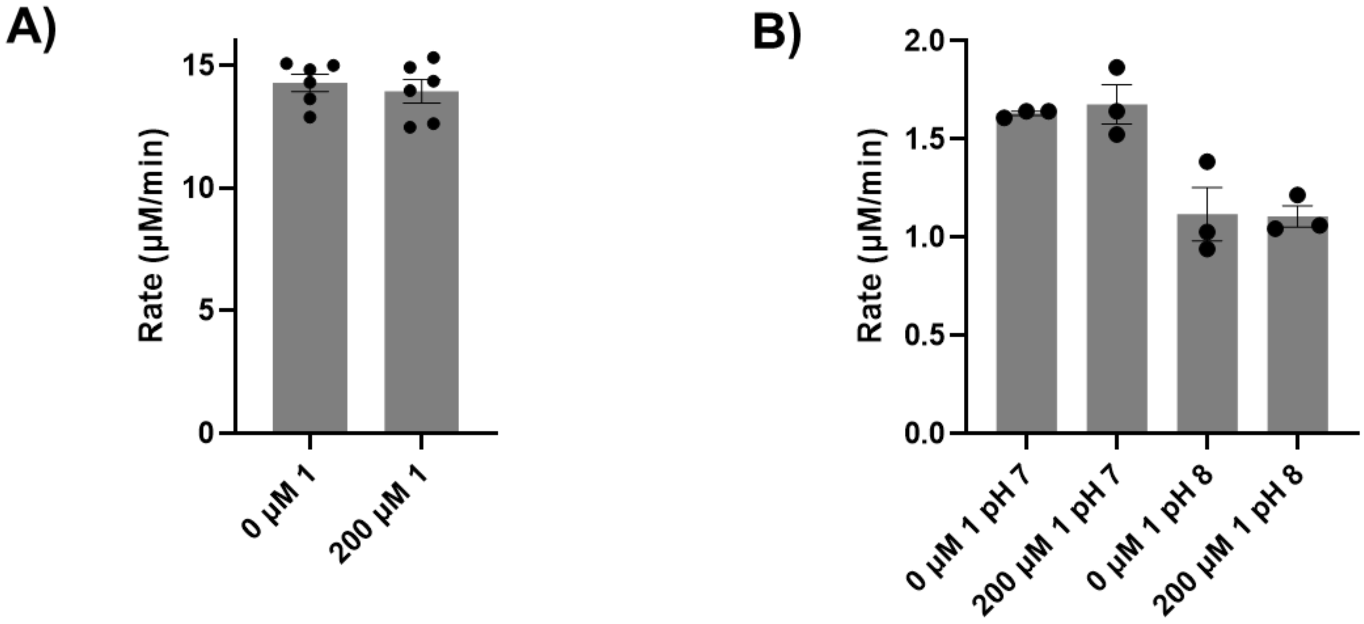


**Figure S2. Evaluation of inhibitory activity of 1 in the PDC-ADH coupled system. A)** ADH was not inhibited significantly by **1** up to a concentration of 200 μM, the highest concentration that does not significantly interfere with absorbance measurements in the coupled assay. Initial rates were arithmetically corrected for background (no substrates). Standard error of the mean was calculated from six replicates **B)** Inhibition of PDC by **1** (0 and 200 μM) is not observed at pH 7 or 8. Standard error of the mean was calculated from three replicates.

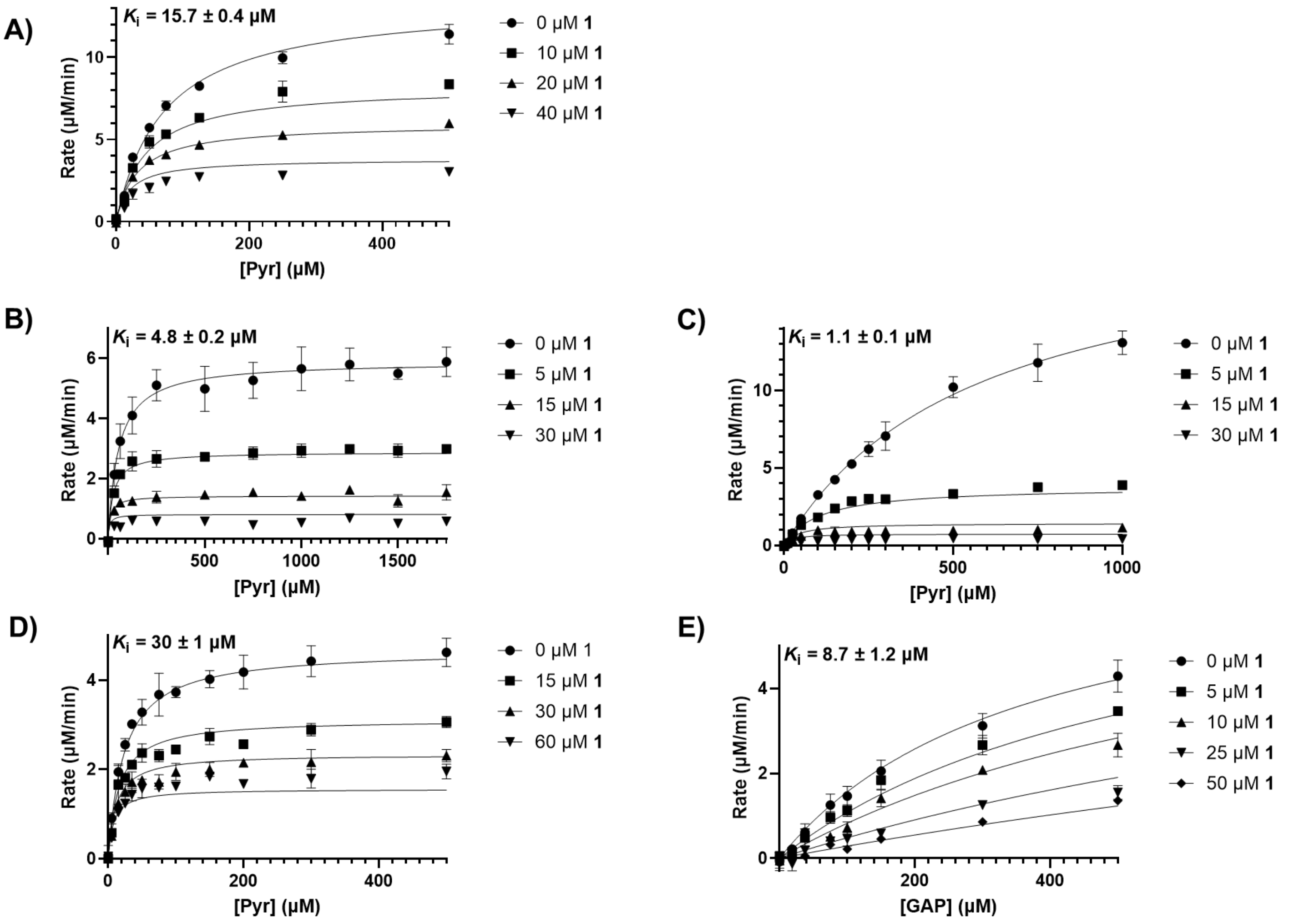


**Figure S3. $K_i$ determination of 1 on *Ec*R478A, *Ec*D427A, *Ec*H431A, and *Ec*R99A**. Determination of apparent $K_i$ of **1** under conditions of varying pyruvate (uncompetitive) on **A**) *Ec*R478A, **B**) *Ec*D427A, **C**) *Ec*H431A, and **D**) *Ec*R99A. **E**) Apparent $K_i$ determination of **1** varying D-GAP (competitive) on *Ec*R99A. Standard error of the mean for reported Ki values was determined from 3 replicates.

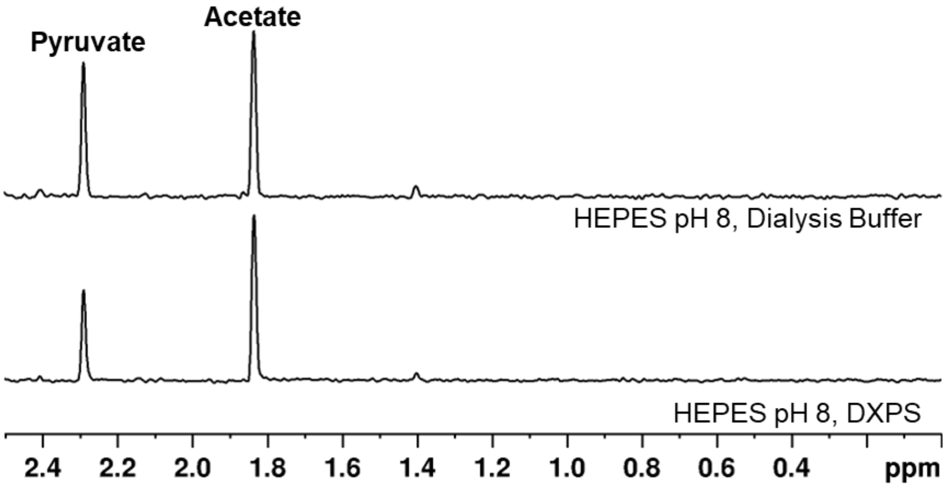


**Figure S4**. **NMR detection of nonenzymatic decarboxylation of pyruvate**. In the anaerobic chamber, reaction mixtures containing HEPES (50 mM, pH 8), $MgCl_2$ (2 mM), NaCl (100 mM), ThDP (1 mM), $D_2O$ (10%), and either DXPS (200 nM) or an equivalent volume of DXPS dialysis buffer were incubated with sodium pyruvate-$^{13}C_3$ for 5 min at 4°C. Reactions were initiated via addition of **1** (500 µM) and incubated for 15 min before quenching by boiling at 4°C for 5 min followed by vortex. Samples were removed from the chamber and filtered using 10 kDa cutoff microcentrifuge filters via centrifugation at 13000 rpm for 10 minutes at 4°C. Gadobutrol (250 µM) was added post centrifugation. 1D $^1H$ detected $H\{^{13}C\}$ and $H_3\{^{13}C\}$ spectra via a proton multiplicity filter were acquired (Bruker Avance 600 MHz). Significant pyruvate consumption detected as the evolution of acetate was observed in the absence of DXPS, suggesting an alternative reaction occurred.

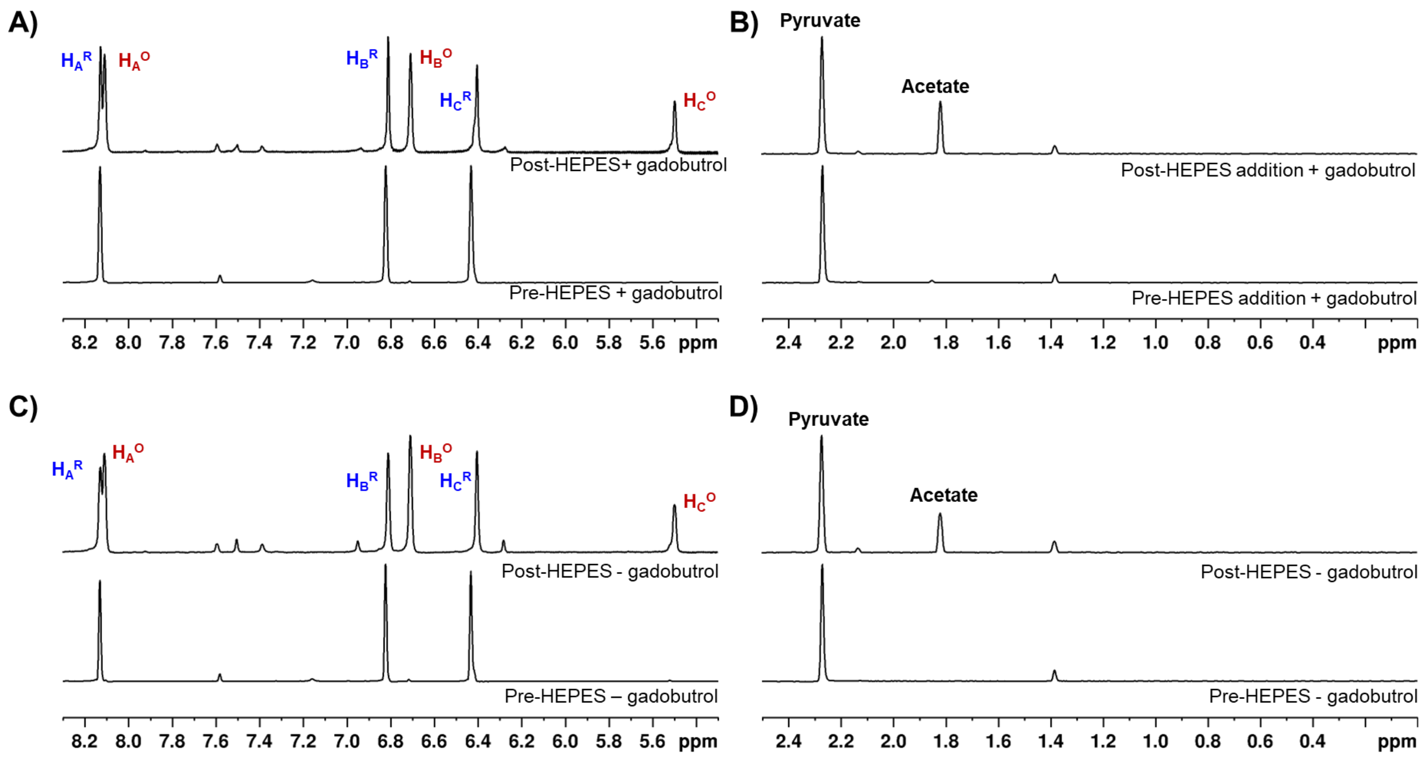


**Figure S5. Detection of pH dependence of nonenzymatic decarboxylation of pyruvate**. Reaction mixtures containing $D_2O$ (10%), sodium pyruvate-$^{13}C_3$, and either gadobutrol (250 μM) (**A**, **B**) or an equivalent volume of water (**C**, **D**) were incubated for 5 min at 4°C. Reactions were initiated via addition of **1** (500 μM) and incubated or 15 min before transfer to NMR tubes. 1D $^1$H-NMR and 1D $^1$H detected H{$^{13}$C} and $H_3${$^{13}$C} spectra via a proton multiplicity filter were acquired (Bruker Avance 600 MHz). HEPES (3 mM, pH 8.2) was then added to NMR tubes and spectra were acquired again. Nonenzymatic decarboxylation of pyruvate was observed as the formation of acetate post-addition of HEPES (**B**, **D**) regardless of gadobutrol. Additionally, changes in $^1$H-NMR spectra of **1** indicate oxidation occurred after addition of HEPES (**A**, **C**).

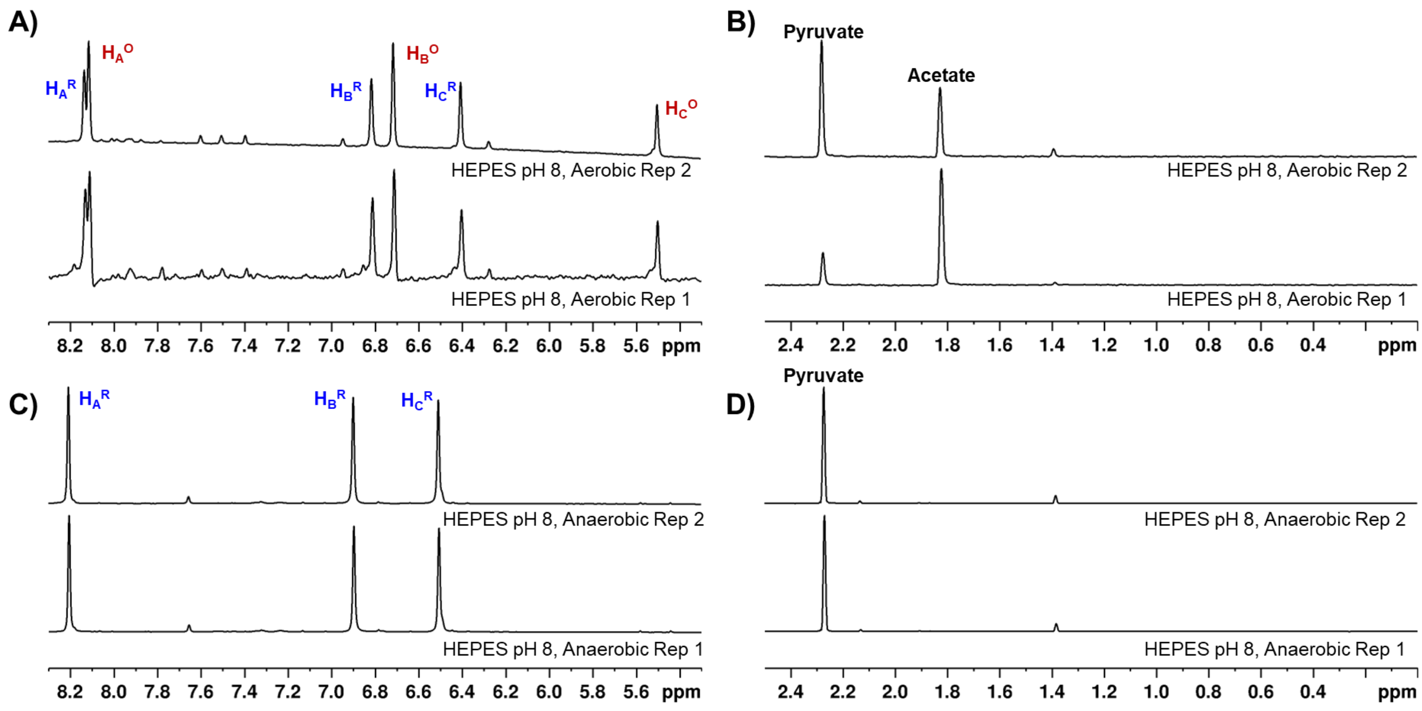


**Figure S6. NMR detection of pyruvate consumption in HEPES pH 8**. Oxime **1** (500 µM) was incubated with sodium pyruvate-$^{13}C_3$ (500 µM) in the presence of HEPES buffer (50 mM, pH 8) either aerobically (**A**, **B**) or anaerobically (**C**, **D**). Oxidation state of **1** (**A**, **C,** Bruker Avance, 600 MHz) and consumption of pyruvate (**B**, **D**, Bruker Avance 600 MHz) was determined by $^1$H-NMR (**A**, **C**) or 1D $^1$H detected H{$^{13}$C} and $H_3${$^{13}$C} spectra via a proton multiplicity filter (**B**, **D**) as described in the Materials and Methods section. Oxime **1** was partially oxidized in HEPES pH 8 under aerobic conditions but not in the absence of $O_2$.

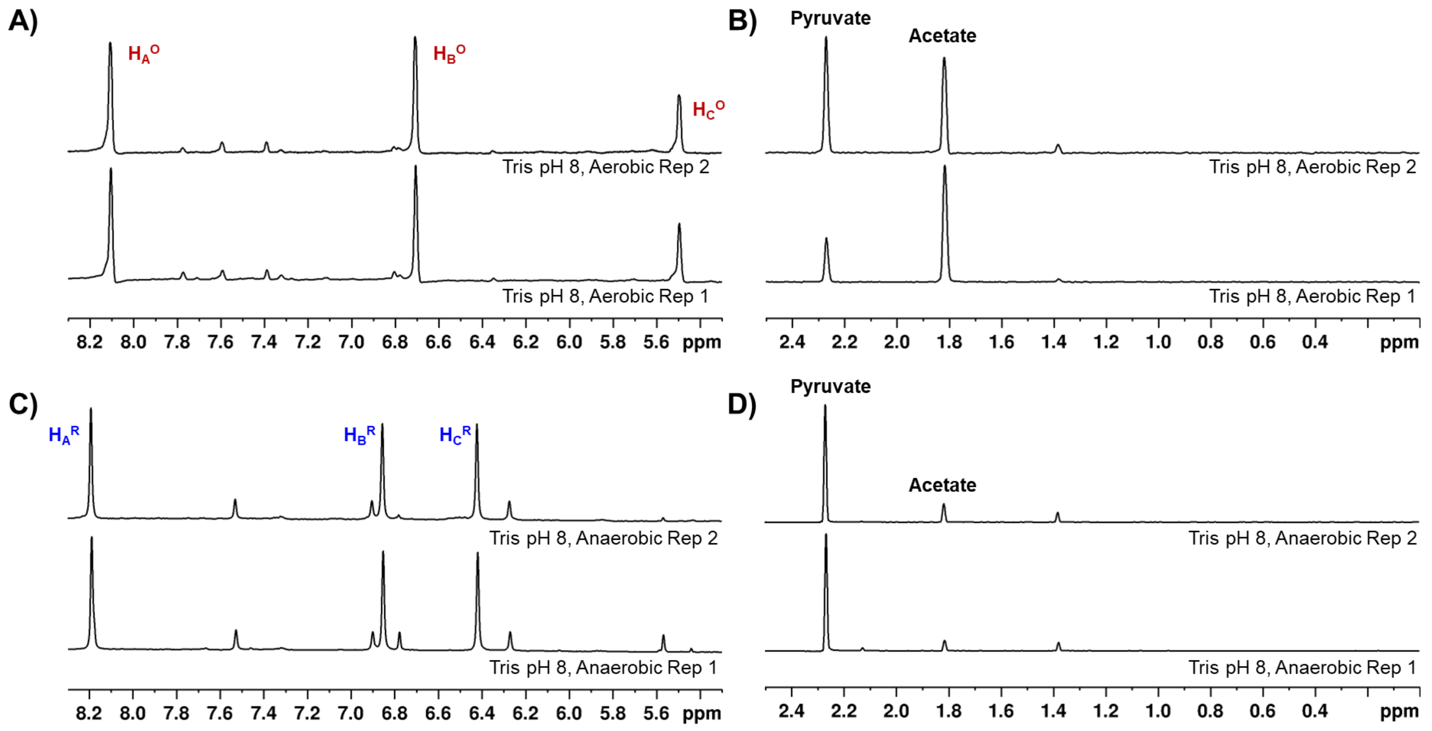


**Figure S7. NMR detection of pyruvate consumption in Tris pH 8**. Oxime **1** (500 µM) was incubated with sodium pyruvate-$^{13}C_3$ (500 µM) in the presence of Tris buffer (50 mM, pH 8) either aerobically (**A**, **B**) or anaerobically (**C**, **D**). Oxidation state of **1** (**A**, **C,** Bruker Avance, 600 MHz) and consumption of pyruvate (**B**, **D**, Bruker Avance 600 MHz) was determined by $^1$H-NMR (**A**, **C**) or 1D $^1$H detected H{$^{13}$C} and $H_3${$^{13}$C} spectra via a proton multiplicity filter (**B**, **D**) as described in the Materials and Methods section. Oxime **1** was mostly oxidized in Tris pH 8 under aerobic conditions but not in the absence of $O_2$.

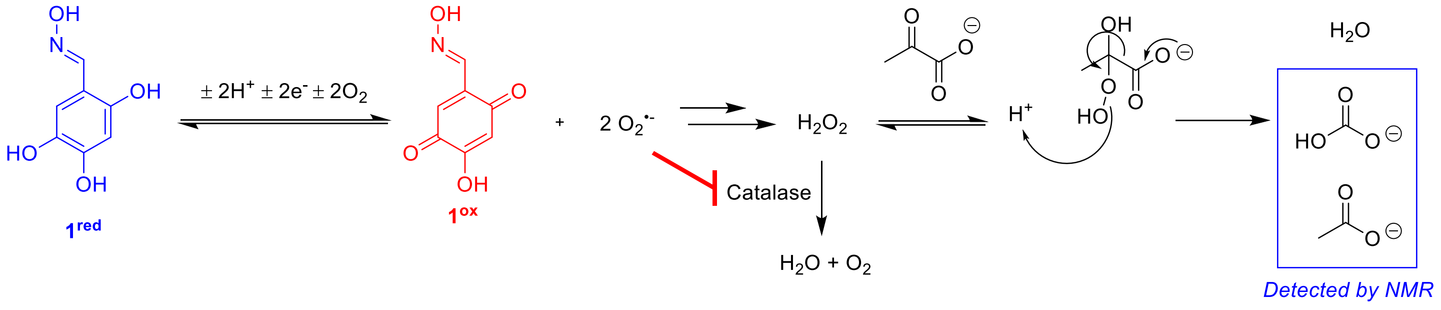


**Figure S8. Mechanism of nonenzymatic oxidative decarboxylation of pyruvate by ROS.** Oxime **1** is oxidized under alkaline aerobic conditions, generating ROS superoxide in the process. Superoxide reacts to form $H_2O_2$ which further reacts with pyruvate to induce oxidative decarboxylation, resulting in the end products acetate and bicarbonate we observe by NMR. Catalase (CAT) catalyzes the reduction of $H_2O_2$ to form water and $O_2$ which in turn oxidizes **1**. Superoxide is also capable of inhibiting catalase, so an excess of CAT was used to prevent loss of ROS scrubbing activity.

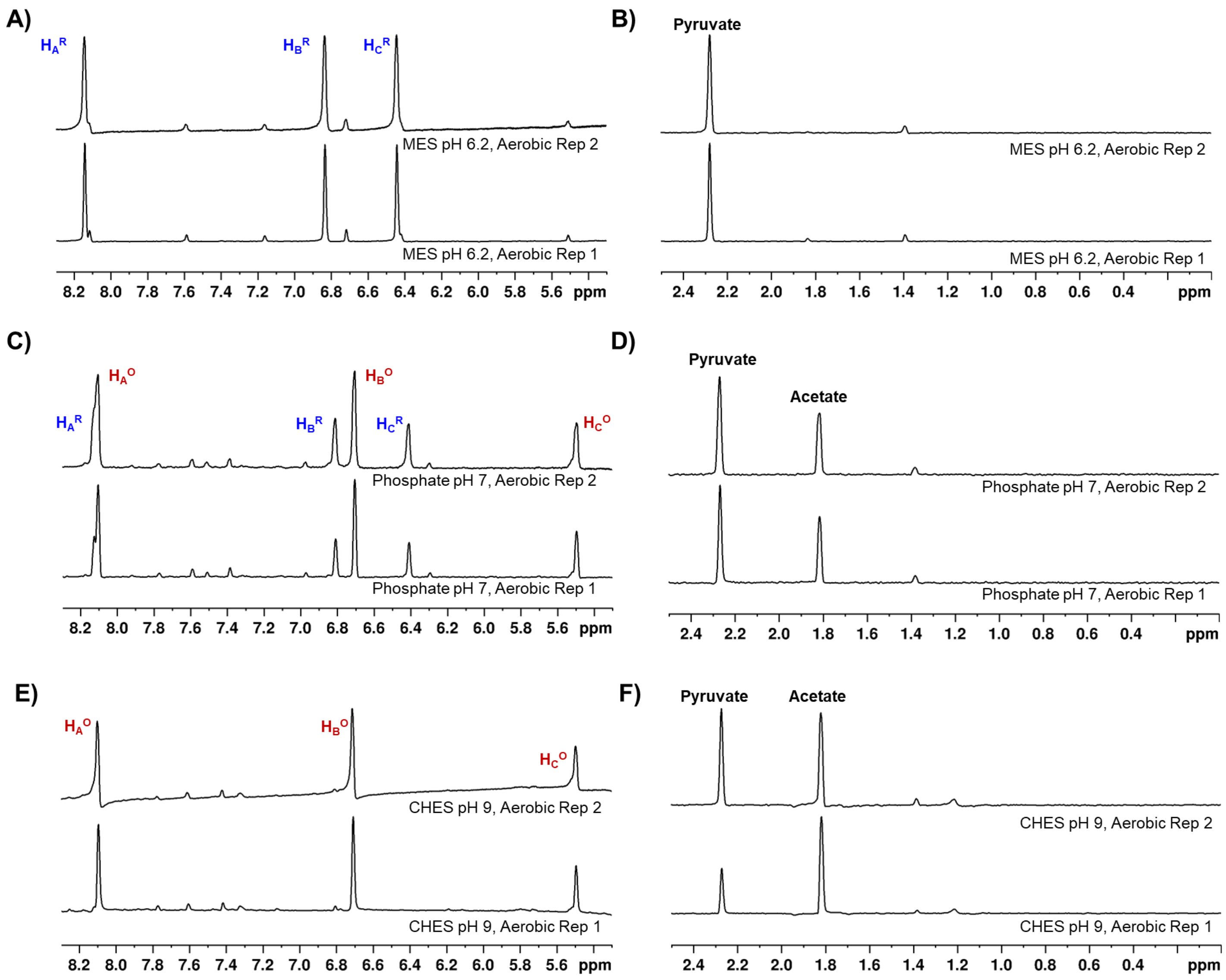


**Figure S9. pH dependence of autoxidation of 1.** Oxime **1** (500 µM) was incubated aerobically with sodium pyruvate-$^{13}C_3$ (500 µM) in the presence of MES buffer (50 mM, pH 6.2) (**A**, **B**), phosphate buffer (50 mM, pH 7) (**C**, **D**), or CHES buffer (50 mM, pH 9) (**E**, **F**). Oxidation state of **1** (**A**, **C, E**, Bruker Avance, 600 MHz) and consumption of pyruvate (**B**, **D**, **F**, Bruker Avance 600 MHz) was determined by $^1$H-NMR (**A**, **C**, **E**) or 1D $^1$H detected H{$^{13}$C} and $H_3${$^{13}$C} spectra via a proton multiplicity filter (**B**, **D**, **F**) as described in the Materials and Methods section. Oxime **1** was oxidized at neutral and basic pH, and coinciding pyruvate consumption was observed. At acidic pH, **1** was mostly reduced and no acetate formation was detected.

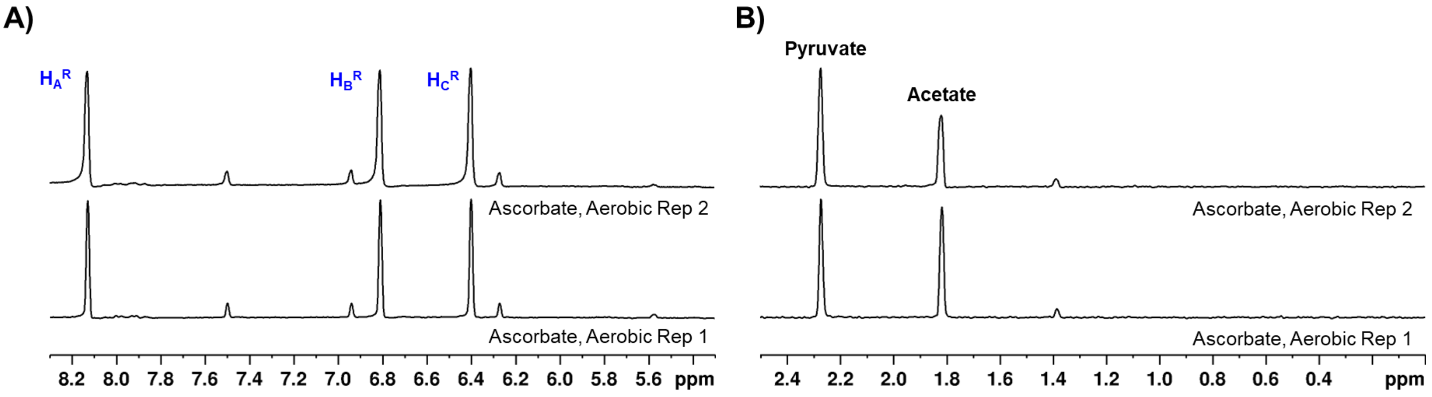


**Figure S10. NMR detection of pyruvate consumption in the presence of ascorbate**. Oxime **1** (500 µM) was incubated aerobically with sodium pyruvate-$^{13}C_3$ (500 µM) in the presence of HEPES buffer (50 mM, pH 8) and ascorbate (5 mM). Oxidation state of **1** (**A**, Bruker Avance, 600 MHz) and consumption of pyruvate (**B**, Bruker Avance 600 MHz) was determined by $^1$H-NMR (**A**) or 1D $^1$H detected H{$^{13}$C} and $H_3${$^{13}$C} spectra via a proton multiplicity filter (**B**) as described in the Materials and Methods section.. Oxime **1** was reduced in the presence of ascorbate but nonenzymatic consumption of pyruvate still occurred.

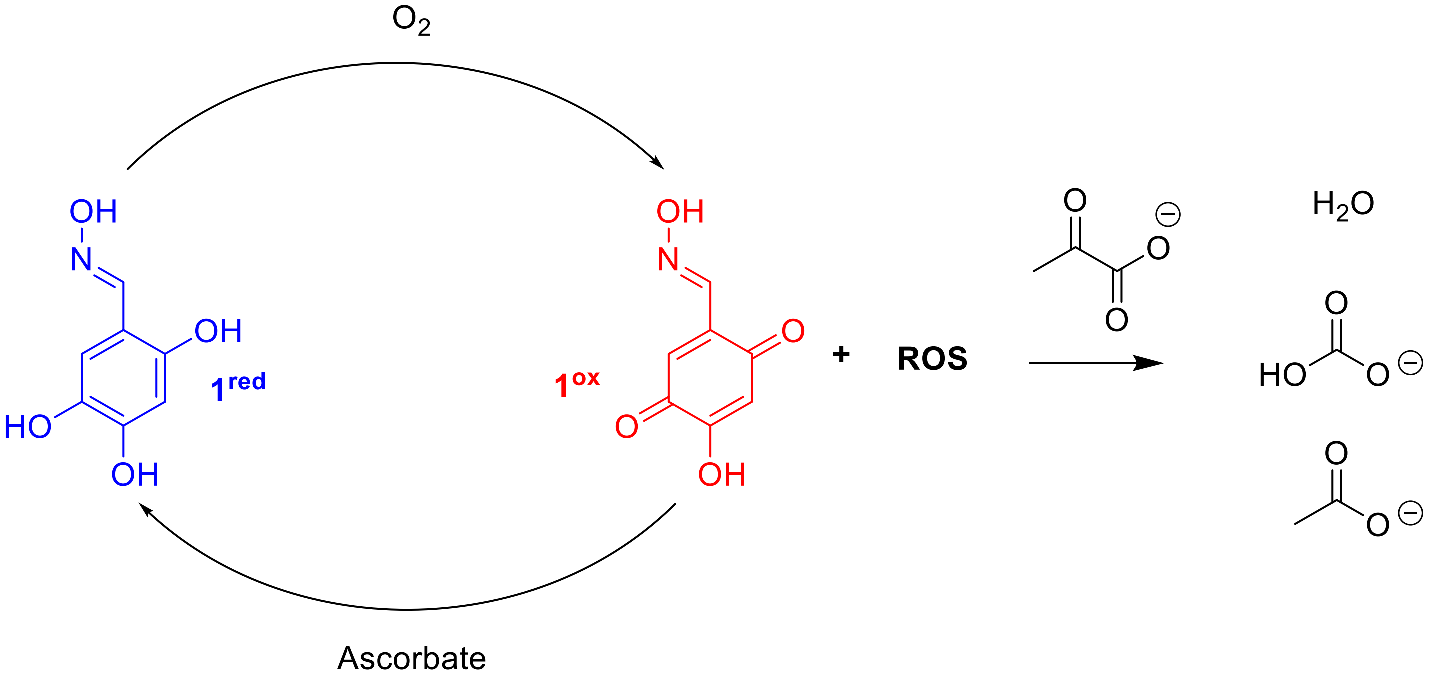


**Figure S11. Mechanism of redox cycling of 1 in the presence of ascorbate**. Ascorbate exacerbates the formation of ROS from the autoxidation of **1** by reducing the oxidized form of **1** to initiate a redox cycle in which ROS are continuously generated while there is a supply of oxidant ($O_2$) and reductant (ascorbate).

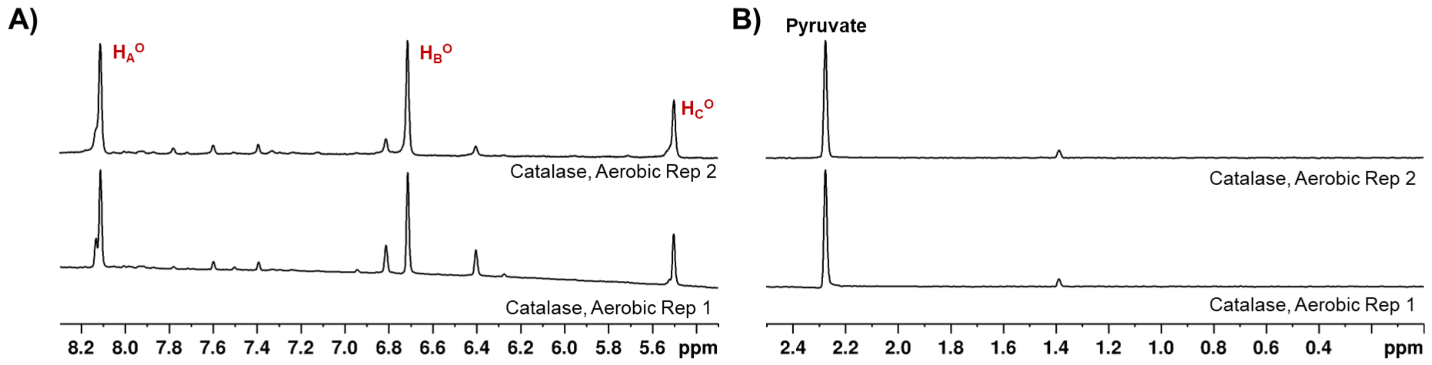


**Figure S12. NMR detection of pyruvate consumption in the presence of catalase.** Oxime **1** (500 μM) was incubated aerobically with sodium pyruvate-$^{13}C_3$ (500 μM) in the presence of HEPES buffer (50 mM, pH 8) and catalase (0.25 mg/mL). Oxidation state of **1** (**A**, Bruker Avance, 600 MHz) and consumption of pyruvate (**B**, Bruker Avance 600 MHz) was determined by $^1$H-NMR (**A**) or 1D $^1$H detected H{$^{13}$C} and $H_3${$^{13}$C} spectra via a proton multiplicity filter (**B**) as described in the Materials and Methods section. Oxime **1** was mostly oxidized in the presence of catalase and nonenzymatic consumption of pyruvate was not observed.

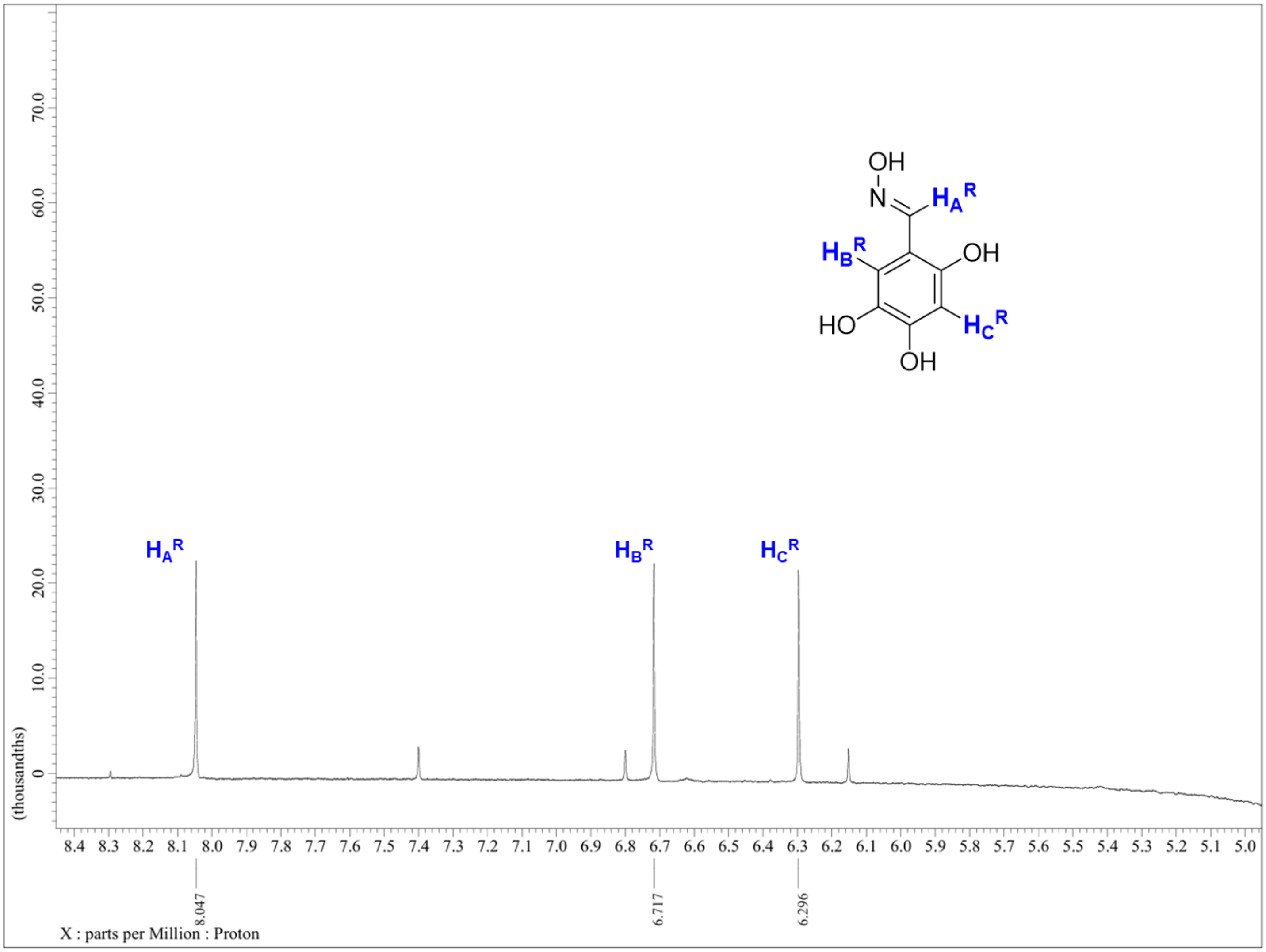


**Figure S13. Pre-saturation $^{1}$H-NMR of > 10 mM 1 in HEPES pH 8**. Oxime **1** (12.5 mM) was prepared aerobically in HEPES (100 mM, pH8) and $D_2O$ (10%). $^{1}$H-NMR (JEOL JNM-ECZL500R 500 MHz) was collected using a pre-saturation method to suppress the water peak. Oxime **1** was primarily in the reduced state at this concentration in contrast with other NMR experiments performed at sub-mM concentrations.

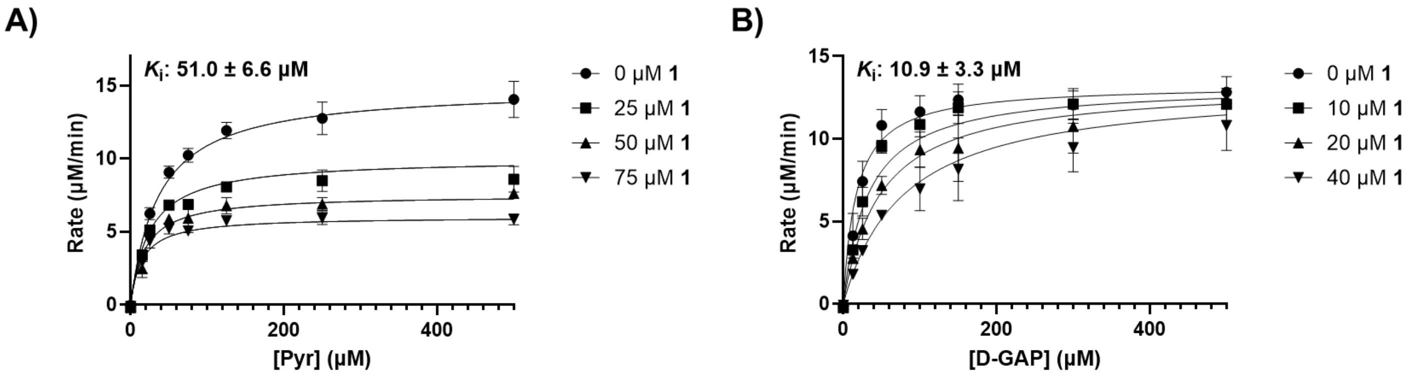


**Figure S14. $K_i$ determination of 1 under anaerobic conditions.** Determination of apparent $K_i$ of **1** under anaerobic conditions varying pyruvate (**A**, uncompetitive) and D-GAP (**B**, competitive) on WT *Ec*DXPS is shown. Standard error of the mean for reported $K_i$ values was determined from 3 replicates

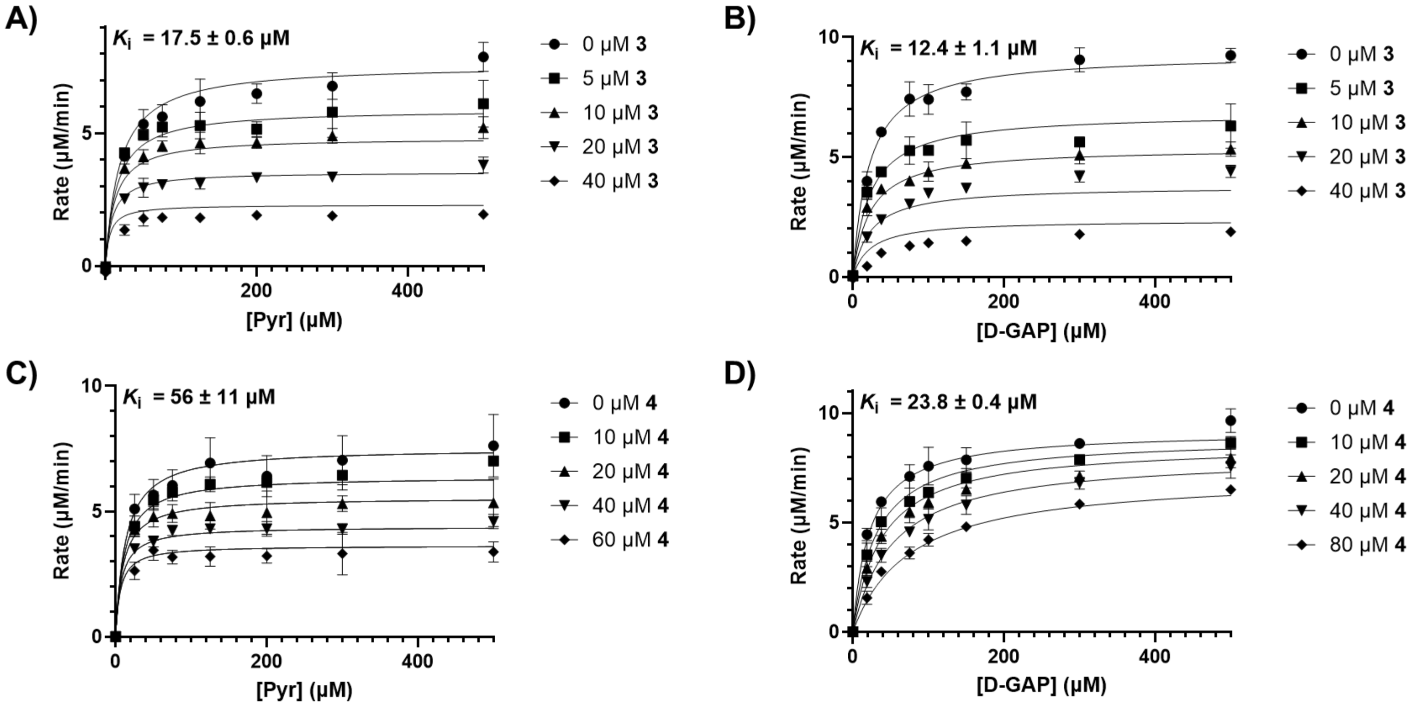


**Figure S15. $K_i$ determination of 3 and 4.** Determination of apparent $K_i$ of **3** (**A**, **B**) and **4** (**C**, **D**) varying pyruvate (**A**, **C**, uncompetitive) or D-GAP (**B**, competitive; **D** mixed) on WT *Ec*DXPS is shown. Initial rates in experiments with compound **4** were corrected arithmetically for background. Standard error of the mean for reported $K_i$ values was determined from 3 replicates

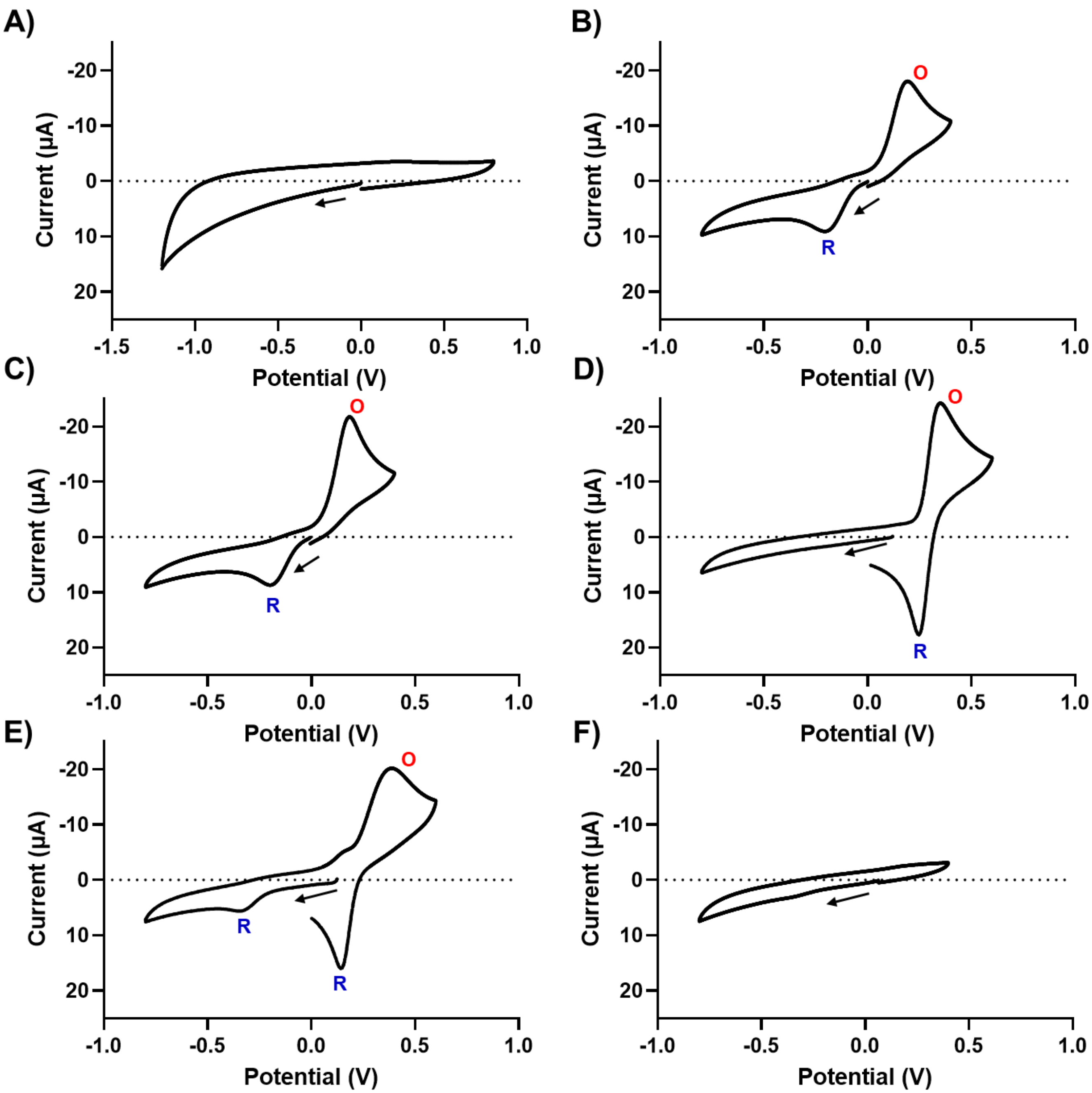


G)

| Compound | Oxidation (V) | Reduction (V) |
|---|---|---|
| **1** | 0.2 | -0.2 |
| **2** | 0.18 | -0.2 |
| **5** | 0.35 | 0.24 |
| **6** | 0.39 | -0.34, 0.14 |
| **7** | ND | ND |

**Figure S16. Cyclic Voltammograms of catechol SAR.** CVs of buffer (**A**), **1** (**B**), **2** (**C**), **5** (**D**), **6** (**E**), and **7** (**F**) are shown. Oxidation (**O**) and reduction (**R**) peaks are marked for clarity. The start and direction of voltage scanning are indicated by a black arrow. **G**) Oxidation and reduction potentials of **1**, **2**, **5**, **6** and **7**.

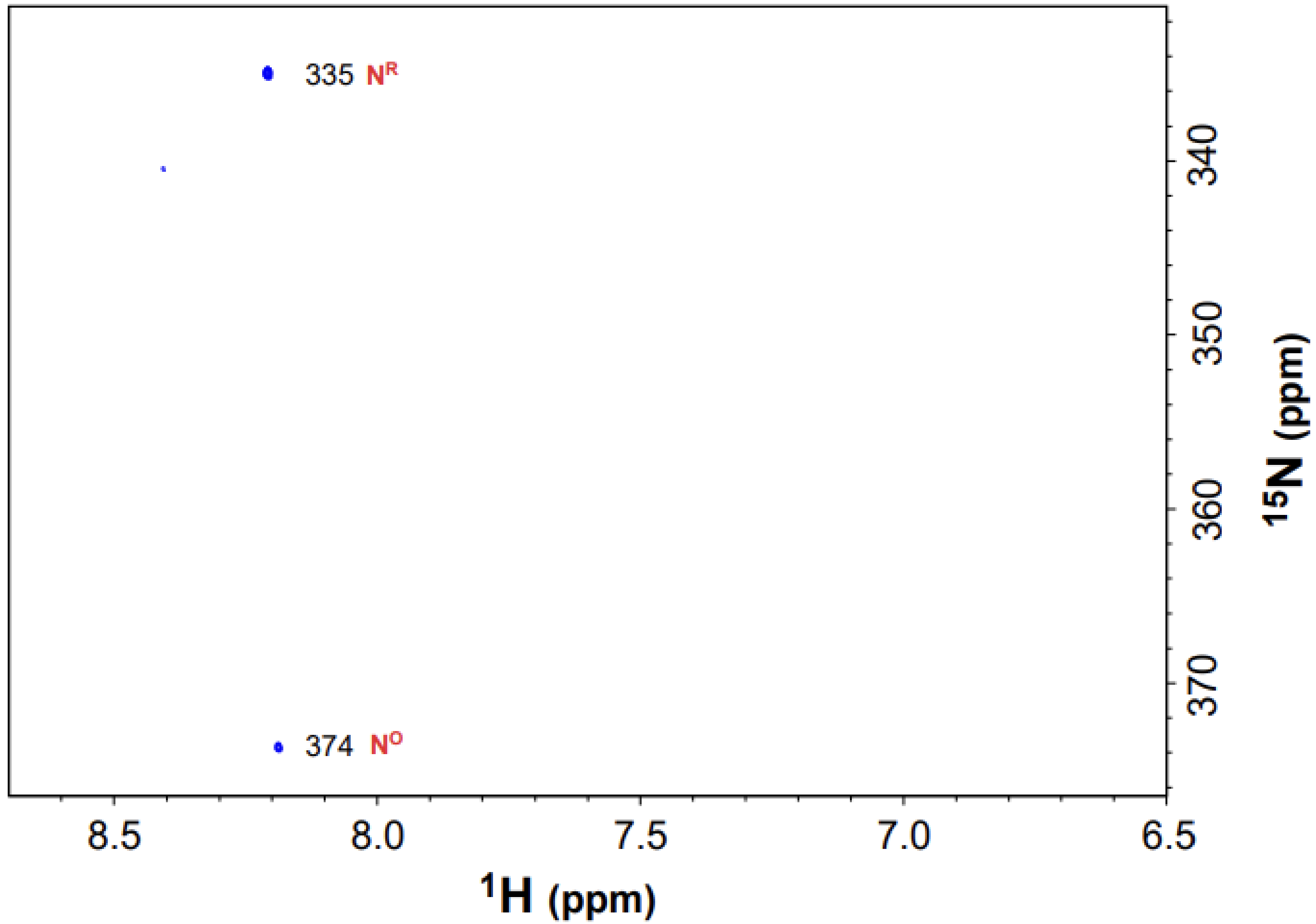


**Figure S17. Long Range $^1$H-$^{15}$N HSQC NMR of 8**. The chemical shift of $^{15}$N detected in the indirect dimension varied by ~40 ppm depending on redox state of **8**. Moving forward, the spectral width along $^{15}$N has been reduced for optimized data acquisition, resulting in the folding of the $N^R$ and $N^O$ peaks along the $^{15}$N dimension (Fig 8, S19-23). Here, the actual $^{15}$N chemical shifts for $N^R$ (335 ppm) and $N^O$ (374 ppm) are shown.

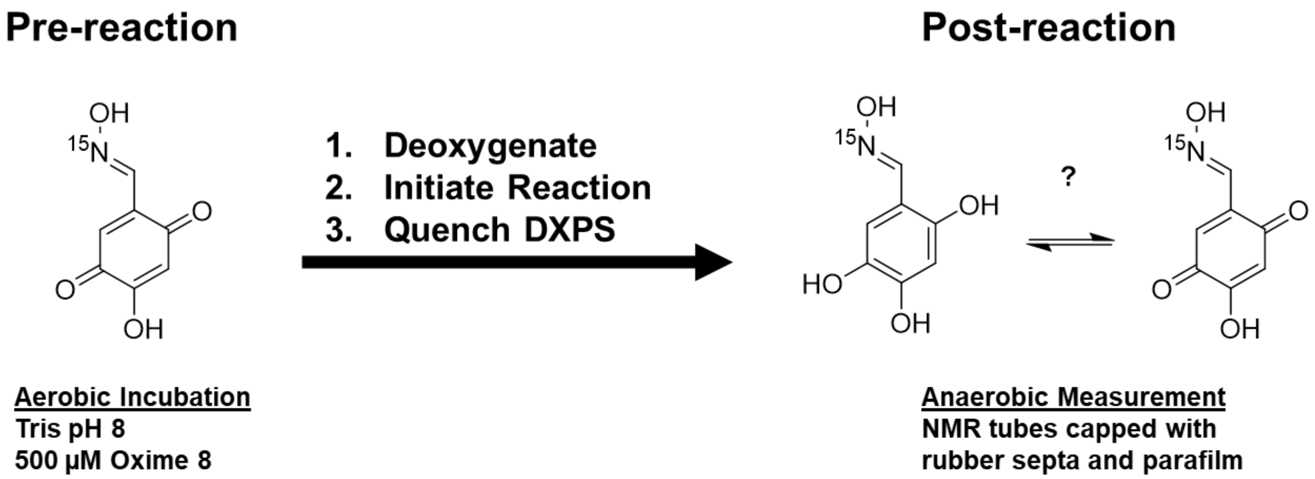


**Figure S18. Experimental design to detect DXPS-catalyzed reduction of 1**. In this experiment, **8** was preincubated in Tris (50 mM, pH 8) under aerobic conditions to promote the oxidized form. Solutions were then brought into the anaerobic chamber to deoxygenate. Reactions were then initiated and mixtures incubated at 25 °C for 1 hour before being quenched by boiling at 95 °C for 5 min and vortexing. Reactions were then transferred to NMR tubes, capped with septa, and wrapped with parafilm to exclude oxygen during spectra acquisition.

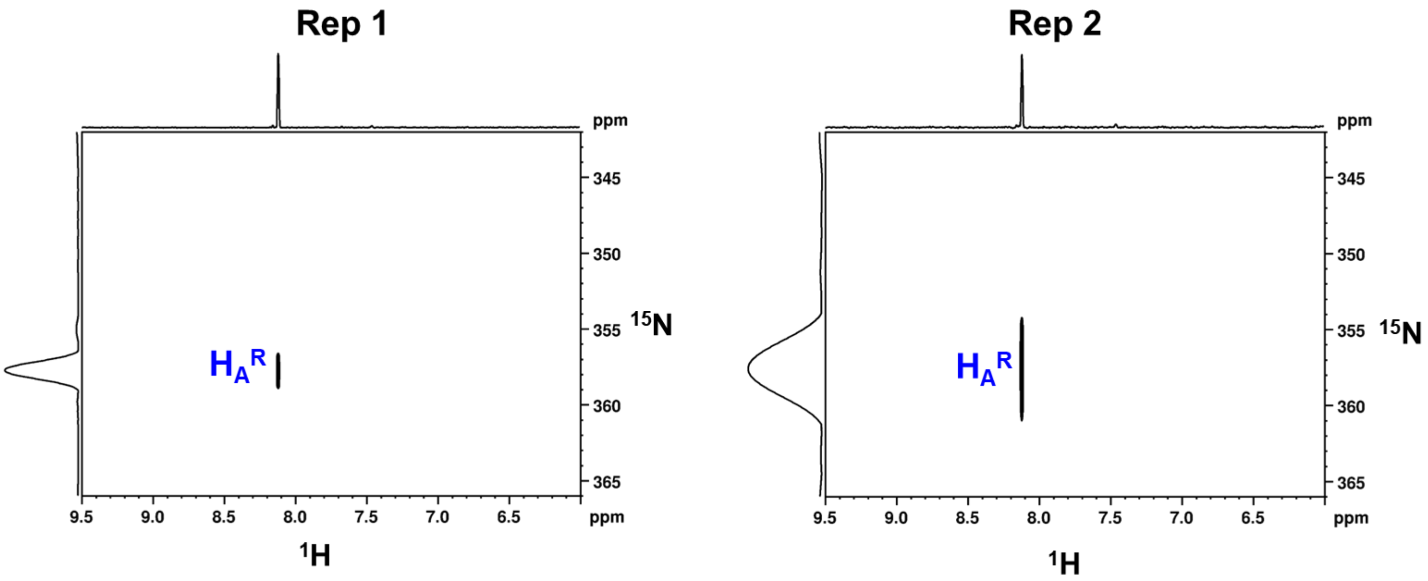


**Figure S19. Long Range $^{1}H$-$^{15}N$ HSQC NMR of 8 incubated with ascorbate**. Oxime **8** (500 µM) was incubated (as described in the Materials and Methods section) with ascorbate (5 mM) to verify the experimental workflow could be used to detect reduction of **8**. Total reduction was seen in the presence of ascorbate.

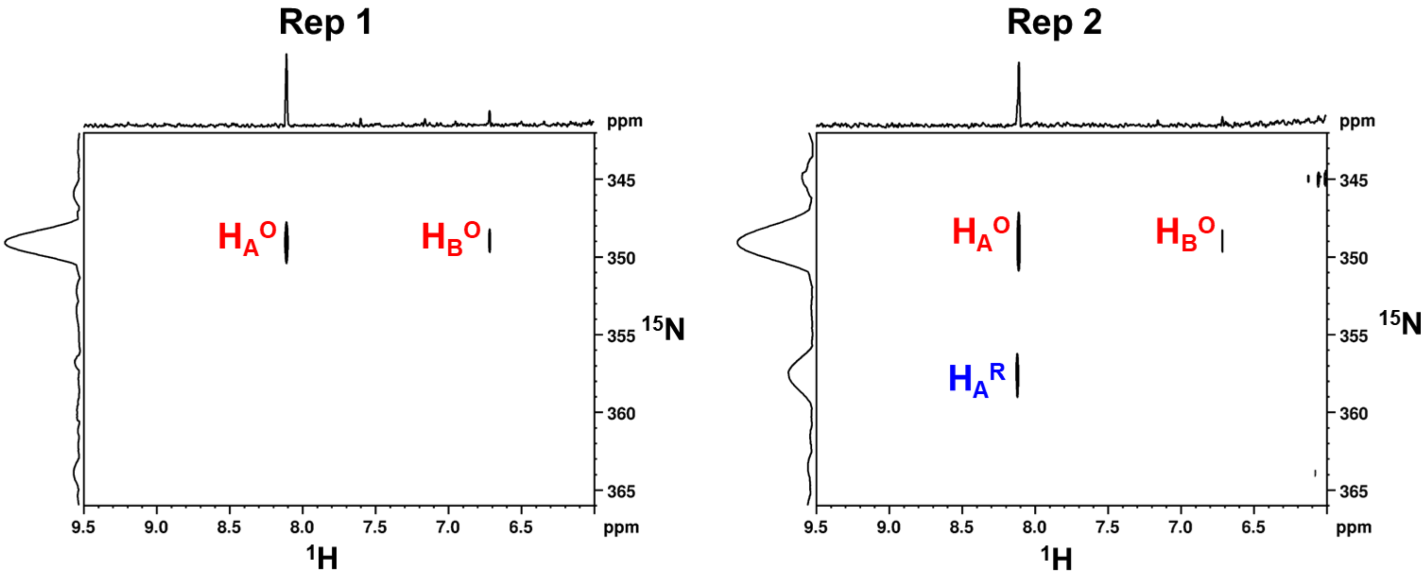


**Figure S20. Long Range $^{1}H$-$^{15}N$ HSQC NMR of 8 incubated with DXPS**. Oxime **8** (500 µM) was incubated (as described in the Materials and Methods section) with DXPS (1 µM) to verify the dependency of substrates on reduction of **8**. Oxime **8** was maintained mostly in the oxidized form in accordance with predictions.

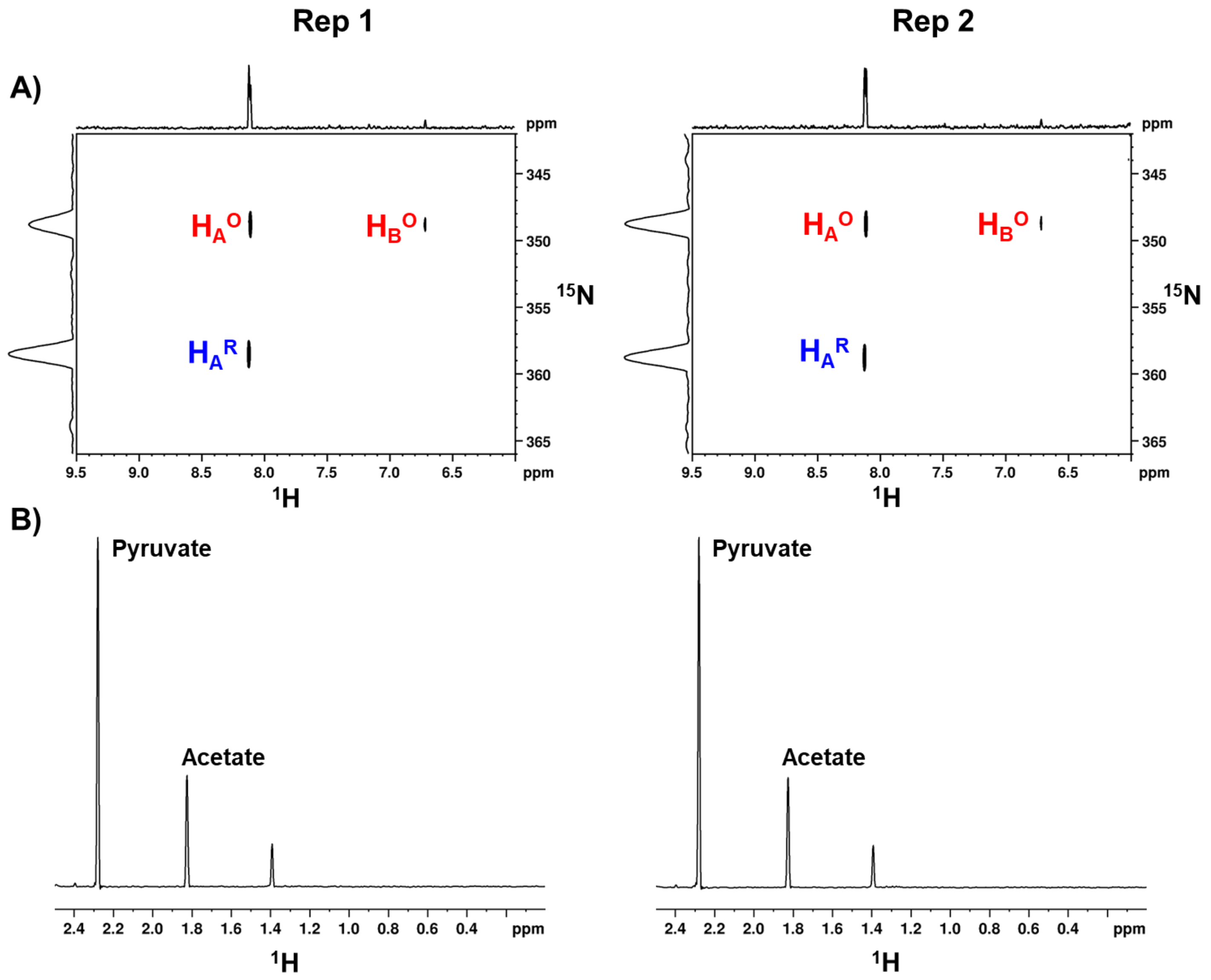


**Figure S21. Long Range $^{1}H$-$^{15}N$ HSQC NMR of 8 incubated with pyruvate and D-GAP**. Oxime **8** (500 µM) was incubated (as described in the Materials and Methods section) with pyruvate (2 mM) and D-GAP (500 µM) to verify the dependence of DXPS in the reduction of **8**. Some reduction of **8** (**A**) was observed via LR $^{1}H$-$^{15}N$ HSQC concurrent with acetate formation in the 1D H{$^{13}C$} and $H_3${$^{13}C$} filtered experiment (**B**) which could result from residual ROS produced in solution during the initial aerobic incubation.

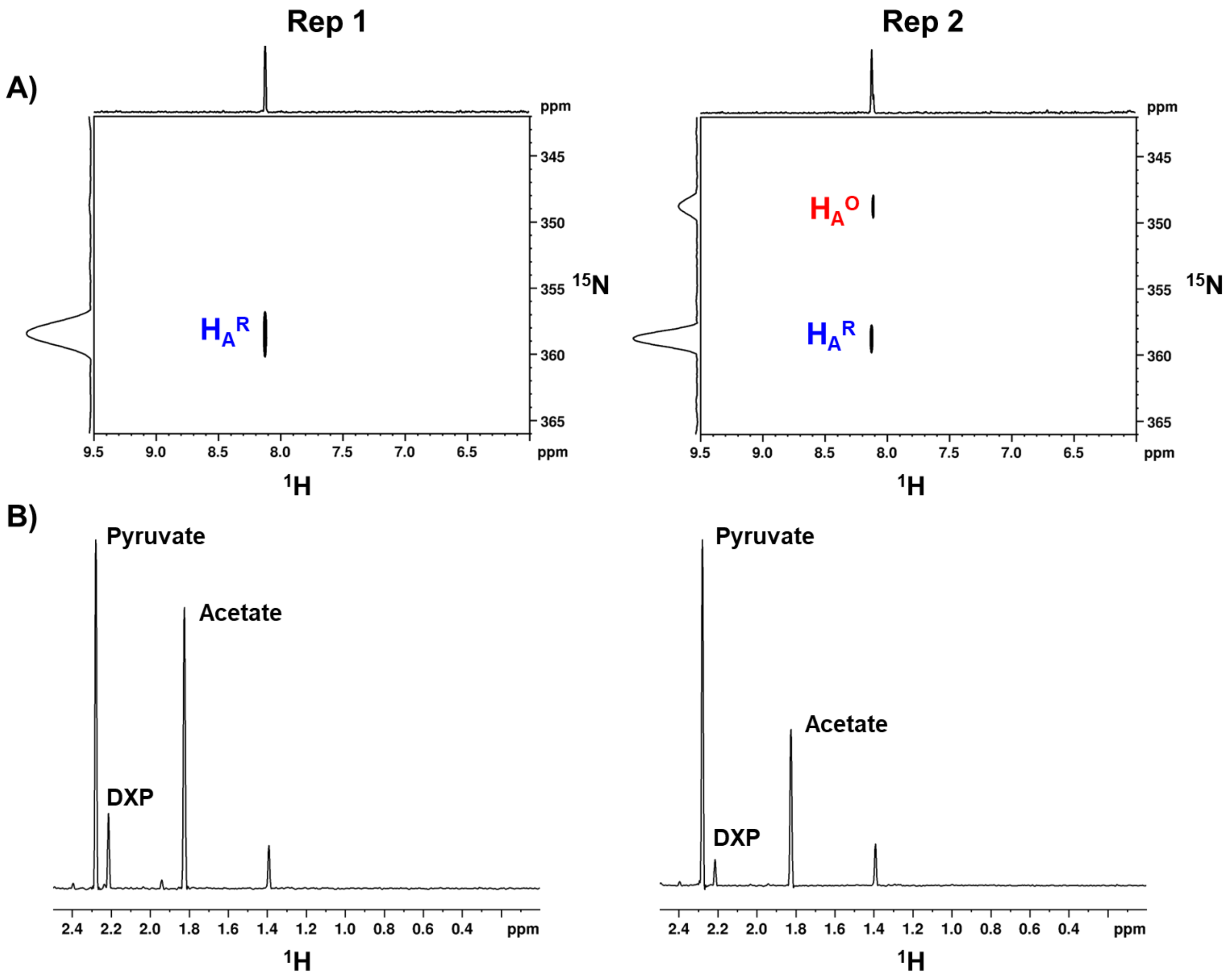


**Figure S22. Long Range $^{1}$H-$^{15}$N HSQC NMR of 8 incubated with DXPS, pyruvate and D-GAP**. Oxime **8** (500 µM) was incubated (as described in the Materials and Methods section) with DXPS (1 µM), pyruvate (2 mM) and D-GAP (500 µM) to detect reduction of **8**. Reduction of **8** (**A**) was observed via LR $^{1}$H-$^{15}$N HSQC concurrent with acetate and DXP formation in the 1D H{$^{13}$C} and $H_3${$^{13}$C} filtered experiment (**B**).

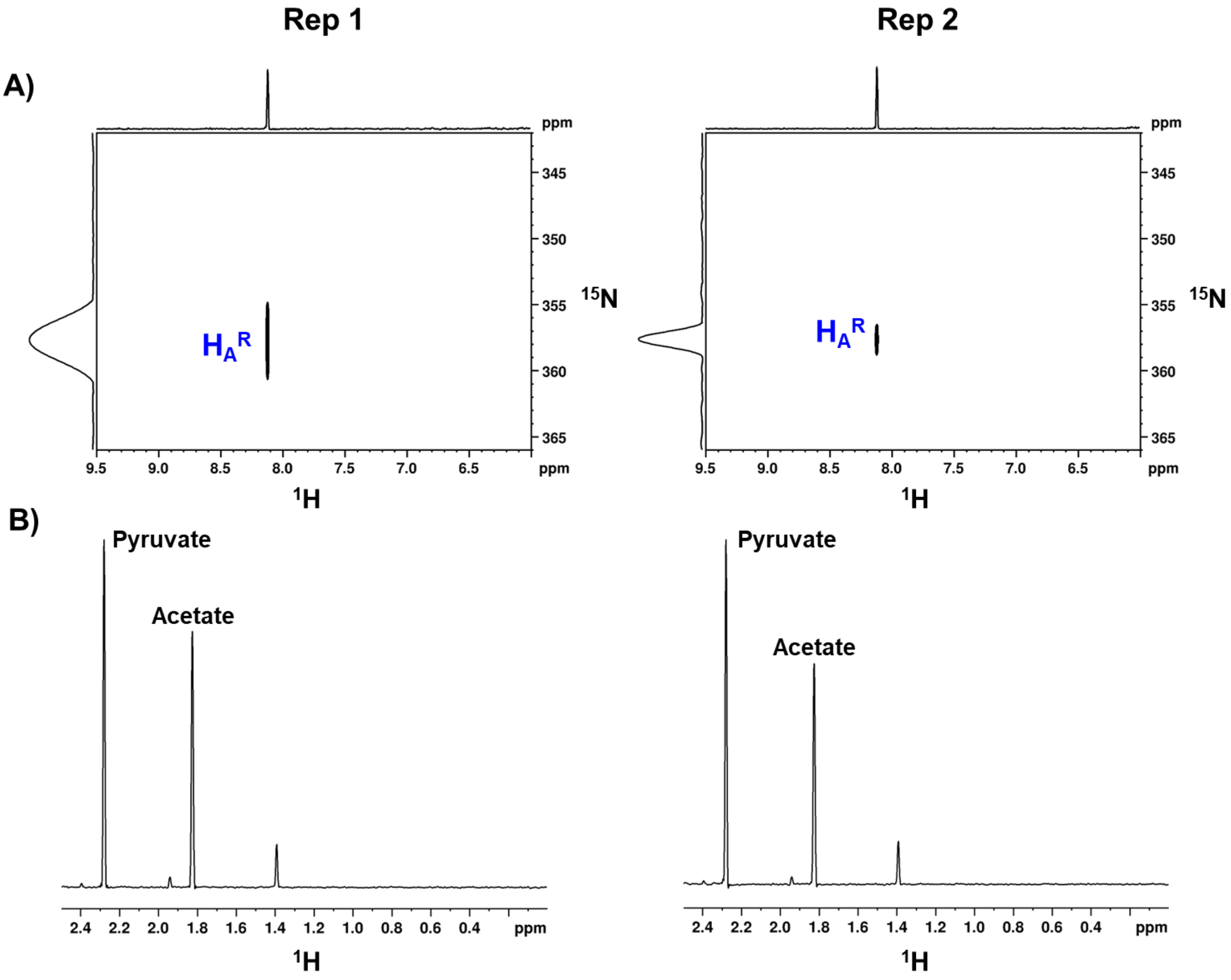


**Figure S23. Long Range $^1$H-$^{15}$N HSQC NMR of 8 incubated with DXPS and pyruvate**. Oxime **8** (500 µM) was incubated (as described in the Materials and Methods section) with DXPS (1 µM) and pyruvate (2 mM) to detect reduction of **8**. Reduction of **8** (**A**) was observed via LR $^1$H-$^{15}$N HSQC concurrent with acetate formation in the 1D H{$^{13}$C} and $H_3${$^{13}$C}filtered experiment (**B**).

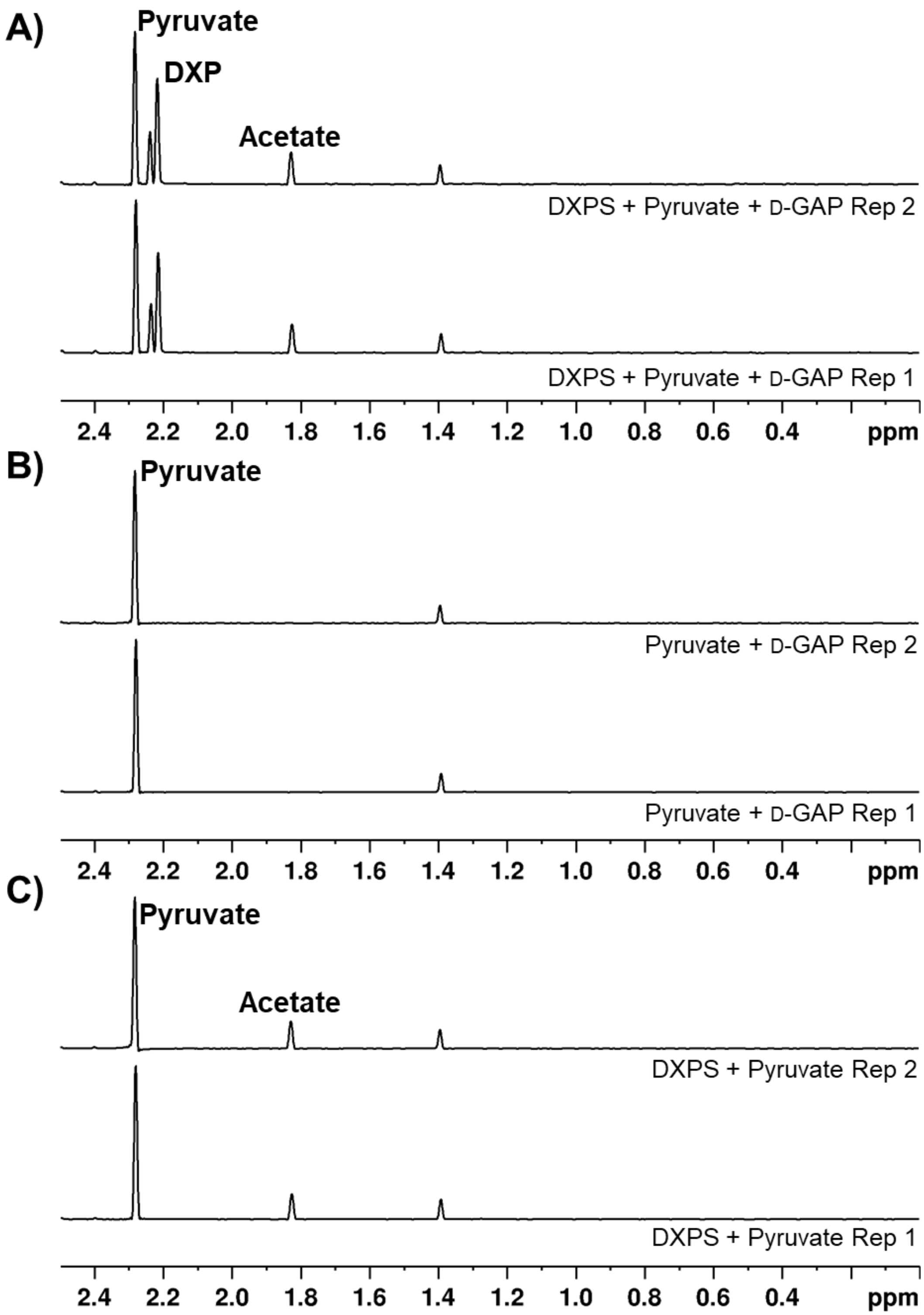


**Figure S24. NMR detection of DXPS reaction products**. Reactions with DXPS (1 µM), pyruvate (2 mM) and D-GAP (500 µM) were conducted in the absence of **8** as described in the LR $^{1}H$-$^{15}N$ HSQC experiments (S19-23, Materials and Methods) as controls to detect product formation via the 1D H{$^{13}C$} and $H_3${$^{13}C$} filtered experiment. **A**) Reactions containing DXPS, pyruvate, and D-GAP produced larger quantities of DXP relative to acetate. **B**) A mixture of pyruvate and D-GAP was stable under these conditions. **C**) A small amount of acetate was formed in the presence of DXPS and pyruvate.

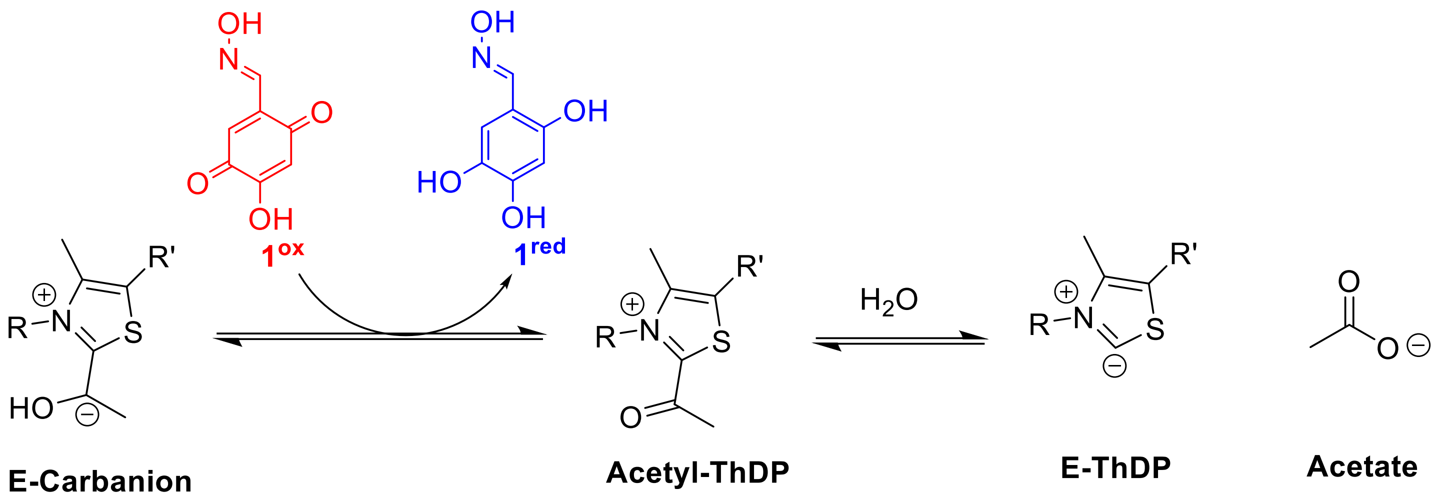


**Figure S25. Mechanism of acetate formation from reduction of 1**. After decarboxylation of LThDP, **1** is reduced by subsequent E-Carbanion, forming acetyl-ThDP which undergoes hydrolysis to form acetate.

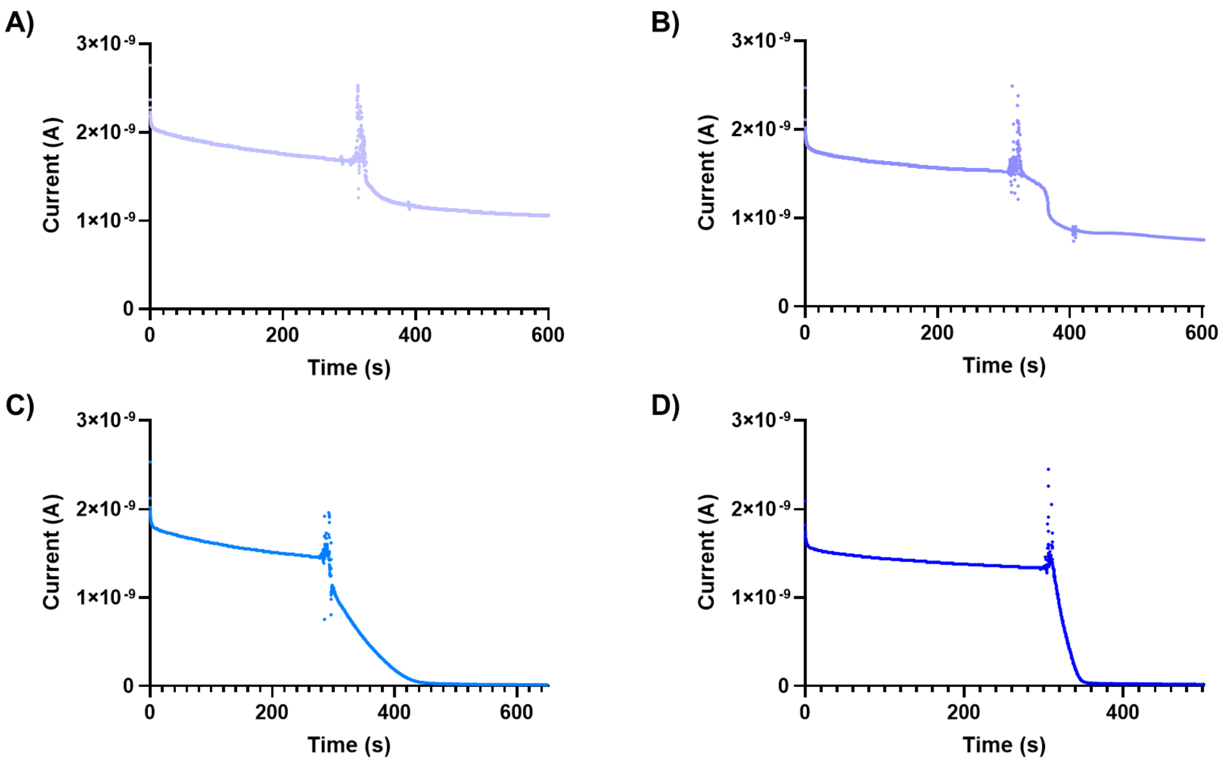


**Figure S26. Raw amperoteric i-t measurements of reactions with 1**. Raw traces of amperoteric i-t measurements of reactions containing **1** (1 mM) and **A)** DXPS (10 µM), **B)** pyruvate (2 mM) + D-GAP (500 µM), **C)** DXPS (10 µM) + pyruvate (2 mM) + D-GAP (500 µM), and D) DXPS (10 µM) + pyruvate (2 mM).

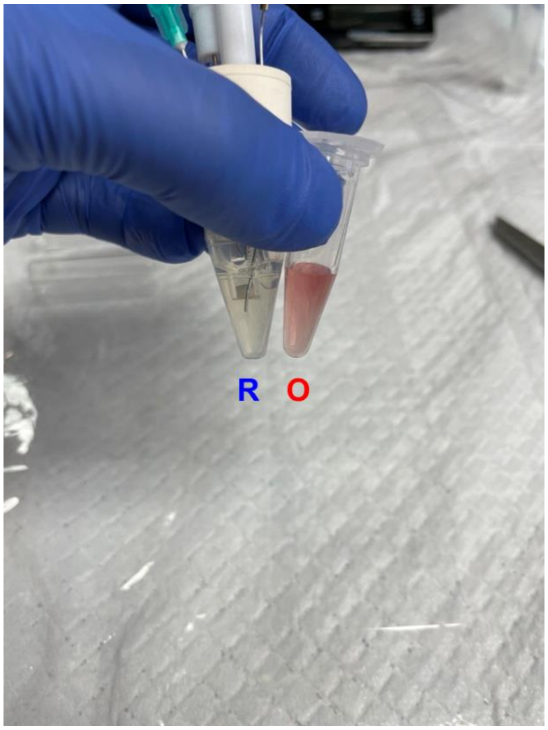


**Figure S27. Color change of oxime 1 based on redox form.** Oxime **1** displays distinct color differences depending on redox state. When oxidized (right), oxime **1** turned the solution pink, and when reduced (left), oxime **1** turned the solution yellow.

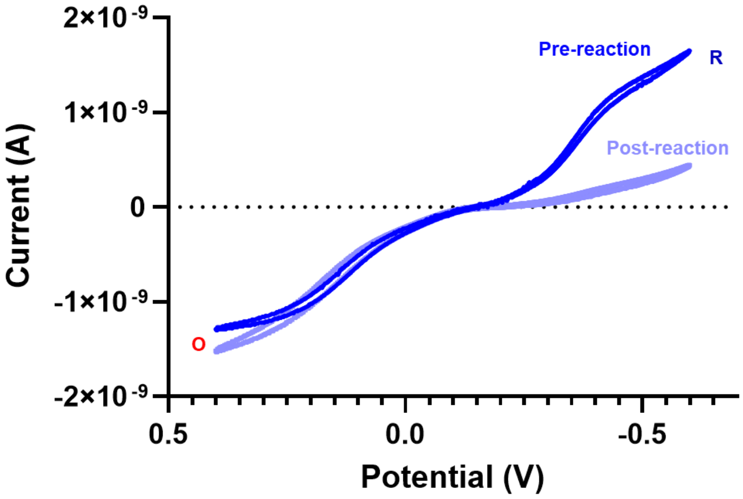


**Figure S28. Full CVs of 8 measured on gold UME**. CV of **8** before and after reaction with DXPS (10 µM), pyruvate (2 mM), and d-GAP (500 µM). A dramatic decrease in measured electrochemical reduction is seen with no corresponding change in oxidation, suggesting they are not coupled.

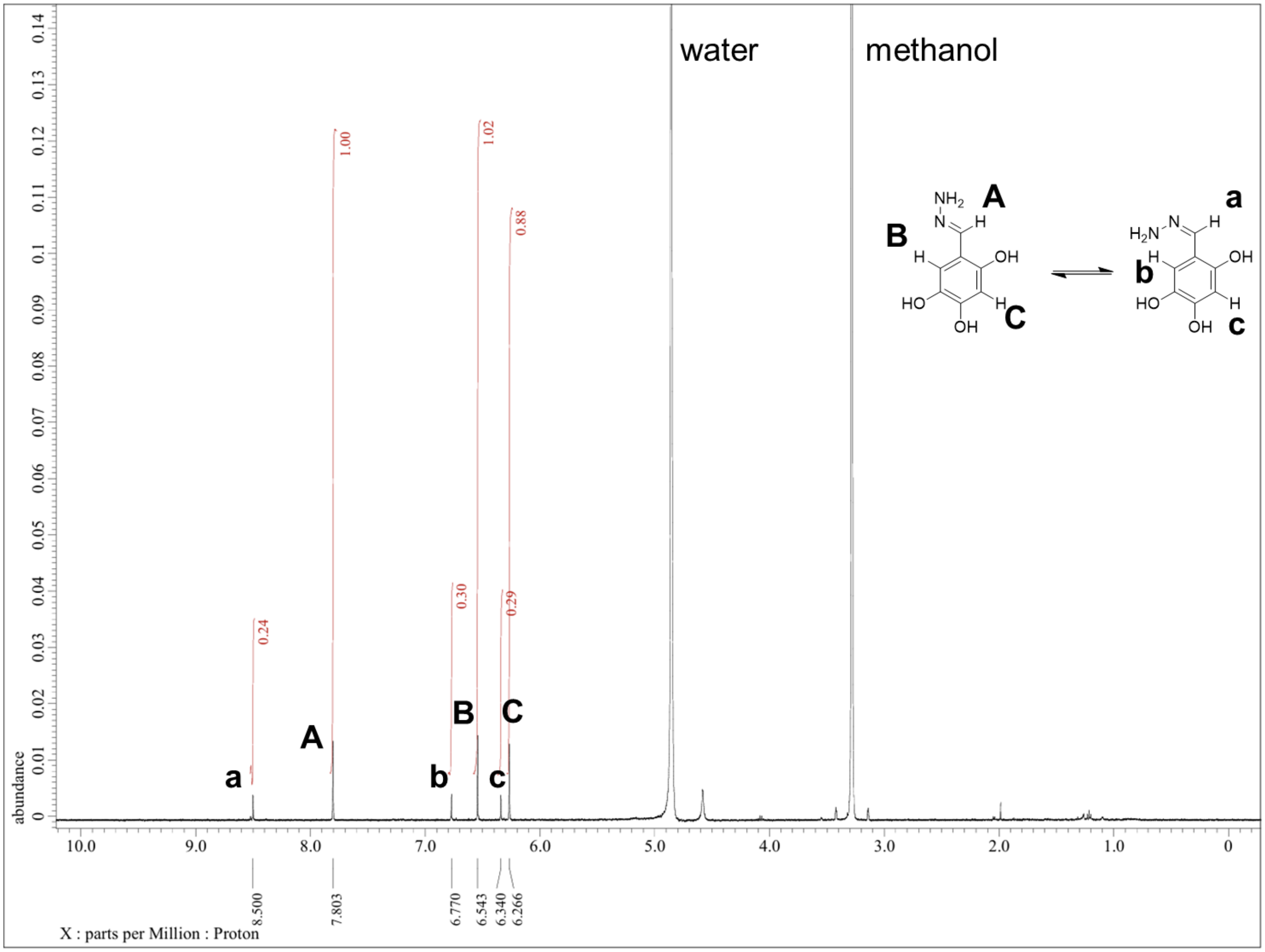


**Figure S29.** $^{1}$**H-NMR of 3**

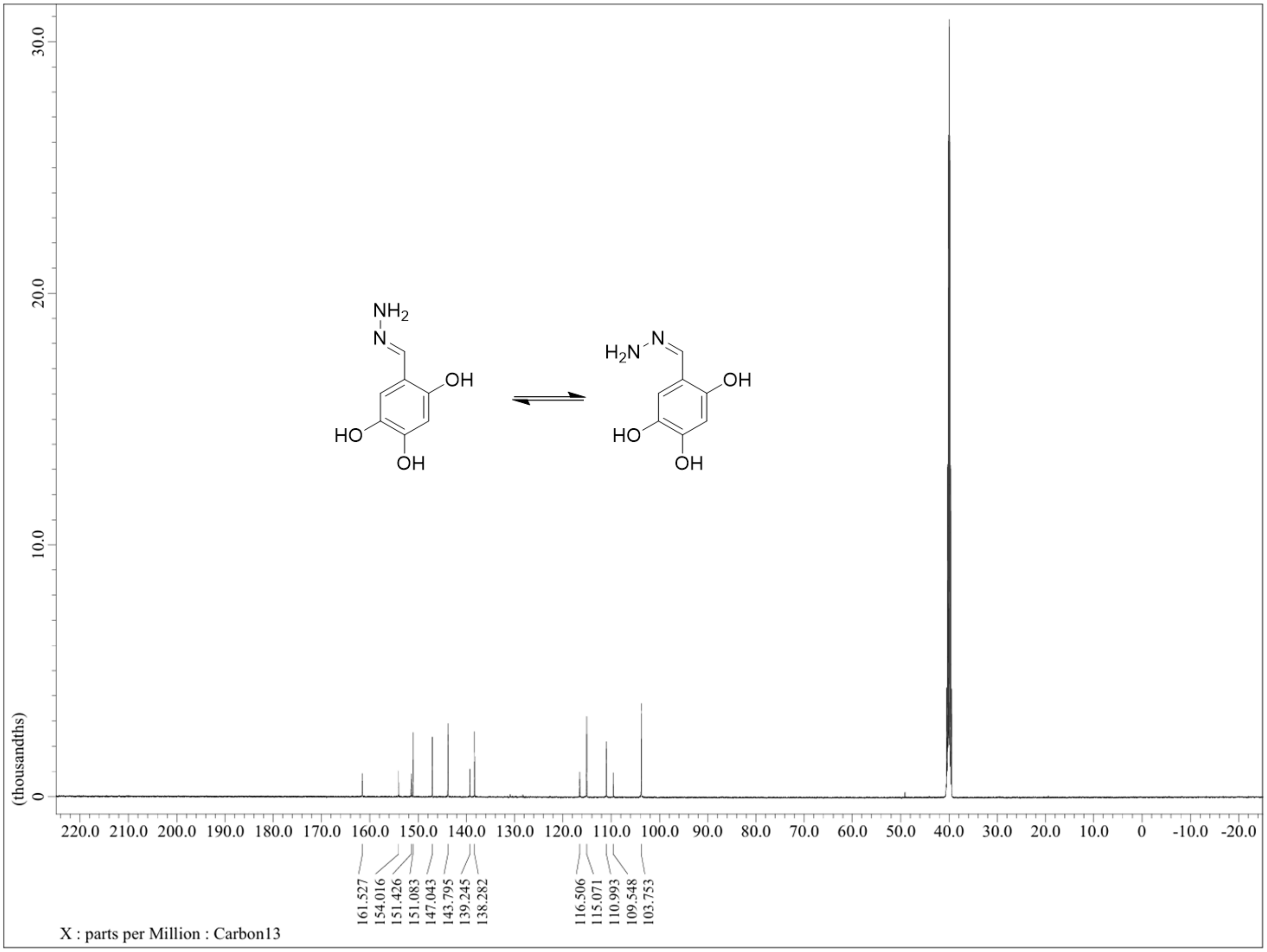


**Figure S30. $^{13}$C-NMR of 3**

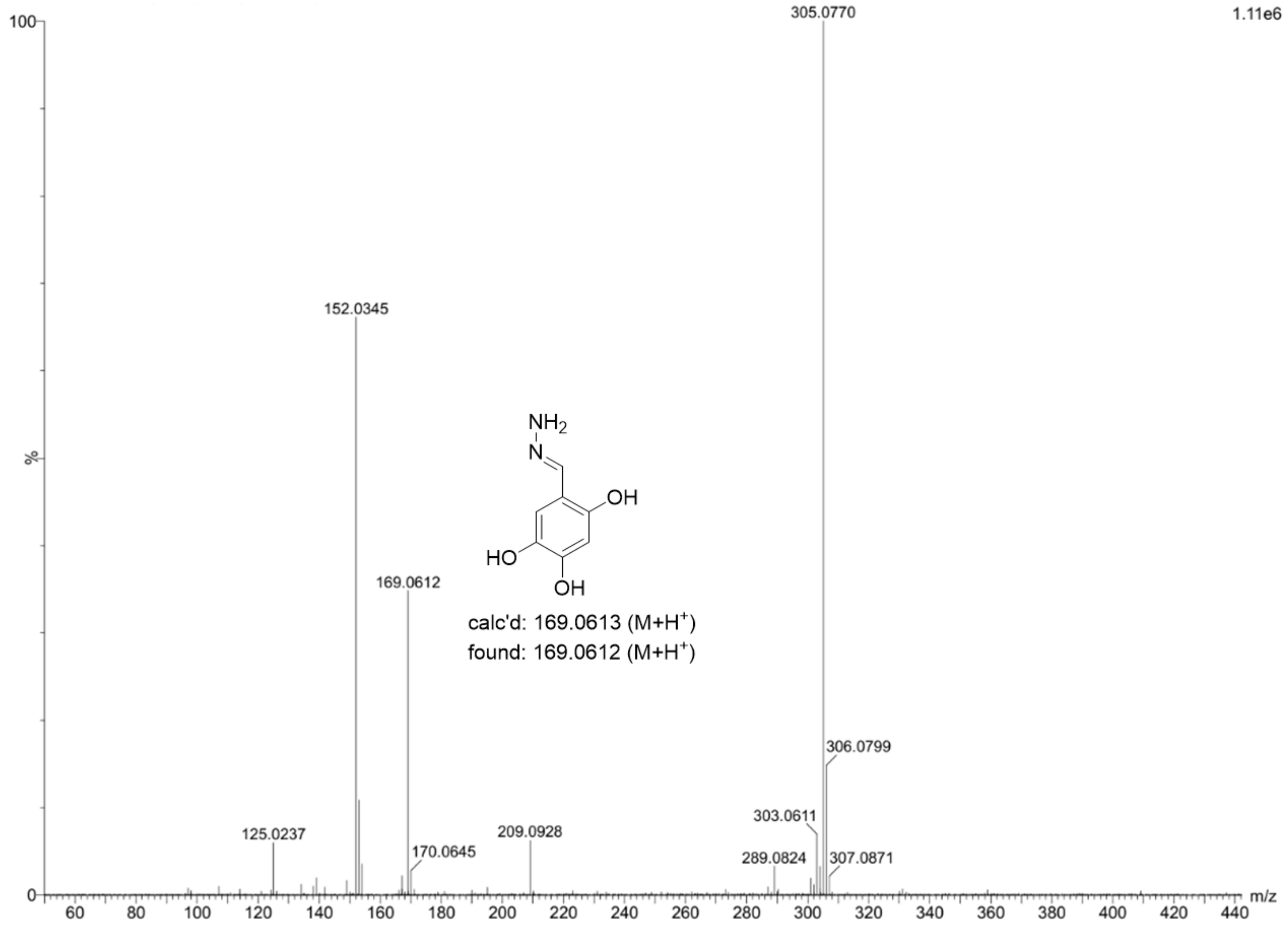


**Figure S31. HRMS detection of $3^R$**

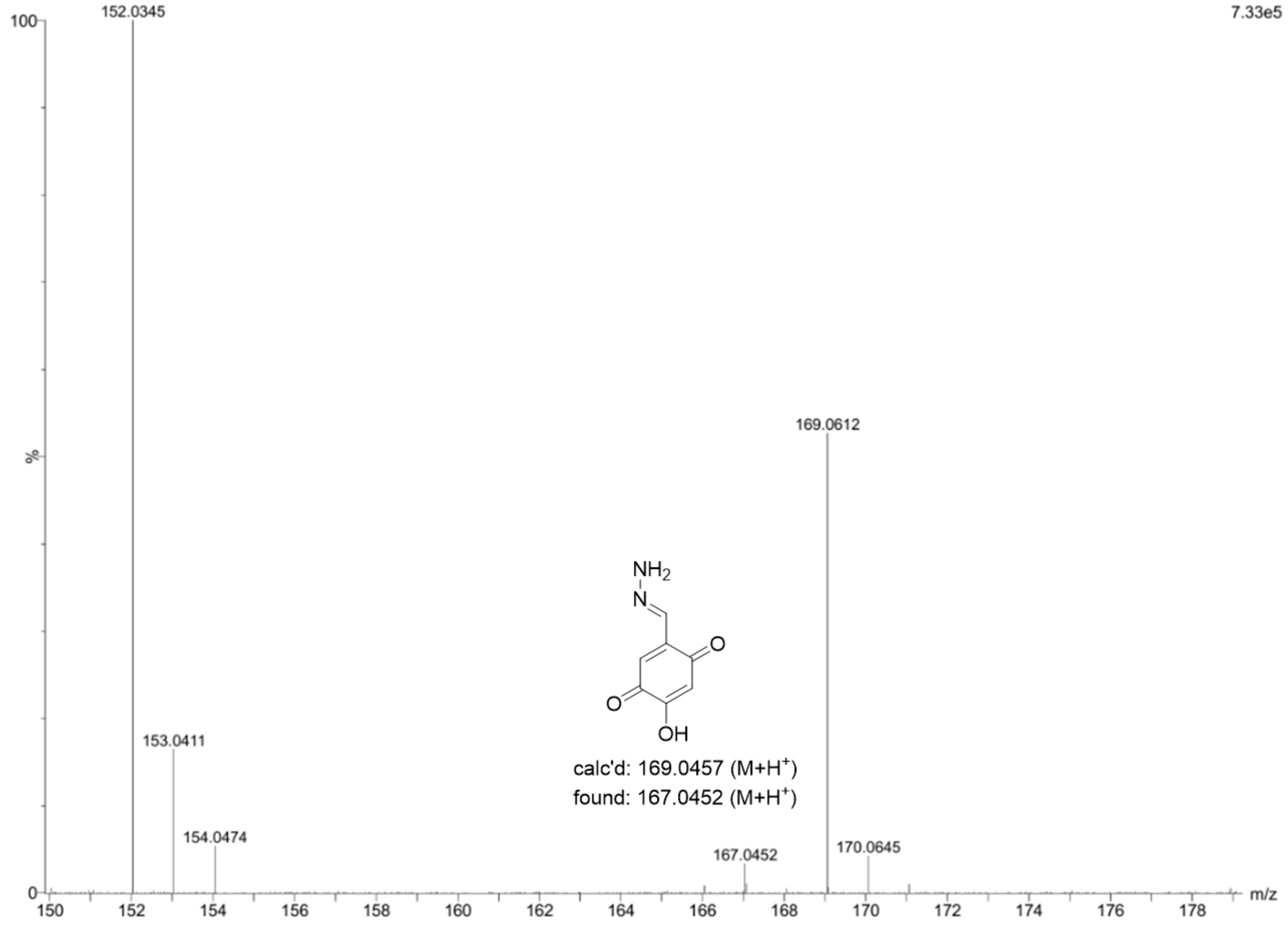


**Figure S32. HRMS detection of 3^O**

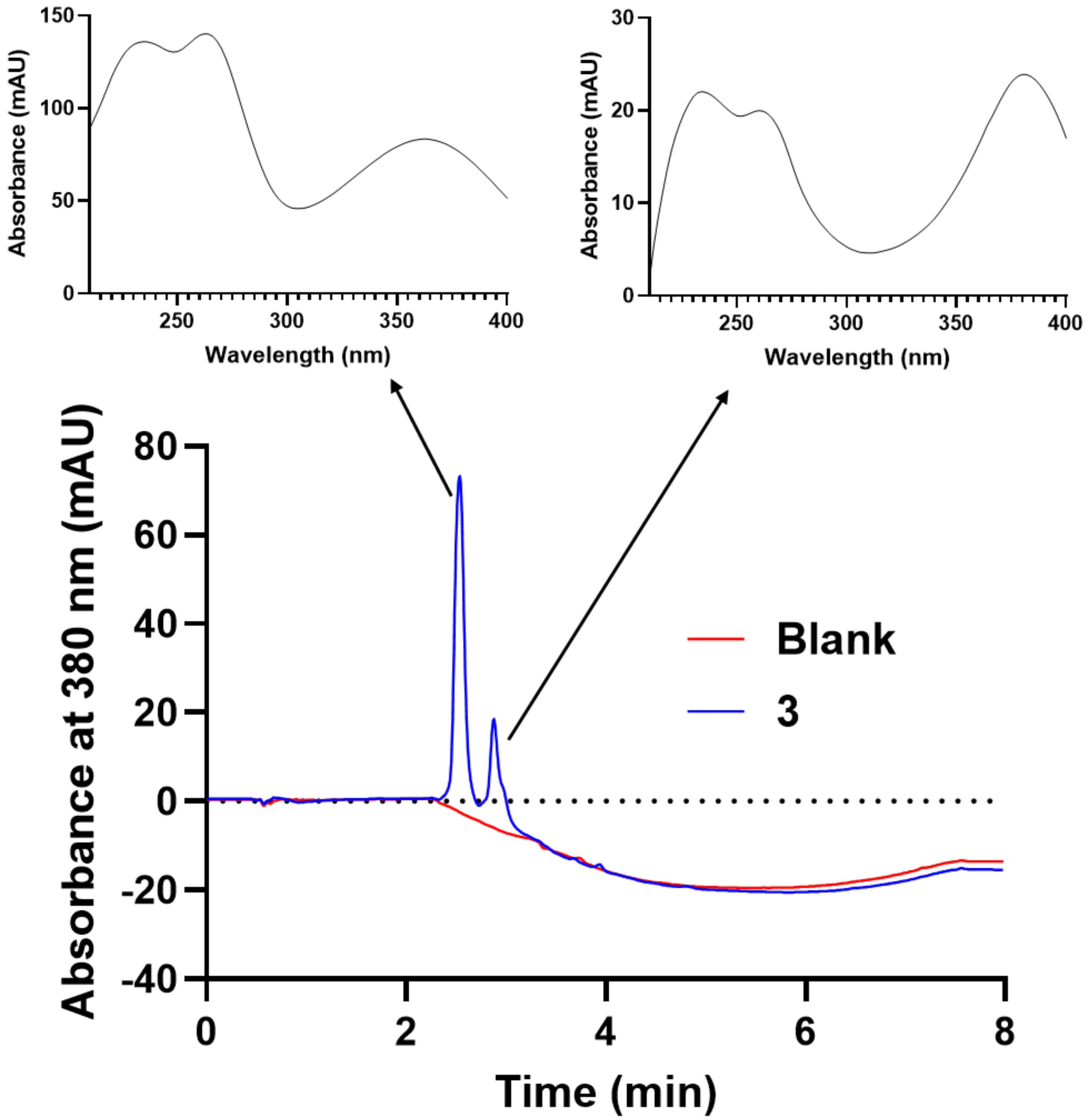


**Figure S33. HPLC of 3**

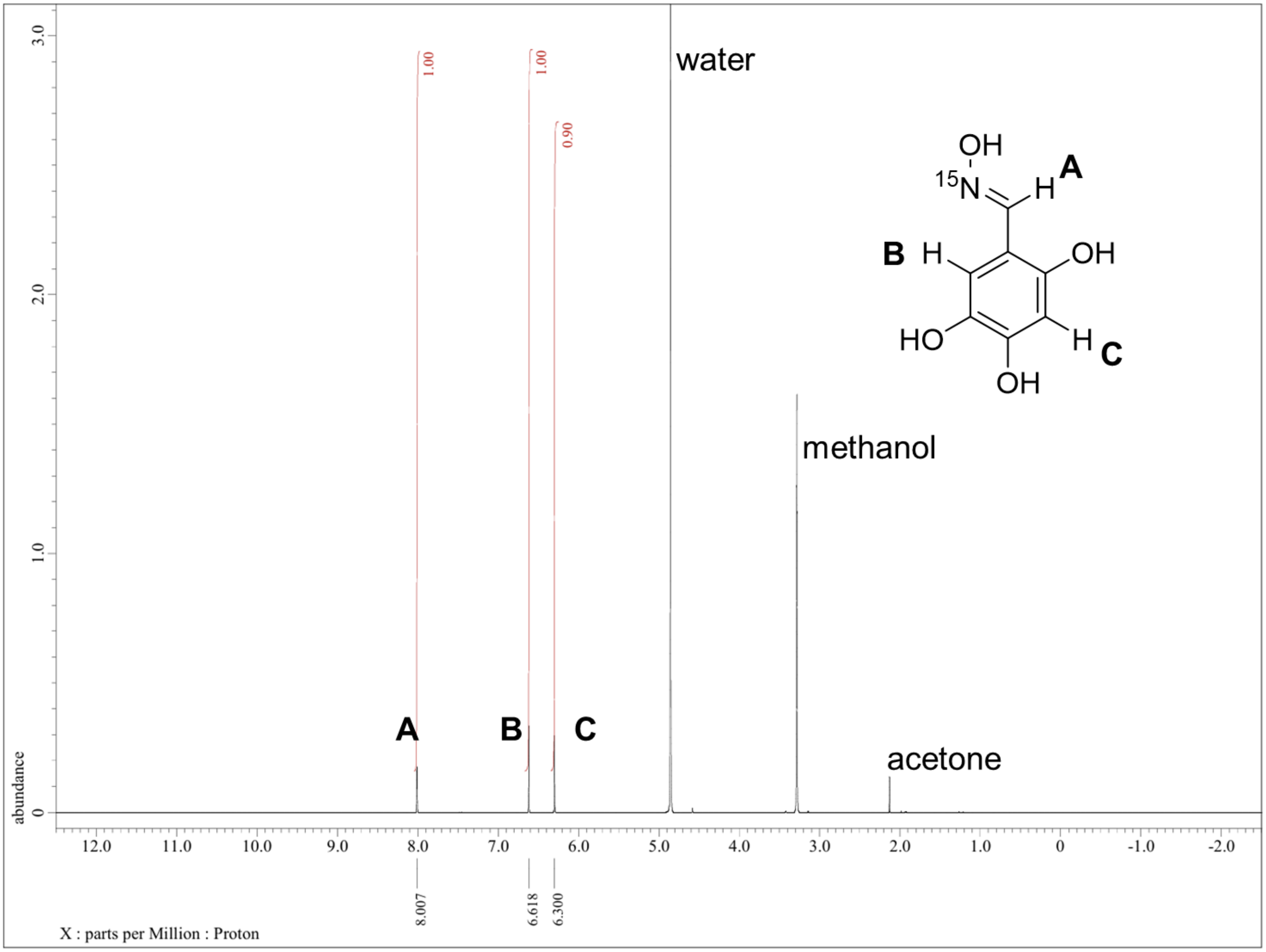


**Figure S34. [1]H-NMR of 8**

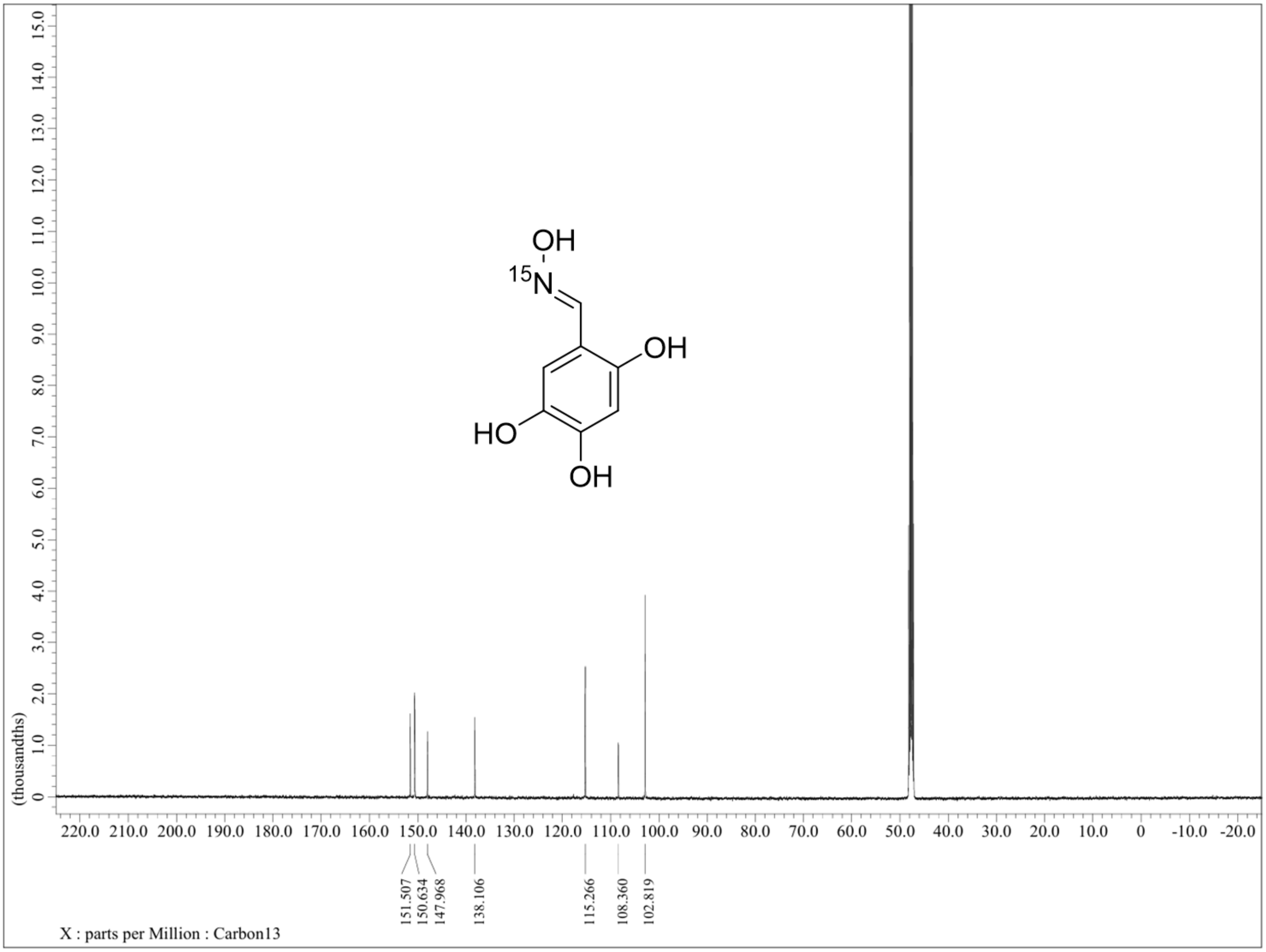


**Figure S35. $^{13}$C-NMR of 8**

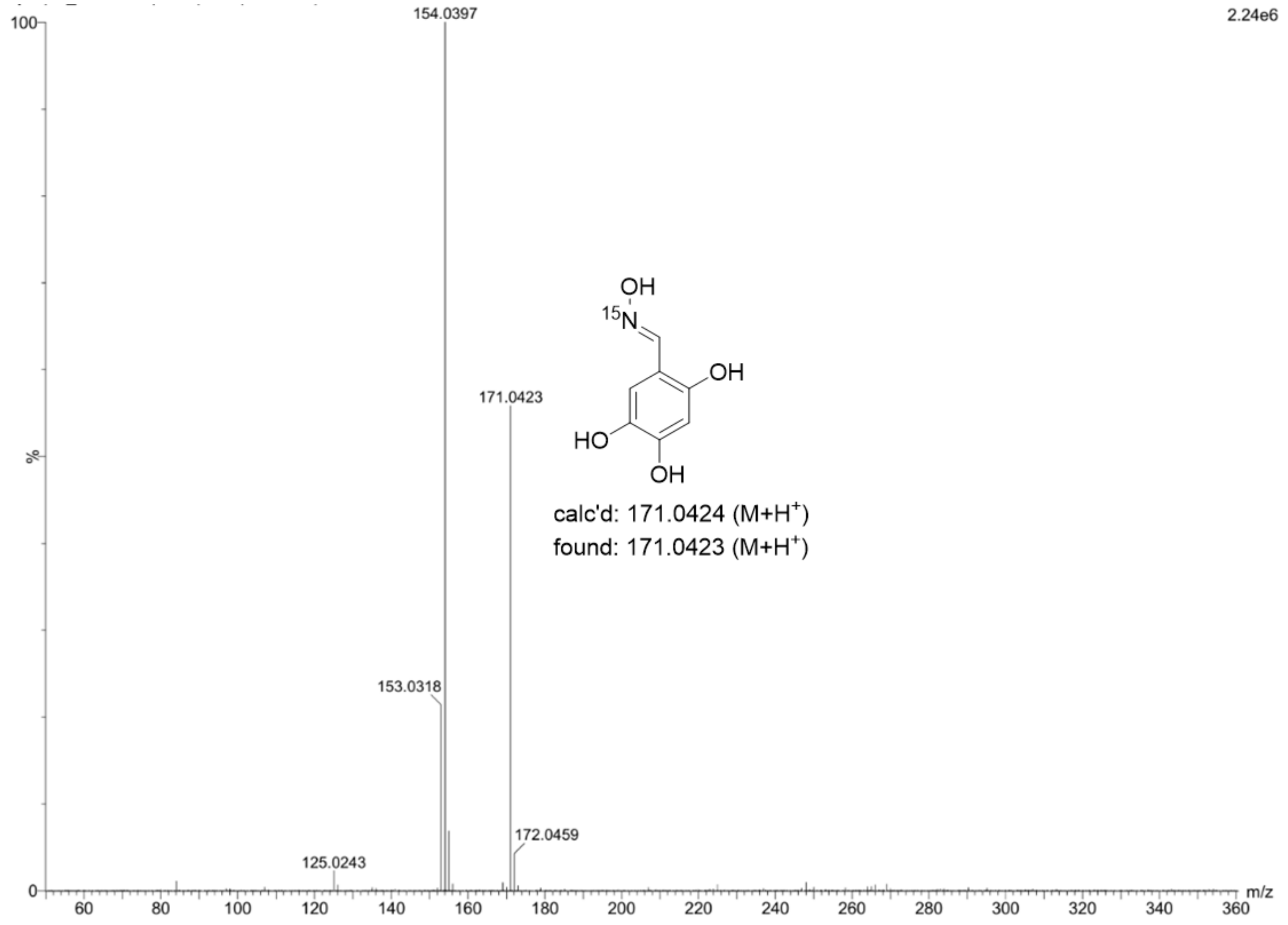


**Figure S36. HRMS detection of 8[R]**

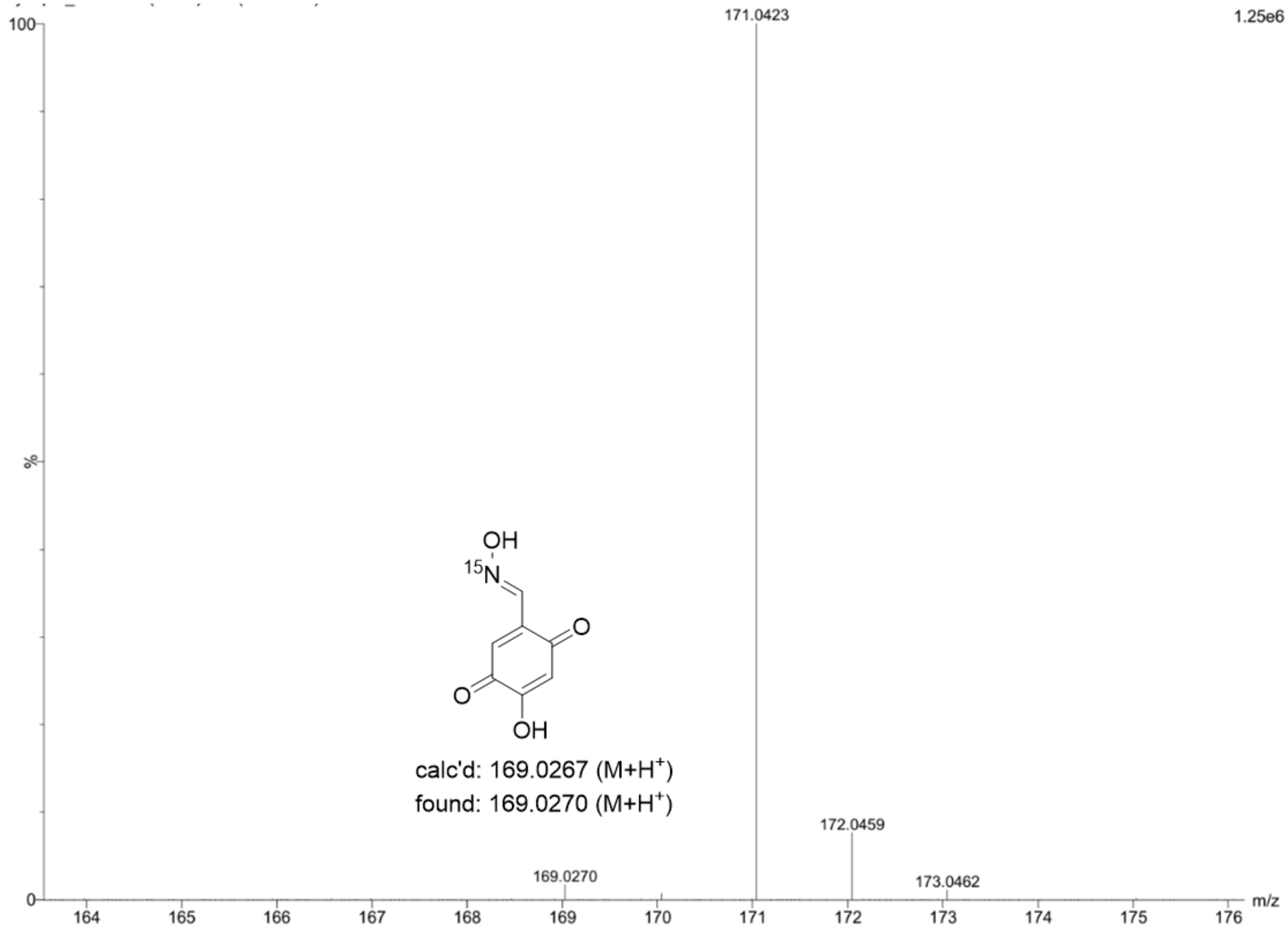


**Figure S37. HRMS detection of 8<sup>O</sup>**

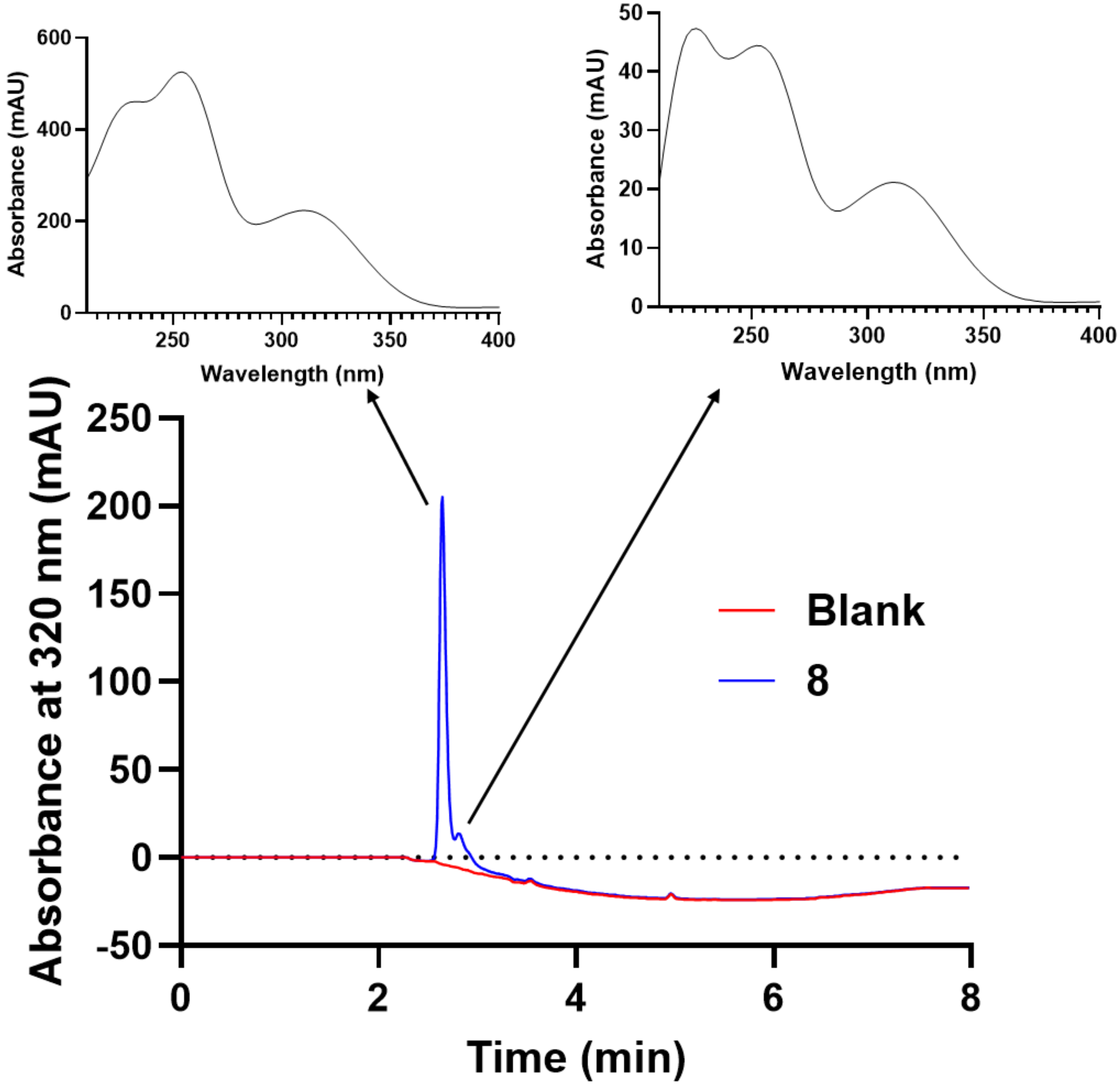


**Figure S38. HPLC of 8.** Shoulder peak is predicted to be **8** in a different oxidation state from the primary peak.

| Enzyme | $K_i$ Vary Pyruvate (μM) | $K_i/K_i^{WT}$ | $K_i$ Vary D-GAP (μM) | $K_i/K_i^{WT}$ |
|---|---|---|---|---|
| WT *Ec*DXPS | 4.7 ± 0.5 | 1 | 2.3 ± 0.1 | 1 |
| WT *Pa*DXPS | 16.4 ± 2.2 | 3.5 | 6.5 ± 1.5 | 2.8 |
| WT *Dr*DXPS | 2.8 ± 0.2 | 0.6 | 3.3 ± 0.2 | 1.4 |
| WT *Ec*DXPS ($-O_2$) | 51.0 ± 6.6 | 10.9 | 10.9 ± 3.3 | 4.7 |
| *Ec*D427A | 4.8 ± 0.24 | 1.0 | ND | ND |
| *Ec*H431A | 1.1 ± 0.1 | 0.2 | ND | ND |
| *Ec*R99A | 30 ± 1 | 6.5 | 8.7 ± 1.2 | 3.8 |

**Table S1. Summary of apparent *K*ᵢs of 1**.

| Compound | $K_i$ Vary Pyruvate (μM) | MOI | $K_i$ Vary D-GAP (μM) | MOI |
|---|---|---|---|---|
| 1 | 4.7 ± 0.5 | Uncompetitive | 2.3 ± 0.1 | Competitive |
| 2 | 15.6 ± 0.7[a] | Noncompetitive[a] | 3.92 ± 0.58[a] | Competitive[a] |
| 3 | 17.5 ± 0.6 | Uncompetitive | 12.4 ± 1.9 | Mixed |
| 4 | 56 ± 11 | Uncompetitive | 23.8 ± 0.6 | Mixed |

**Table S2. Summary of apparent $K_i$s of 1, 2, 3, and 4**.
[a]Values for **2** reported by Bartee et al.[1]

### References

(1) Bartee, D.; Morris, F.; Al-Khouja, A.; Freel Meyers, C. L. Hydroxybenzaldoximes Are D-GAP-Competitive Inhibitors of E. Coli 1-Deoxy-D-Xylulose-5-Phosphate Synthase. *Chembiochem* **2015**, *16* (12), 1771–1781.
